# Identification and characterization of host-directed therapeutics for tuberculosis using a versatile human 3D tuberculoma bioplatform

**DOI:** 10.1101/2025.08.27.672670

**Authors:** Suraj B. Sable, Wen Li, Allison Kline, Sakthivel Govindaraj, Vijayakumar Velu, James E. Posey

## Abstract

Host-directed therapies (HDT) represent a pivotal strategy in combating both drug-susceptible and drug-resistant tuberculosis (TB). Current evaluations, however, show limited success in animal models and clinical settings, emphasizing the need for more effective HDT candidates. With few druggable targets validated within granulomas, it is essential to verify the effectiveness of HDT candidates identified through traditional macrophage cultures in the context of the tuberculous granuloma milieus. Bioengineering a scalable, high-throughput screening (HTS) platform that replicates the physiological microenvironments in the hallmark tubercular lesions could significantly improve the identification of relevant HDT candidates and new treatment strategies. Here, we developed a facile, HTS-compatible bioplatform that generates tuberculoma-emulating structures, following three-dimensional (3D) co-cultures of human cells and pathogenic mycobacteria. Employing high-content imaging alongside immunological and transcriptomic approaches, we demonstrated that these 3D structures exhibit classic tuberculoma attributes and develop crucial transformations. Utilizing this system, we screened antibody biosimilars and potential HDT compounds. Our findings demonstrate the system’s versatility in discovering antimicrobials and HDT candidates that effectively reduce mycobacterial burdens and granuloma lesions, while elucidating their immune mechanisms within 3D milieus. Many compounds effective in two-dimensional (2D) cultures were ineffective once granulomas formed in our 3D model. Notably, several promising compounds were found to induce rapid autophagy flux, and we validated the effectiveness of one such compound, the multi-kinase inhibitor AT9283, in a mouse model. Our findings highlight several HDT candidates for potential repurposing in TB treatment, offering a robust tool for accelerating therapeutic discoveries and advancing translational research for TB and other granulomatous diseases.

**Summary:** Safer and shorter treatment regimens for tuberculosis (TB) are urgently needed. Host-directed therapies (HDT) are being explored to enhance antibiotic regimens and address antimicrobial resistance. To expedite the discovery of HDT candidates and the development of new treatment strategies, scalable *in vitro* systems are needed that can replicate the critical features and microenvironments of TB lesions in a format compatible with high-throughput screening (HTS). We developed an HTS-compatible bioplatform using 3D co-cultures of human cells and fluorescent mycobacteria, creating tuberculoma-like structures with classic solid, necrotic, and cavitary transformations that exhibit crucial microenvironments. This facile system enabled us to screen antibody biosimilars and HDT compounds in 3D human *in vitro* tuberculomas, identifying candidates that inhibited mycobacterial growth and granuloma lesions while revealing potential innate immune mechanisms. The study revealed several promising HDT candidates that could be repurposed for TB treatment, introducing a versatile tool for screening therapeutic libraries. Additionally, it presents a framework for enhancing human *in-vitro* granuloma models, as advances in tissue-like systems emerge to recapitulate the architecture and multilineage differentiation in the lungs.

## Introduction

Tuberculosis (TB) remains the leading cause of death from a single infectious agent worldwide, claiming approximately 1.25 million lives annually [1]. Current antibiotic treatments for TB are lengthy, often toxic, and last between 4 and 24 months, depending on the regimen and drug susceptibility [2, 3]. This pathogen-targeted approach can lead to microbial resistance, with ∼400,000 cases of multidrug- or rifampicin-resistant TB emerging each year [1]. Despite advances in antibiotics and treatment regimens, *Mycobacterium tuberculosis* (*Mtb*) can quickly develop resistance, complicating disease management [2–4]. Even after successful treatment, individuals often suffer from post-TB lung disease due to persistent inflammation and tissue damage [5]. These challenges underscore the urgent need for safer, shorter, and more effective therapeutic strategies [6].

Host-directed therapies (HDT) are being explored to improve TB treatment by modulating the host immune response rather than directly targeting the pathogen [7–9]. This approach aims to shorten treatment duration and mitigate microbial resistance by enhancing immunity and targeting cellular processes that *Mtb* exploits for survival. Additionally, it seeks to alleviate immunopathology and prevent disease relapse [7–9]. Progress has been made in preclinical evaluations, with candidates, including *N*-acetylcysteine, metformin, statins, vitamin D, and imatinib, entering clinical trials [9, 10]. However, substantial translational advancements are hindered by challenges in identifying key druggable host pathways and predicting therapeutic efficacy within the pathological lesions. Granuloma formation is the hallmark pathological feature of TB, where macrophage-rich aggregates develop to control the infection, but can also support *Mtb* and cause lung damage [11–13]. These granulomas limit drug penetration and immune access, facilitating *Mtb* persistence and drug resistance [14, 15]. The granulomas can consolidate into three-dimensional (3D) structures called ’tuberculomas’ [16–18], often with hypoxic and necrotic cores surrounded by a florid granulomatous response featuring macrophages in various stages of transformation and a lymphocyte-rich cuff. Tuberculomas may cavitate and facilitate *Mtb* transmission, leading to poor treatment outcomes and increased risk of drug resistance [19].

Animal models remain essential for preclinical studies of potential therapeutics [3, 20]. However, their low throughput and high costs restrict large-scale drug discovery. Traditional culture methods often fail to replicate microenvironments within granulomas [21], which has prompted the development of several two-dimensional (2D) and 3D *in-vitro* granuloma models using human cells [22–29]. These models have provided new insights into host-*Mtb* interactions [30], and some have enabled high-content screening (HCS) of select antibiotics [31]. While contemporary 3D human *in-vitro* granuloma models have improved physiological relevance compared to 2D systems [26, 27], advancements are needed regarding diverse granuloma phenotypes and compatibility with high-throughput screening (HTS) [30]. Models utilizing primary cells isolated from biopsies, leukapheresis, or bronchoalveolar lavage are impractical for extensive compound testing. Other 3D systems, such as lung-on-chip and human pluripotent stem cell (hPSC)-derived organoid models, have advanced our understanding of lung epithelial cell-*Mtb* interactions [32, 33]; however, they lacked granuloma formation. Micro-dissected tuberculous granulomas from infected animals, as explant 3D cultures, offer potential but are limited in throughput [34].

A simplified 3D model, analogous to classical human tuberculomas, is needed for creating a scalable, HTS platform that enables rapid, efficient, and cost-effective testing of extensive therapeutic libraries. Additionally, the identification and characterization of candidates that reduce both bacterial burdens and lesions are crucial, given the limited evidence on the effectiveness of HDT as a standalone treatment in infected hosts after granuloma onset [35]. Furthermore, it is necessary to confirm the efficacy of candidates with known mechanisms for treating non-infectious diseases, as identified through 2D systems, within the 3D tuberculoma environment.

To address this, we developed an HTS-compatible physiological system utilizing a 3D co-culture of human primary cells or immortalized monocytes with virulent mycobacteria (**Fig. 1**). This system generates 3D structures that replicate critical traits in classic solid, necrotic, or cavitary tuberculomas, within a 96-well format suitable for immunostaining, fluorescence reading, high-content imaging, and drug screening. In proof-of-concept experiments, we screened a custom HDT compound library and identified several compounds that significantly reduced bacterial burden and resolved lesions. Many identified ‘hit’ compounds promoted autophagy, highlighting the potential to characterize mechanisms of action and identify host pathways amenable to pharmacological intervention. One such compound, a clinically evaluated potent aurora kinase inhibitor, AT9283 [36, 37], exhibited substantial efficacy in the murine model. Our study advances 3D human *in-vitro* tuberculoma technology for HTS and identifies several active HDT candidates with potential for TB therapy.

**Fig. 1.**
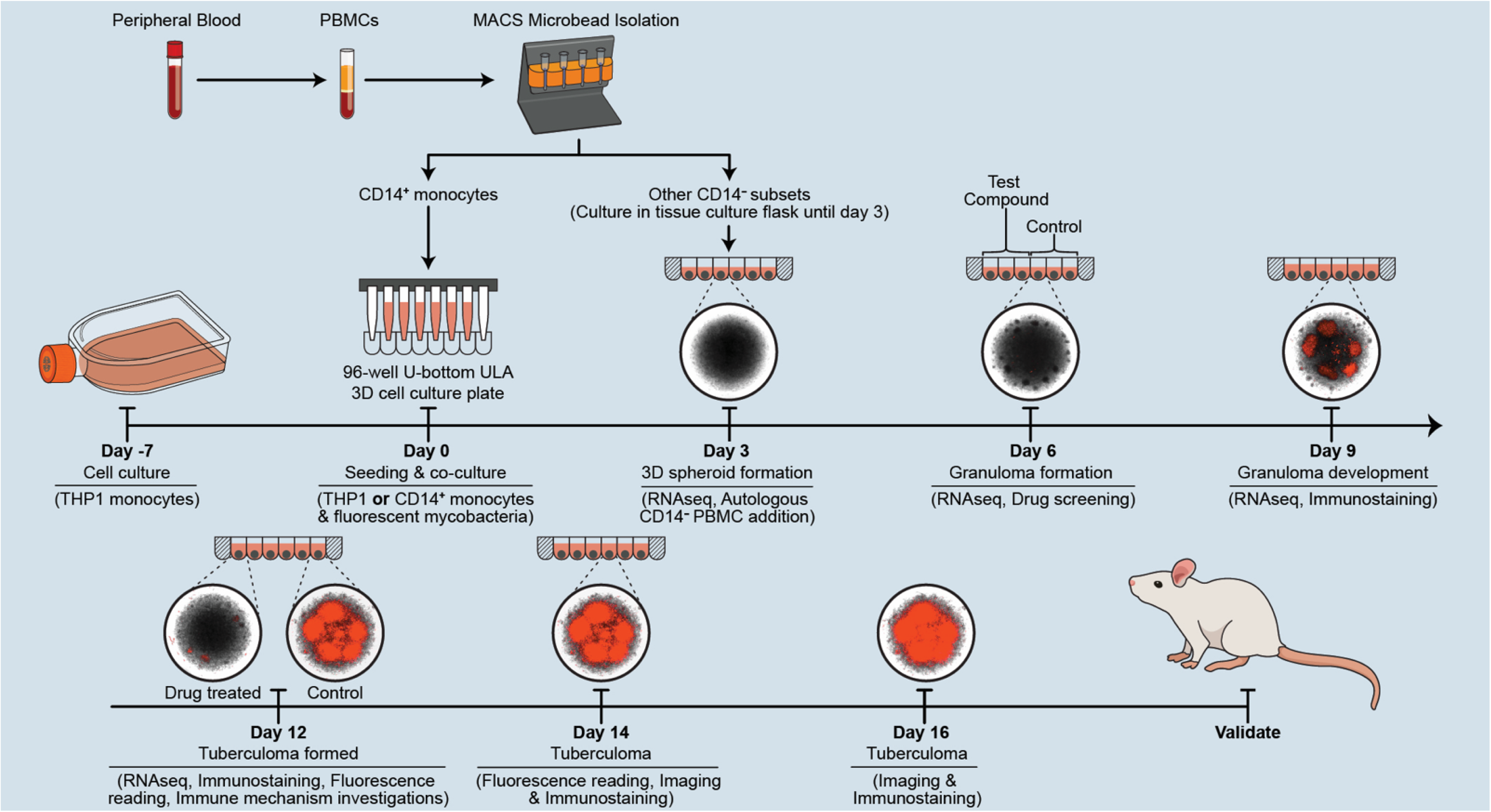
Schematic overview of the ‘mycobacteria-in-spheroid’ co-culture workflow, the resultant 3D tuberculoma-like model, and the bioplatform. Human CD14^+^ monocytes purified from the peripheral blood mononuclear cells (PBMCs) using magnetic-activated cell sorting (MACS) or immortalized THP-1 monocytes are co-cultured with fluorescent pathogenic mycobacteria in the rationally selected 3D cell culture microplates. On day 3, co-cultures of purified CD14^+^ monocytes are supplemented with autologous CD14^−^ PBMC subsets. Alternatively, mycobacteria-in-spheroid co-cultures are generated using reporter monocytes and mycobacteria to investigate microenvironments and immune responses. 3D tuberculoma-like structures generated with conglomerating granulomas are used for fluorescence-based imaging and investigating host-pathogen interactions. The resultant bioplatform is utilized for the rapid screening of biologics and chemical compounds to assess drug efficacy and host-directed immune mechanisms *in situ*. Subsequently, animal models can validate the identified ‘hit’ candidates.

## Results

### A ‘mycobacteria-in-spheroid’ co-culture establishes a facile human 3D tuberculoma-like model

We aimed to create a simple 3D model that reproduces the microenvironments of classic tubercular lesions in a standard 96-well plate and sought to identify the minimum components and culture conditions needed for a dynamic, HTS-compatible granuloma model that reflects prolonged host-pathogen interactions. We developed self-assembling ‘mycobacteria-in-spheroid’ co-cultures using pathogenic mycobacteria and human macrophages in early iterations. To enhance efficiency, we utilized *Mycobacterium marinum* (*Mm*), a risk group-2 pathogen naturally resistant to several antibiotics and a genetically close relative of *Mtb*, in the BSL-2 laboratory as an *Mtb* surrogate [11–14]. We assessed two strains, 1218 and ‘M’, that constitutively expressed green fluorescent protein (GFP) and tdTomato (red) to track bacterial growth dynamics. Since *Mm* grows well at 30°C in poikilothermic animals, we compared the ability of the two strains to grow at 37 °C *in vitro*. We found the strains grew well at 37°C in Middlebrook 7H9 medium, although relatively slower compared to 30°C, and showed robust and comparable growth kinetics over three weeks (**S1A Fig**). However, unlike the 7H9 growth medium, both strains grew very slowly in the 3D cell culture medium without macrophages.

We tested the growth of two *Mm* strains in 3D cell cultures of several human and mouse monocyte-macrophage cell lines (**S1B** and **S1C Fig**). Such cell lines have been previously used for high-throughput chemical compound, siRNA, and CRISPRi library screens in 2D cell cultures, identifying HDT candidates and druggable targets [38–42]. We used a low multiplicity of infection (MOI) of 0.008 (range, 0.005–0.01) without washing the co-cultures after mixing of cells and bacteria. Since pinocytosed aminoglycosides can reach phagosomes and contribute to macrophages’ antibacterial activity, we avoided using aminoglycoside antibiotics, which are believed to kill only extracellular bacteria. We omitted mitogens, such as phorbol myristate acetate, for macrophage differentiation, as these could alter host-pathogen interactions. An automated multimode plate reader equipped with a cell imager was used for fluorescence monitoring and *in situ* imaging. Human THP-1 monocytes and U937 histiocytes (tissue macrophages) allowed for the most consistent and robust bacterial growth in 3D co-culture (**S1B** and **S1C Fig**). Conversely, mouse macrophage cell lines J774A.1, RAW-264.7, and AMJ2-C11 demonstrated relatively poor growth of *Mm* strains over 12 days. Only THP-1 cultures with mycobacteria formed cellular aggregates resembling organized collections of macrophages that supported mycobacterial growth, whereas no aggregates were observed in uninfected controls.

Next, we investigated the spatiotemporal dynamics of monocyte and mycobacterial proliferation in 3D co-cultures. Real-time serial imaging revealed that by day 2, solitary spheroids were forming in the microwells (**Fig 2A** and **2B**). During this formation, the inoculated bacteria were completely gathered by the amassing monocytes, proliferating primarily within the spheroids. By day 3, compact spheroids formed, increasing in size by day 6. Cellular aggregates developed around days 6 and 8 post-co-culture in the spheroids infected with strain ‘M’ and 1218, respectively. These cellular aggregates grew over time, and large granulomatous aggregates, supporting robust bacterial growth, formed by days 12 and 16 (**Fig 2A** and **2B**). At this stage, the 3D ‘mycobacteria-in-spheroid’ structure resembled a conglomeration of florid granulomatous foci in the solid tuberculoma.

**Fig. 2.**
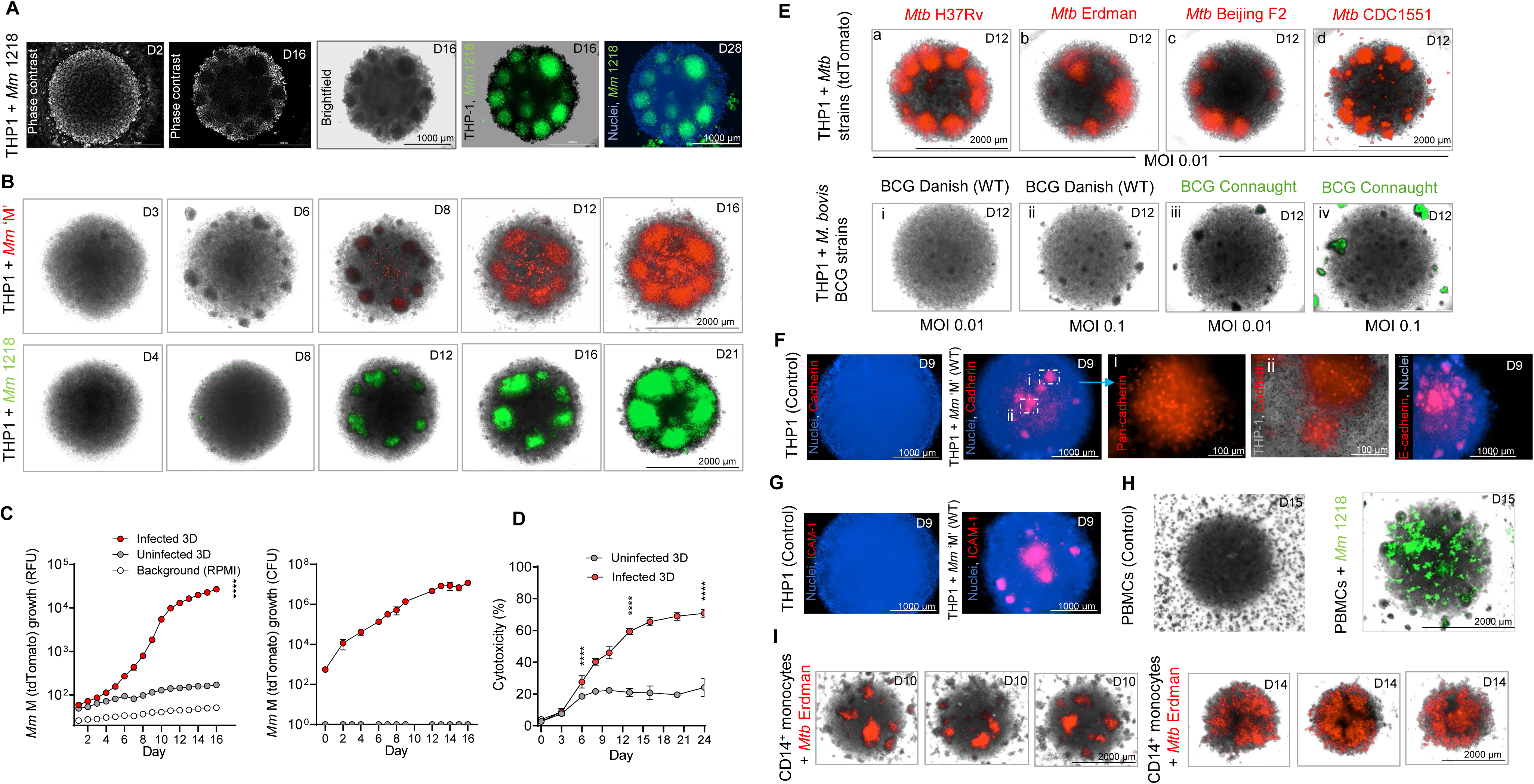
3D cell culture of human monocytes and pathogenic mycobacteria establishes a facile *in-vitro* tuberculoma model. (**A**) Microscopy of co-cultures generated using THP-1 monocytes (1×10^5^) and *Mm* 1218 GFP (MOI 0.006), demonstrating 3D spheroid formation (day 2) and mature structures (days 16 and 28) that comprise cellular aggregates resembling florid granulomatous foci in solid tuberculomas (average diameter 2098 µm). Images represent one of the three early experiments with 60 spheroids in a 3D cell culture microplate (Corning 4515). The day of image capture is indicated in each image. (**B**) Serial imaging over three weeks in co-cultures generated using *Mm* ‘M’ (tdTomato) or 1218 (GFP). Images represent one of the two experiments (n = 6 spheroids, MOI 0.006) that investigated growth kinetics in 200 µl medium per well, supplemented with fresh 50 µl medium on days 6 and 14. (**C**) *Mm* ‘M’ (tdTomato) growth in 3D co-cultures, measured using relative fluorescence unit (RFU) and colony-forming unit (CFU) counts. (**D**) Host cell cytotoxicity measured by CytoTox-Glo^TM^ assay in *Mm* ‘M’ (tdTomato)-infected or control 3D cell cultures. Data in **C** and **D** are from one of the two experiments performed (spheroids n = 204–282 infected and 18 uninfected for RFU, 6 infected per time point for CFU, and 12 infected and 12 uninfected per time point for cytotoxicity). Error bars indicate SD. \*\*\*\**p* < 0.0001 using 2-Way ANOVA with Dunnett’s test (**C**) and Welch’s t-test (**D**) compared to uninfected controls. (**E**) 3D co-cultures of THP-1 monocytes and virulent *Mtb* strains (tdTomato) using MOI 0.01 (a– d) or attenuated *M. bovis* BCG strains (WT or GFP) using MOI 0.01 (i and iii) or ten-fold higher MOI 0.1 (ii and iv). Images represent one of the two experiments, each containing 6 to 12 spheroids per strain and MOI. (**F** and **G**) Uninfected controls and 3D co-cultures of THP-1 monocytes and *Mm* ‘M’ WT (low MOI 0.004) with early developing aggregates (day 9), stained (red) for (**F**) Pan-cadherin or E-cadherin and (**G**) ICAM-1 using whole mount staining. Images represent one of the two experiments (n = 3 spheroids/marker). (**H**) 3D spheroid culture of human whole PBMCs with or without mycobacterial infection. A representative image of *Mm* 1218 GFP-infected co-culture (MOI 0.01) is from one of two experiments (n = 3 healthy donors, 54 infected and 6 control spheroids per donor PBMCs). (**I**) 3D co-culture of MACS-purified primary CD14^+^ blood monocytes (2×10^5^) and *Mtb* Erdman (tdTomato) (MOI 0.05) supplemented with autologous CD14^−^ PBMC subsets (4 ×10^5^) on day 3. The triplicate images represent 30 infected spheroids generated using purified subsets from one of the four human donors investigated.

Corresponding with the growth of cellular aggregates, bacterial fluorescence increased progressively until day 16 (**Fig 2C**). Plating THP-1 lysate on 7H10 agar plates confirmed this increased bacterial burden through colony-forming units (CFU). In this model, developed without exogenous extracellular matrix (ECM) embedding, pipetting enabled easy dissociation of mature 3D spheroids, allowing cells to be released for downstream applications without enzymatic treatment or potential changes in their established characteristics. However, THP-1 cell survival in *Mm*-infected spheroids decreased progressively over three weeks compared to uninfected ones, as shown by a cytotoxicity assay (**Fig 2D**). These results suggest that the model can facilitate quantifiable characterization of the growth dynamics of both the host and pathogen within a physiologically relevant 3D microenvironment.

During our investigations of *Mm* strains, tdTomato provided a brighter signal and less autofluorescence noise compared to GFP, consistent with the established observation that less biological autofluorescence occurs at longer wavelengths in diverse cell types and tissues. The fluorescence intensity (FI) readings further revealed more background fluorescence in the uninfected control spheroids using the GFP channel (**S1C Fig**). Therefore, we selected the *Mm* strain expressing tdTomato for further FI reading experiments. We found a good correlation between FI and bacterial number in 3D spheroids or liquid media, establishing fluorescence reading as a dependable indicator of relative bacterial number (**S1D Fig**). Co-culturing THP-1 monocytes with *Mm* in arbitrary 3D cell-culture-ware did not always yield organized tuberculoma-like structures. When we screened several vendor-provided 96-well plates with distinct microwell-surface chemistries and bottom geometries, spheroids with large granulomatous lesions were formed only in the 3D microplates with ultralow attachment (ULA)-surface and U-bottom, but not in the microplates with V-bottom or culture-surface with microcavities (**S2 Fig**). Notably, the granulomatous aggregates formed in 2D microplates or 3D polymer-microparticles encapsulating *Mm*-infected monocytes were miniature (<75 µm) and loosely organized. Among various microplates tested, only Corning ULA 3D spheroid plates with a U-bottom and neutrally charged, hydrophilic surfaces featuring a covalently bonded inert hydrogel coating produced well-defined, compact, and relatively stable granulomatous lesions, averaging 338.30 ± 60.83 µm on day 12. Corning 3D spheroid microplates were selected for future experiments.

Human TB granulomas exhibit divergent trajectories and fates [43]. We explored their evolution in a 3D co-culture model using THP-1 monocytes expressing either GFP or RFP (red fluorescent protein) along with *Mm* tdTomato or *Mm* GFP. We observed dynamic lesion formation and coalescence (**S3A Fig**), with some lesions remaining relatively stable, while others progressed with higher bacterial burdens. By day 16 post-infection, fluorescing monocytes supporting bacterial growth were predominantly observed in granulomatous aggregates, whereas the remainder of the spheroid exhibited a marked decrease in fluorescence. This suggests that mycobacteria may either promote the survival of permissive monocytes or their recruitment to these granulomatous aggregates for their benefit. Comparable organized lesions were developed with *Mtb* tdTomato strains (H37Rv, Erdman, Beijing F2, and CDC1551) using a low MOI (0.01). In contrast, live attenuated Bacillus Calmette–Guérin (BCG) strains, Danish (WT) and Connaught (GFP), produced a few (0–5) loose aggregates by day 12 (**Fig 2E**), in the absence of the virulence locus ESX-1/RD1 [44, 45]. A 10-fold higher infection dose of BCG strains produced more aggregates. Still, they remained small and loosely organized, which corroborated findings in animal models that the infection with RD1-deficient bacteria results in defects in granuloma formation and expansion [11, 45]. We used whole-mount immunostaining *in situ*, avoiding the need for single-cell imaging that requires 3D tissue clearing or labor-intensive tissue sectioning, and identified canonical tuberculous granuloma markers [46], such as cadherins, ICAM-1, and γ-catenin in the early developing granulomatous aggregates on day 9 in the *Mm* (WT)-infected spheroids (**Figs. 2F, 2G, S3B, and S3C**), which suggested that the cellular aggregates represented bona fide nascent granulomas.

We next developed 3D co-cultures using human peripheral blood mononuclear cells (PBMCs) and pathogenic mycobacteria. Whole PBMCs contain only 5–10% monocytes. They formed small granulomatous aggregates (<75 µm) (**Fig 2H**). In contrast, larger granulomas (375 ± 65 µm) developed with purified CD14^+^ monocytes and *Mtb* Erdman (tdTomato) supplemented with autologous CD14^−^ PBMC subsets after 3D spheroid formation (**Fig 2I**). No apparent difference in granuloma structure was observed with or without lymphocyte-rich CD14^−^ PBMC supplementation. However, these granulomas of primary monocytes, which lack self-renewal capabilities, progressed quickly and began to disintegrate by day 14 due to the absence of continuous monocyte-derived macrophage influx required to sustain their structural organization. These results collectively demonstrate that the ‘mycobacteria-in-spheroid’ co-culture establishes a facile 3D model that mimics human tuberculomas.

### Crucial attributes and microenvironments in human tuberculomas develop in the 3D model

To assess the phenotypic and functional attributes of monocytic cells delineating the microenvironments in the model, we performed high-dimensional flow cytometry on cells dissociated from 3D spheroids. Uniform Manifold Approximation and Projection (UMAP) analysis revealed a distinct cell cluster pattern in the infected 3D model, indicating extensive transformation of infected monocytes (**Fig 3A**). The concatenated heatmap expression plots suggested higher expression levels of several cell subsets in the infected 3D model (**Fig 3B**). While THP-1 spheroids primarily contained CD45^+^HLA-DR^+^ myeloid cells, several distinct myeloid-cell subsets, resembling those in human tuberculomas [47–50], were detected over 12 days (**S4 Fig**). In the infected 3D model, greater frequencies of cells expressed phenotypes characterizing CD14^+^CD16^-^ (classical monocytes), CD14^+^CD16^+^ (intermediate monocytes), CD68^+^CD14^+^ (macrophages), CD68^+^CD16^+^ (M1 macrophages), and CD68^+^CD163^+^ (M2 macrophages) among HLA-DR^+^ cells compared to controls (**S4 Fig**). Interestingly, they also expressed phenotypes characteristic of CD11C^+^ (myeloid dendritic cells) and CD66abce^+^ (neutrophils) among HLA-DR^+^ cells and CD33^+^ (myeloid-derived suppressor cells; MDSCs) among HLA-DR^-^ cells. In addition, a substantially higher percentage of cells within these subsets, except for a neutrophil-like subset, expressed an activated phenotype over time in infected spheroids, as indicated by increased co-expression of ICAM-1 and PD-L1 (**S4 Fig)**. Similar to processes in persons and animal models with TB [46–49, 51], cells in the granulomas and spheroid core matured into macrophages, with greater expression of vascular endothelial growth factor receptor-2 (VEGFR-2) and α-tubulin (conserved in eukaryotic cells), and galactin-9 compared to control spheroids (**Figs 3C** (a, b, c) and **S5A**). *Mtb*-biofilm formation, stained for biofilm matrix, was detected in the core of infected spheroids (**S5B Fig**). These results demonstrate that monocytic cell maturation, differentiation, activation, and immunoregulation occur in the model.

**Fig. 3.**
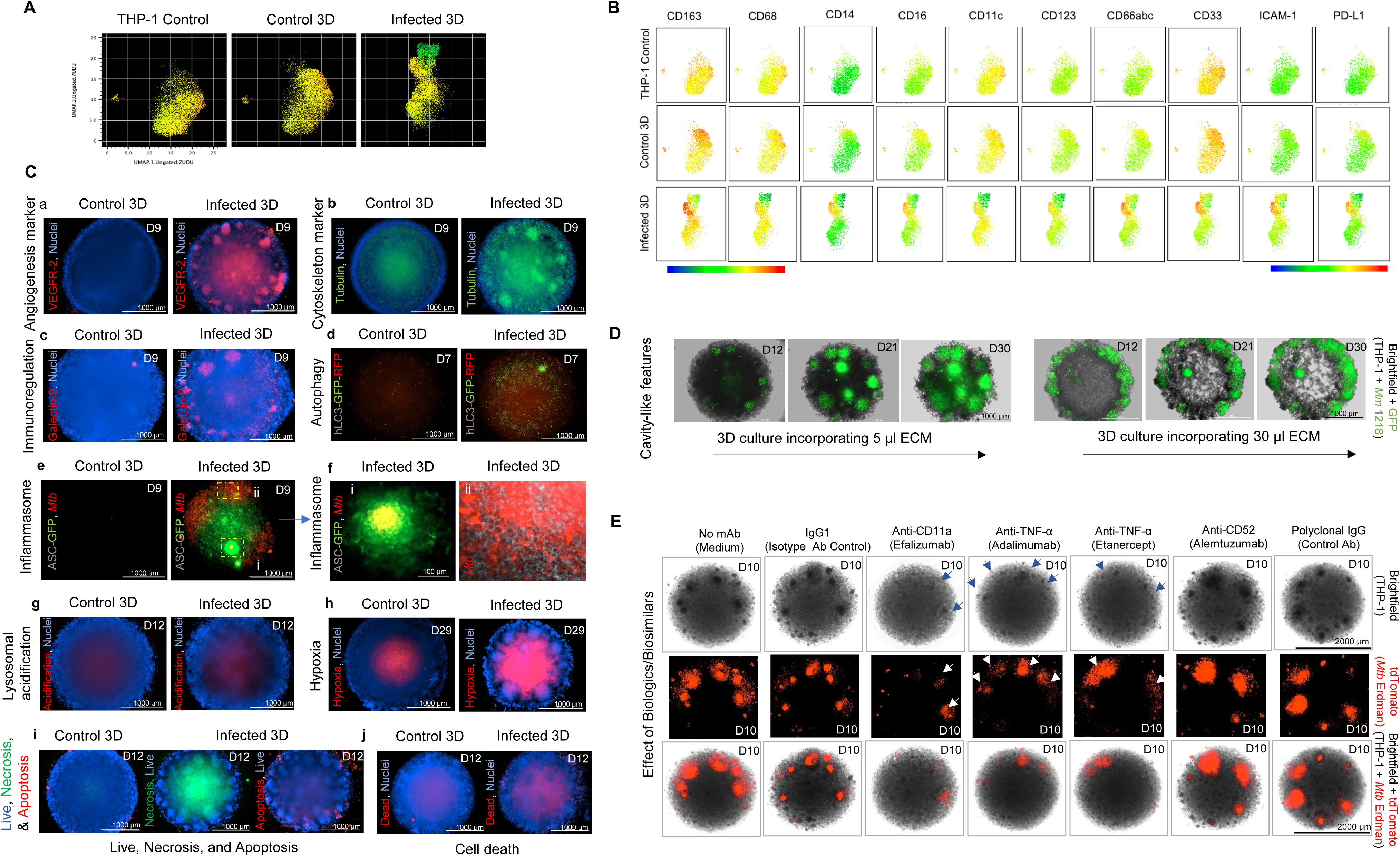
The 3D model expresses classical features and milieus in human tuberculomas and exhibits structural organization. **(A)** High-dimensional 2D UMAP graphical illustration of flow cytometry data depicting cell clusters in the THP-1 control (suspension 2D culture), uninfected 3D control, and *Mm* ‘M’ WT-infected 3D culture. (**B**) Expression of myeloid cell subset markers among three cell culture groups. The flow cytometry plots in panels **A** and **B** represent 3–6 respective cell culture samples collected and analyzed on day 9. (**C**) Increased expression of angiogenesis marker VEGFR-2 (**a**), microtubule and cytoskeleton reorganization marker α-tubulin (**b**), and immunoregulatory and endomembrane damage marker galectin 9 (**c**) in the granuloma cells detected by immunostaining (red) in the 3D co-cultures of THP-1 and *Mm* ‘M’ WT. Uninfected spheroids are used as controls. The day of image capture is indicated in each image. (**d**) 3D co-cultures of autophagy reporter THP-1 monocytes and *Mtb* Erdman WT, indicating a block in the autophagy flux with inhibition of autophagolysosome formation (yellow-green puncta) in the macrophages of the granuloma zone and granulomatous foci. (**e**) 3D co-cultures of inflammasome reporter THP-1 monocytes and *Mtb* Erdman tdTomato showing inflammasome activation (green specks) in the cells of the spheroid framework. (**f**) Dotted boxes in (**e**) show increased inflammasome activation in the granuloma epithelioid macrophages attempting to sequester *Mtb* (yellow) (box **i**) or signs of inflammasome inhibition allowing *Mtb* (red) growth (box **ii**). (**g**) Increased staining (red) in the *Mtb* Erdman WT-infected THP-1 spheroid center but not in the granuloma zone following Lyso-ID^®^ dye staining, which stains acidic organelles or environment resulting from cell death. Staining with Image-iT^TM^ Hypoxia Red Reagent (**h**), Apoptosis/Necrosis Detection Reagents (**i**), and Live/Dead Staining Reagent (intracellular cell amine sensor dye detecting dead cells) (**j**), showing increased hypoxia, necrosis, and cell death in the centers of *Mm* WT-infected spheroids. Images in panels **a**–**c** and **g**–**j** are from one of the two experiments with 3 infected and control spheroids per staining probe. Images in panels **d**–**e** represent 6 control and at least 30 infected spheroids of reporter monocytes. (**D**) Development of cavity-like features in the *Mm* 1218 (GFP)-infected 3D model of THP-1 cells following deposition of 30 µl but not with 5 µl ECM. Representative images are from one of the two experiments with at least 3–6 infected spheroids per treatment. (**E**) Treatment on day 6 with anti-TNF-α and CD11a biologics or mAb biosimilars (62.5 µg) exhibiting disruption of structural integrity (blue arrows) in the nascent granulomas on day 10 in the *Mtb* Erdman (tdTomato) infected 3D model. Images represent one of the two experiments, containing 6–30 infected spheroids treated with medium, control antibodies, biosimilars, or biologics.

*Mtb* evades macrophage innate defenses by inhibiting processes such as phagosome-lysosome fusion, phagosome acidification, autophagy, and inflammasome activation [52]. To investigate autophagy in 3D tuberculomas, we co-cultured THP-1-Difluo hLC3 reporter monocytes with *Mtb* Erdman (WT). These reporter cells express human LC3B fused with GFP and RFP [53]. Since GFP (acid-sensitive) is quenched or degraded in the acidic environment of the autophagolysosome, whereas RFP remains stable, a block in the autophagy process results in more yellow signals (red-green co-localization) compared to autophagy induction (red fluorescence). On day 7, we observed the accumulation of more yellow-green than red puncta in the outer zone, where granulomas are formed, compared to the spheroid core, indicating that autophagy induction is specifically inhibited or blocked in the *Mtb*-permissive macrophages of granulomas (**Figs 3C** (d) and **S5C**). Since *Mtb* induces assembly of the apoptosis-associated speck-like protein containing a CARD domain (ASC)-dependent inflammasomes, NLRP3 or AIM2, in the infected macrophages, we examined ASC-dependent inflammasome formation using 3D co-culture of *Mtb* Erdman (tdTomato) and THP-1-ASC-GFP reporter monocytes that stably express a gene encoding ASC-GFP fusion protein [54]. On days 7 and 9, we observed more ASC-GFP speck formation and inflammasome activation, specifically in the spheroid-scaffold cells without pronounced *Mtb* growth, and in the epithelioid macrophages of the granulomas attempting *Mtb* containment compared to granulomas permitting unfettered *Mtb* growth (**Figs 3C** (e, f) and **S5D**). These results indicate functional diversity in granuloma lesions within the 3D model and suggest inhibition of autophagy and inflammasome in *Mtb*-permissive granulomas.

Mature tuberculomas can develop hypoxic and necrotic centers over time [19]. We investigated these attributes in the 3D model. Consistent with the autophagy results, we observed a relatively more acidic environment in the spheroid core compared to the granuloma zone (**Fig 3C** (g)). Using different hypoxia probes that fluoresce in the hypoxic milieu, we found increased hypoxia in the centers of *Mm* or *Mtb* (WT)-infected spheroids compared to their peripheries and uninfected controls, with hypoxia increasing over time as the spheroids matured (**Figs 3C** (h) and **S6A**). This suggests that granuloma lesions, which support bacterial proliferation, tend to develop in oxygen-sufficient regions. While hypoxia is generated in both control and infected spheroids, owing to the sizable (>2000 µm) 3D structure, macrophage transformations with granuloma conglomeration likely exacerbate hypoxia in the solid tuberculomas. Although hypoxia induction in *Mtb*-infected macrophages enhances their microbicidal capacity [29, 55], excessive hypoxia and HIF-1 activation can contribute to necrosis and matrix destruction. Likewise, we observed increased cell death, characterized mainly by necrosis rather than apoptosis, in the central regions of infected spheroids using cell-death probes (**Figs 3C** (i, j), **S6B,** and **S6C**).

Central necrosis and ECM destruction drive cavitation in human tuberculomas [19]. Despite the heightened activity of matrix metalloproteinases (MMPs) and matrix remodeling in 3D *in-vitro* granuloma models with exogenous ECM [27], cavitation has not been reported. In a 3D model of *Mm* 1218 (GFP), we investigated the effects of ECM inclusion, which consisted of human collagen type 1 and fibronectin. The *Mm* 1218 strain enabled a co-culture of longer duration (>50 days) with THP-1 monocytes when a low MOI (0.005) was applied, and the culture medium (50%) was exchanged with fresh medium once a week. By adding increasing quantities of ECM (5 to 50 µl) post-spheroid formation, to simulate heightened ECM deposition in human tuberculomas, we observed cavity-like features in infected spheroids with higher quantities (≥30 µl) of ECM after 21 days of co-culture. In contrast, uninfected spheroids and those with a 10-fold lower MOI did not exhibit cavitary morphology (**Figs 3D** and **S7**). These results collectively demonstrate solid, necrotic, and cavitary transformations in the tuberculoma model.

Tumor necrosis factor (TNF) plays a crucial role in maintaining granuloma integrity by restricting bacterial growth and preventing macrophage necrosis. However, excess TNF can trigger programmed necrosis in infected macrophages [56, 57]. We investigated the impact of human TNF blockers, etanercept—a fusion protein of TNF receptor type-II fused to the Fc fragment—and a human anti-TNF monoclonal antibody (mAb) adalimumab biosimilar, on bacterial burden and granulomas in 3D co-cultures of THP-1 cells and pathogenic mycobacteria (**Fig 3E**). We simultaneously investigated 16 additional human recombinant mAb or antagonist protein biosimilars (**S8 Fig**), equivalent to those approved for treating cancers, inflammatory disorders, autoimmune diseases, and other conditions. Following a dose titration, treatments were administered at high (500 ng) and low (62.5 ng) doses before infection or on day 6. By day 10 in 3D co-cultures of *Mtb* Erdman (tdTomato), developing granulomas exhibited a loss of organizational integrity upon exposure to TNF blockers (blue arrows; **Fig 3E**) but not to isotype or polyclonal control antibodies. This occurred regardless of the apparently comparable bacterial burdens in anti-TNF-treated and control co-cultures (white arrows; **Fig 3E**), the treatment dose used, or the timing of treatment exposure (**S8A Fig**). In *Mm* M (tdTomato)-infected spheroids, exposure to anti-TNF mAbs resulted in multiple miniature granulomas (**S8B Fig**).

The biosimilar mAb targeting CD11a, a cell-surface integrin involved in cellular adhesion, but not CD52, which plays a role in anti-adhesion, caused the early loss of granuloma integrity and reduced *Mtb* burden by day 10 (**Figs 3E and S8A**). Yet, relatively normal, mature granuloma formations with comparable *Mtb* loads progressed by day 14 in the 3D spheroids treated with anti-CD11a mAbs, TNF antagonists, or control antibodies (**S8C** and **S8D Fig**), confirming that TNF is required for the integrity of nascent granulomas, rather than the formation [58, 59]. Additionally, exposures to biosimilars targeting VEGF, integrin α4β7, CD30, insulin growth factor-1 receptor (IGF1R), IL-6Rα, IL-1β, or IL-1R on day 6, slowed granuloma expansion by day 10, as evident by their smaller size (**S8E Fig**), compared to medium alone, isotype control mAbs, or biosimilars targeting CTLA-4, PDL1, and CD52. However, by day 14, only high doses (500 ng) of anti-VEGF, anti-α4β7, anti-CD30, and anti-IGF1R mAbs effectively inhibited *Mtb* loads, while anti-CD52 (62.5 ng) increased loads relative to controls (**S8F Fig**). Together, these results demonstrate the development of human tuberculoma-like traits and milieus in the model, highlighting the potential for HCS applications.

### Protein biomarker profiling by multiplex arrays provides insight into the host-pathogen interactions leading to cavitary transformation

To examine molecular reprogramming in macrophages leading to granuloma formation and host determinants of cavity formation, cell lysates and culture supernatants from 3D cell cultures were analyzed using the Quantibody^®^ Human Kiloplex Proteomics Assay (**Fig 4**), quantifying 1,000 central proteins involved in immune processes. The study revealed substantial fold-changes in protein expression between THP-1–*Mm* 3D co-cultures generated with or without exogenous ECM compared to uninfected 3D controls. The results helped identify the modulation of protein expression levels and the effect of ECM deposition on changes in the protein profile, leading to cavitation in 3D tuberculomas. The raw quantities of 1,000 proteins in the cell lysates and culture supernatants are presented in the **S1 Table.** The top-up-regulated or down-regulated proteins are shown as dot plots (**Fig 4A** and **4B)**. Notably, levels of chemokines MCP-2, MIG, GROα/CXCL-1, MIP-3α, I-TAC/CXCL-11 (458.6–103.6-fold), and MIP-1α, IP-10/CXCL-10, IL-8/CXCL-8, and MCP-4 (>20-fold) showed a marked increase compared to controls in the cell lysates of 3D co-cultures without ECM (**Fig 4A**). These same chemokines were also increased (556.5–18.2-fold) in the cell lysates of 3D co-cultures with ECM. Proteins increased solely in the cell lysates of 3D co-cultures without ECM include nucleoporin NUP-85, phagocytic receptor CEACAM-3, growth factor PD-ECGF, FABP-4, enzyme DNMT3A, microtubule binding DCTN-1, and kinase Flt-3 (**Fig 4A**). Many of the above-identified chemokines were also overexpressed in the supernatants of 3D co-cultures with or without ECM, along with LDL-R, MMP-9, HO-1, and other proteins (**Fig 4B**). Albumin and RBP-4, potential biomarkers of TB disease and treatment response [60], as well as the gluconeogenic enzyme PCK-1, IL-17RD, and aurora kinase A activator BORA, and Siglec-1, showed markedly increased levels with the addition of ECM in the supernatants. At the same time, galectin-4, galectin-7, langerin, BCL-10, and ceramidase ASAH2 decreased in the supernatants of co-cultures with ECM. Analysis of proteomics data from lysates of 3D co-cultures without ECM identified the upregulation of several proteins involved in cell adhesion, angiogenesis, matrix remodeling, immunoregulation, and other cellular processes accompanying granuloma formation (**S2 Table**). We further confirmed the expression of five protein biomarkers in the 3D co-culture using the ELISPOT assay (**S9 Fig**).

**Fig. 4.**
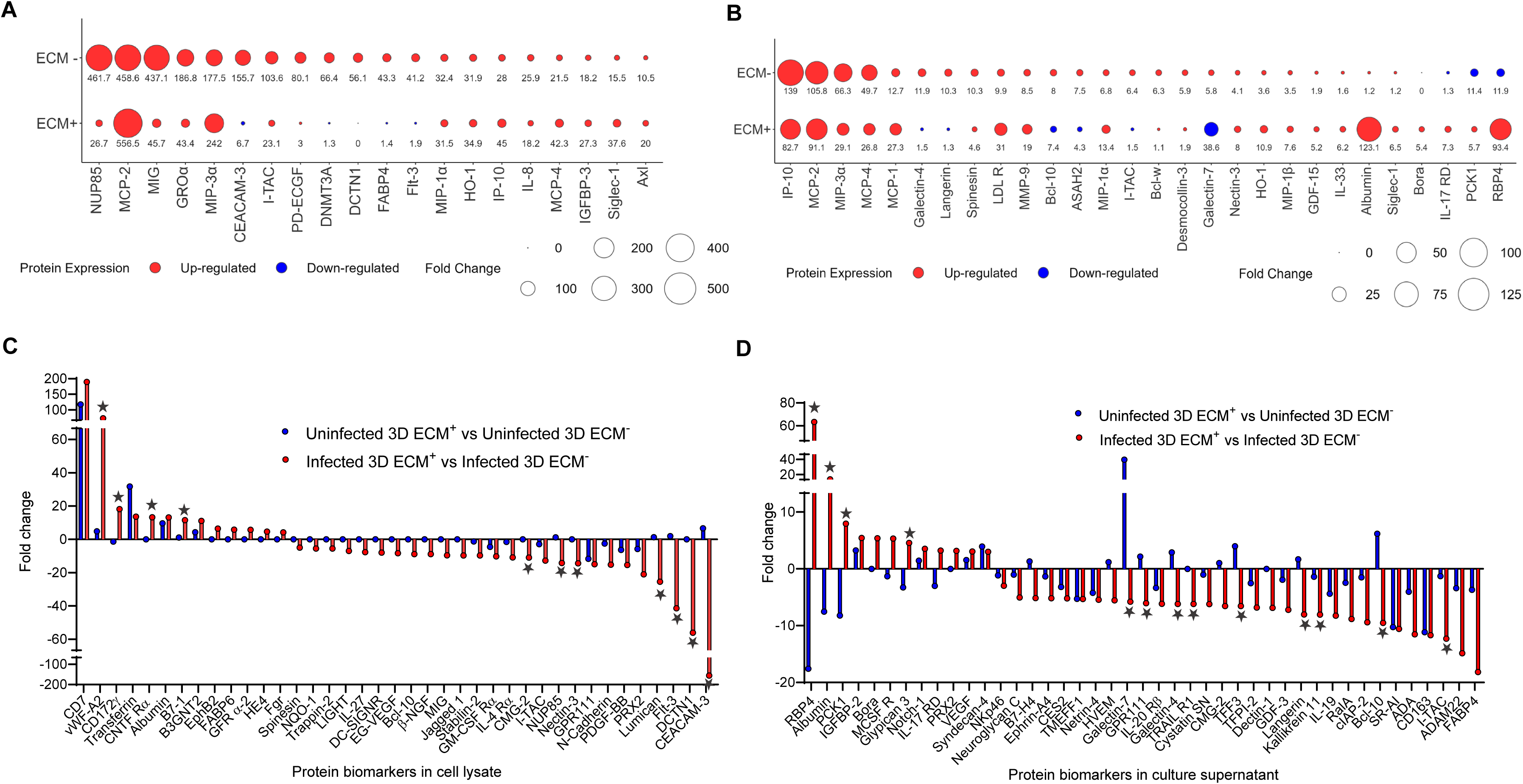
Protein biomarker profiling in the 3D tuberculomas with or without ECM incorporation and cavitary transformation. The cell lysates (**A** and **C**) and culture supernatants (**B** and **D**) were isolated on day 16 from 3D spheroid cultures of *Mm* 1218-infected or uninfected THP-1 monocytes with or without ECM (30 µl) addition (MOI 0.005) and subjected to Proteomics Arrays. (**A**–**B**) Relative fold-change in the levels of top differentially-expressed proteins in the infected compared to uninfected cultures is presented (at least >5-fold change in either with or without ECM). Dot plots of top proteins with change in expression in the cell lysates (**A**) and culture supernatants (**B**) of the infected compared to uninfected cultures are shown. ECM− indicates 3D co-cultures without ECM, and ECM+ indicates 3D co-cultures with ECM. The size of the dots indicates the relative fold change in protein levels (pg/ml), and the average fold-change values are shown at the bottom of each dot. Red dots denote up-regulation, and blue dots denote down-regulation in protein expression levels. (**C**–**D**) Protein markers with a change in expression levels in the cell lysates (**C**) and culture supernatants (**D**) following ECM addition and cavity formation. Red bars show fold-change in protein expression levels in the infected 3D co-cultures with ECM (showing cavitation) compared to those without ECM (no cavitation). Blue bars indicate the fold-change in protein expression levels in the uninfected control 3D cultures with ECM compared to those without ECM (both with no cavitation). Stars indicate >10-fold change in infected compared to uninfected. (**A–D**) n = 30 spheroid cell lysate or culture supernatant pools per condition. Each pooled sample was analyzed in quadruplicate. The average fold-change values are plotted.

The ECM plays a critical role in regulating host-pathogen interactions and granuloma necrosis. The results revealed that ECM deposition exclusively modulated the expression of specific molecules in *Mm*-infected compared to control 3D cultures (**Fig 4C** and **4D**). In the lysates or supernatants of infected cultures, ECM substantially (>10-fold) increased vWF-A2, CD127g, CNTFR, B7-1, RBP4, albumin, PCK-1, M-CSFR and decreased CEACAM-3, DCTN1, Flt-3, lumican, Nectin-3, NUP85, ITAC/CXCL11, BCL-10, Kallikrein 11, Langerin, TFF3, TRAIL-R1, Galectin-4, GPR-11 and galectin-7 levels compared to control 3D cultures. Many of these molecules are known to interact with collagen-rich ECM and participate in matrix remodeling, suggesting that their expression is modulated by ECM addition, leading to cavitation in tuberculomas. Further studies are required to understand the mechanisms behind the ECM’s role in these processes.

### RNA sequencing offers insight into the transcriptional changes accompanying granuloma formation

To identify host-pathogen interactions leading to granuloma formation and expansion in the model, we compared the transcriptomic profiles of cells isolated from 3D co-cultures of THP-1 cells and *Mm* ‘M’ at four different time points with those from control spheroids (**S10 Fig**). Principal component analysis revealed differential clustering between infected samples and the matched uninfected controls (**Fig 5A**). Unsupervised hierarchical clustering of the expression data confirmed the clear separation of clusters of infected and control samples (**S11A Fig**). The quality of transcriptomic data is presented in the **S3 Table**.

**Fig. 5.**
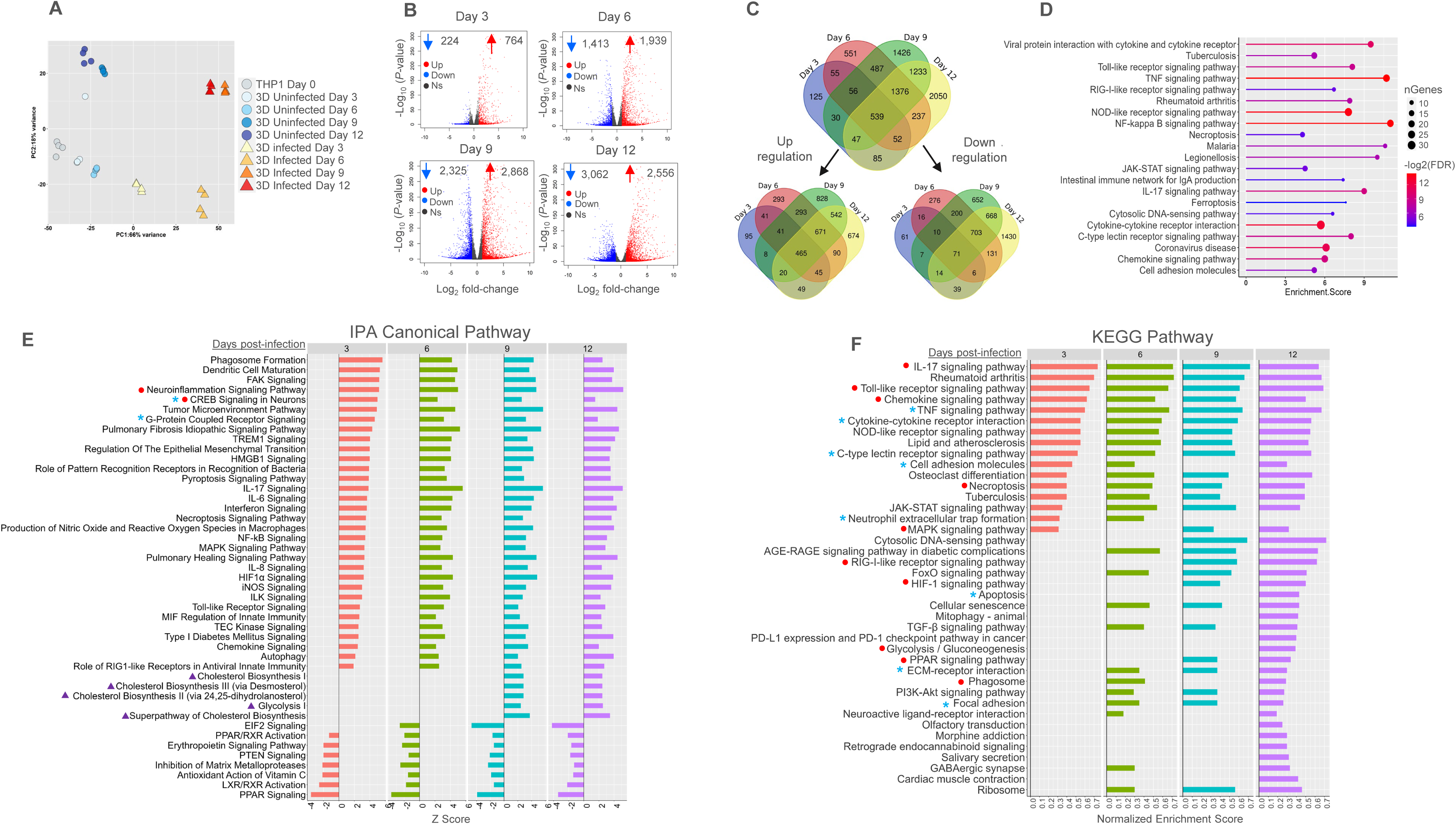
Transcriptomic analysis of host cells from the 3D tuberculomas. (**A**–**F**) RNA-Seq using cells isolated from the *Mm* ‘M’ (tdTomato)-infected or control 3D cultures of THP-1 on days 3, 6, 9, and 12 post-culture (MOI 0.005). n = 30 spheroids per time point and condition. The data are from four biological replicates. (**A**) The PCA plot of RNA-Seq samples. (**B**) The volcano plots show significantly up- and down-regulated genes in the infected compared to the control samples. Red dots denote up-regulated genes (log2 fold-change >1 and adjusted *p*-value < 0.05), blue dots denote down-regulated genes (log2 fold-change <-1 and adjusted *p-*value < 0.05), and black dots represent genes with non-significant changes in expression (adjusted *p-* value ≥0.05). (**C**) The Venn diagrams of the up-regulated and down-regulated genes shared between the four time points. (**D**) The shared differentially expressed genes (DEGs; n = 539) subjected to GSEA using ShinyGo, and the top enriched KEGG pathways are presented as a lollipop plot. (**E–F**) The enriched canonical pathways altered before (day 3), during (day 6), and after (days 9 and 12) granuloma formation in the infected compared to control 3D cultures. (**E**)The set of DEGs (in **D**) subjected to GSEA using the Qiagen IPA canonical pathway analysis tool is plotted. The z-scores of the most enriched canonical pathways (n = 45), ranked based on the day 3 scores, are presented. Red circles denote neurological signaling pathways, purple triangles denote cholesterol or glycolysis-related pathways, and blue asterisks denote pathways showing the most activation on day 3 before granuloma formation. (**F**) The enriched KEGG pathways altered before, during, and after granuloma development in the infected compared to the control 3D cultures. The same sets of DEGs (in **D)** are subjected to GSEA using the clusterProfiler package in R. The top 40 most enriched KEGG pathways are presented as bar plots. Red dots represent pathways shared with IPA analysis. Blue asterisks denote pathways associated with granuloma response identified by the GSEA.

We analyzed differential gene expression between infected and control samples using DESeq2 tools (**Figs 5B** and **S11B**) and identified 988, 3,352, 5,193, and 5,618 differentially expressed genes (DEGs) on days 3, 6, 9, and 12-post-infection, respectively, with substantial upregulation (n = 2,868 genes) on day 9 and downregulation (n = 3,062 genes) on day 12, indicating extensive transcriptomic changes. We explored the core DEGs shared among 4 time points and found 465 up-regulated and 71 down-regulated genes common across all time points (**Fig 5C** and **S4 Table**). Gene Set Enrichment Analysis (GSEA) using ShinyGO of the shared DEGs revealed overrepresentation of genes in multiple immune response pathways (**Fig 5D**), including cytokine-cytokine receptor interaction, TLR, TNF, NOD-like receptor, IL-17, NF-κb, C-type lectin, and chemokine signaling pathways.

Among the shared 539 DEGs (**Fig 5C**), we observed dynamic changes in the gene expression at different time points post-infection. Specific genes, including TNFAIP6, IGFBP3, IFI27/44L, IFIT1, CCL4, IL6, CXCL2, CXCL8, CXCL14, MMP8, SUGCT-AS1, AIM2, MET, and IER3, exhibited progressive upregulation, reaching a peak on day 12 (**S4 Table**). In contrast, genes such as IDO1, MMP1, MMP12, ITGB8, IL1α, CCR7, CCL20, CXCL1, CXCL3, and IL1β reached their maximum expression on day 9, before declining by day 12. Early activated genes, such as CCL8, ADAM7-AS1, CXCL10, CXCL11, and VCAM1, exhibited rapid upregulation of over 100-fold by day 3 (before granuloma formation). Among the down-regulated genes, STYXL2, EVPL, KIT, and CEACAM6 showed a peak decrease on day 9, while the rest showed a steady decline in expression as the infection and granulomas progressed. These results reveal dynamic modulation of transcripts for several chemokines, cytokines, and MMPs during granuloma formation. Additionally, they suggest that the upregulation of previously unrecognized TNF-α-inducible protein 6, insulin-like growth factor binding protein-3, interferon-induced proteins (IFI44L, IFI27, IFIT1), lncRNA (SUGCT-AS1), and integrin beta-8 in macrophages plays a role in granuloma development.

We performed GSEA using Ingenuity Pathway Analysis (IPA) (**Fig 5E**) and Kyoto Encyclopedia of Genes and Genomes (KEGG) analysis using clusterProfiler (**Fig 5F**) to gain insights into the pathways underlying host-pathogen interactions and granuloma formation. Among the top 45 differentially regulated canonical pathways identified using IPA, 25 pathways mainly related to innate immune responses remained activated throughout the infection period. Interestingly, two nervous system-related pathways (red dots) were markedly activated, while seven pathways, including PPAR signaling, LXR/RXR activation, inhibition of MMPs, and PTEN signaling, were consistently inhibited. In contrast, five cholesterol and glycolysis-related pathways (purple triangles) were substantially activated, and EIF2 signaling, which regulates bacterial invasion, was inhibited only after granuloma onset (**Fig 5E**). Ten of the top 40 differentially regulated pathways identified using KEGG overlapped with those from IPA (Red dots; **Fig 5F**). The pathways identified solely using KEGG included those previously associated with TB granulomas, including TNF signaling, cytokine-cytokine receptor interactions, cell adhesion, focal adhesion, and ECM-receptor interaction pathways. Corroborating our identification of proteins related to cell adhesion, epithelial-mesenchymal transformations, cytokine and chemokine responses, angiogenesis, autophagy, matrix-remodeling, necroptosis, and pyroptosis, many genes associated with these pathways were differentially expressed (**S12 Fig** and **S5 Table**). Furthermore, we identified several DEGs linked to neuroactive ligand-receptor interactions, checkpoints, and immunoregulatory pathways. Together, these results demonstrate widespread transcriptional reprogramming accompanying granuloma formation and support the development of tuberculoma-like features.

Our model, despite using only myeloid cells, exhibited more DEGs than the models described previously [61], and shared 529 DEGs with a polymer-encapsulated 3D model of *Mtb*-infected human PBMCs with collagen (1,176 total DEGs) and 728 DEGs with human lymph node TB samples (2,106 DEGs) (**S13A** and **S13B Fig**). It also shared 796 DEGs with the lung tissues of diversity-outbred mice (2,992 DEGs) and 843 DEGs with those from non-human primates (NHP; 4,404 DEGs) (**S13C Fig**) that developed TB disease [62]. Notably, our model showed upregulation of >87% of genes from the previously reported 16-gene signature that predicted the onset of TB disease across host species (58) and a 70-gene signature consistently upregulated in the blood of persons with TB compared to those with latent infection [63] (**S13D** and **S13E Fig**). These results highlight that key transcriptomic changes identified in human and animal TB specimens are represented in our model.

### 3D tuberculomas in microplates can be employed as an HTS-compatible platform to identify potential HDT compounds

Given the central role of macrophages in granuloma formation and the difficulties in targeting bacilli in tuberculous granulomas [14, 64], we investigated the use of 3D tuberculomas in microplates to identify therapeutics that modulate macrophage responses. HDTs modulating the perturbed granuloma-macrophage defense pathways likely have the maximum impact in controlling early mycobacterial infection. We screened a custom library of 65 known potential HDT compounds (**Fig 6A** and **S6 Table**), including 44 United States Food and Drug Administration (FDA)-approved drugs, 10 drugs in clinical trials or human use for diverse indications, and 11 investigational compounds. They were selected from the literature published between 2000 and 2020 based on their known capability to reduce mycobacterial burdens in 2D cell cultures or induce protective responses in animal models or humans. These compounds span a broad range of categories and exhibit diverse host-directed effects, including anti-cancer, anti-inflammatory, kinase modulation, ion-channel blocking, neuroleptic, antioxidant, epigenetic modulation, anti-diabetic, and other effects **(S6 Table)**. Their efficacy in reducing bacterial growth and granuloma lesions was assessed after a single treatment of 20 µM on day 6 in 3D co-cultures, using FI measurements and imaging 6–8 days post-treatment.

**Fig. 6.**
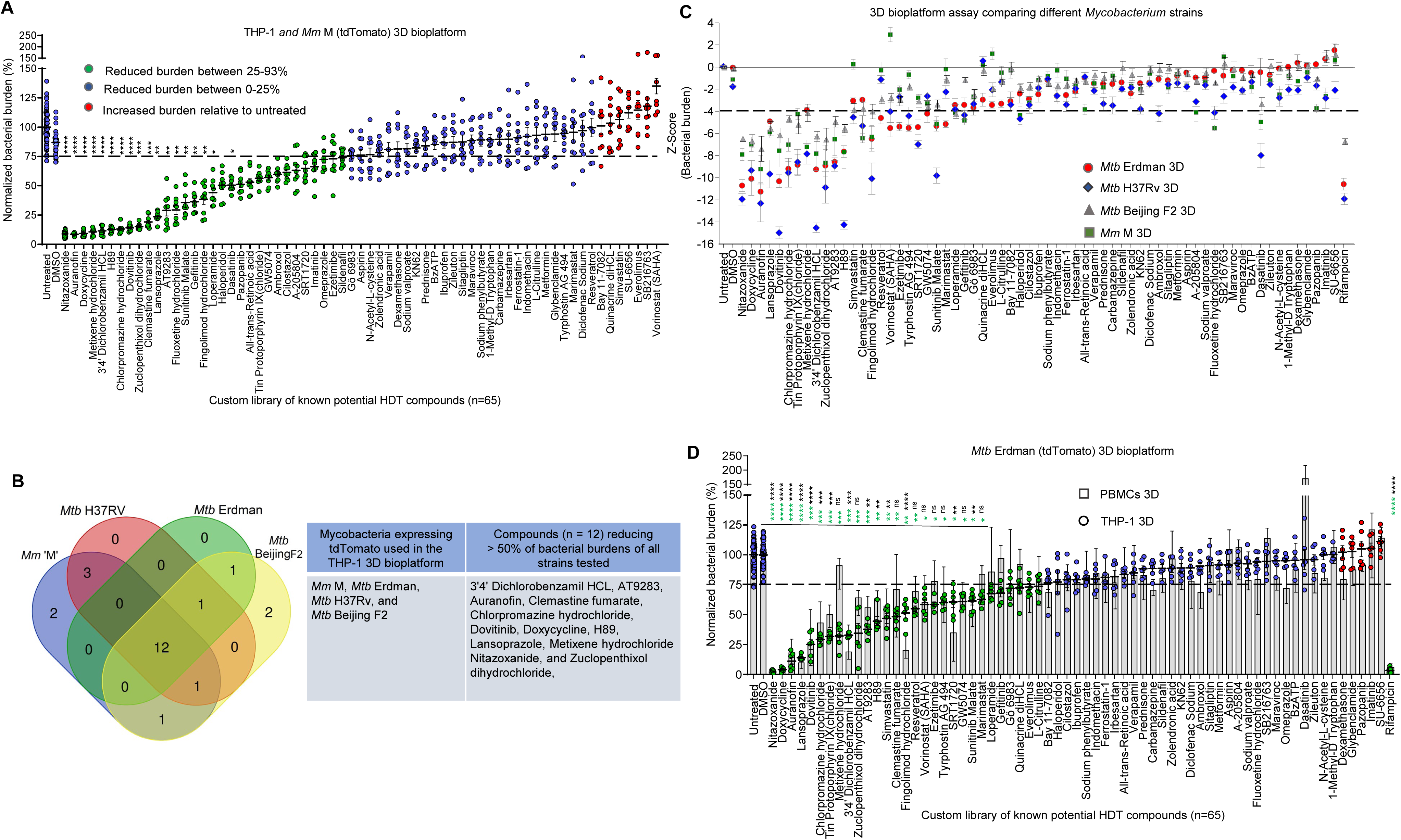
Identification of HDT compounds inhibiting pathogenic mycobacteria in the 3D tuberculoma bioplatform. (**A**) Screening of chemical compounds in a custom library of 65 known potential HDT compounds at 20 µM in the 3D tuberculomas of THP-1 monocytes and *Mm* ‘M’ (tdTomato). Tuberculomas were treated on day 6, and fluorescence intensity (FI) was measured on day 12 (Day 6 post-treatment). Results are expressed as a normalized bacterial burden (%). Data are from three independent experiments (n = 9 tuberculomas/compound and 36–72 tuberculomas/DMSO or untreated control), and filled circles display bacterial burden in individual tuberculomas. (**B**) The Venn diagram illustrates the 12 ‘top-hits’ identified in the compound screens in 3D tuberculomas of four individual strains (*Mm* and *Mtb* tdTomato). The 12 ‘top-hit’ compounds, which reduced bacterial burdens by >50% compared to untreated controls, are listed in the table. (**C**) Results of the compound screens in the 3D tuberculoma bioplatform of THP-1 monocytes, investigating inhibitory effects on four different mycobacterial strains. Rifampin served as a positive control for bacterial burden reduction. The average z-score and standard error for each compound and control, screened in 2–4 experiments per mycobacterial strain, are displayed using four different symbols. The dashed line represents a robust cutoff at a z-score of −4 (2× traditional cutoff of z-score −2). (**D**) Identification of HDT compounds inhibiting *Mtb* Erdman (tdTomato) in the 3D tuberculomas of PBMC subsets (purified CD14^+^ monocytes supplemented with CD14^−^ subsets) or THP-1 monocytes. Data are from two THP-1 experiments and four human donor PBMC experiments (n= 6–12 tuberculomas/compound). (**A**, **C,** and **D**) The mean ± standard error of the mean (SEM) is shown. \**p* < 0.05, \*\**p* < 0.01, \*\*\**p* < 0.001, and \*\*\*\**p* < 0.0001 compared to DMSO using Kruskal-Wallis with Dunn’s post-hoc test. Green and black asterisks in **D** denote significance levels in THP-1 and PBMC co-cultures, respectively.

In total, 31 compounds (47.6%) reduced average *Mm* burdens by more than 25% in 3D tuberculomas compared to the untreated (no-drug control) group. However, the burden-reducing effects of only 18 compounds were statistically significant (*p* = 0.049 to < 0.0001) compared to DMSO (drug-carrier) controls (**Fig 6A**). Interestingly, 7 compounds increased *Mm* burdens compared to controls, albeit insignificantly. Significant anti-bacterial effects of a known pathogen- and host-directed drug nitazoxanide and antibiotic rifampicin (*p* < 0.0001) were detected in the model (**S14 Fig**), which served as positive controls and validated the assay’s reliability. The assays performed in the *Mm*-infected model demonstrated excellent quality and robustness for HTS applications (Z’-factors >0.5) using a low MOI (range, 0.007–0.012) (**S14A Fig**). For slow-growing *Mtb* (tdTomato) strains representing a heterogeneous group of genotypes, a relatively higher MOI (0.012–0.05) was necessary for excellent assay quality (Z’-factor >0.5) and reliable results (**S14B Fig**), especially for H37Rv, Erdman, and Beijing F2 strains with diverse growth rates in THP-1 monocytes [41]. For the CDC1551 strain, which proliferates in small clusters outside the 3D structure and appears to evade containment within the spheroid, the assay quality could not reach an excellent level, even with an MOI of 0.05, despite the formation of granuloma lesions. CDC1551 is known to trigger a quicker and stronger immune response, although it is not more virulent than other strains [65]. Additional studies are required to understand its ability to evade early immune response, leading to granuloma formation. The assay performance was validated across multiple plates (n = 10) using the H37Rv strain, at an optimum MOI of 0.025. Reproducible *Mtb*-growth inhibition using nitazoxanide and rifampicin was detected across all wells of plates tested, and results in each plate showed excellent assay quality (**S14C Fig**). These results collectively informed the suitability of the tuberculoma model as an HTS-compatible bioplatform.

Using *Mtb* strains H37Rv, Erdman, and Beijing F2 in 2−4 independent screens, we identified 29−31 compounds that reduced *Mtb* burdens by >25% compared to untreated controls (**S15 Fig**) despite strain-specific differences in bacterial growth inhibition by HDT compounds. Using the *Mtb* CDC1551 infection model and the same parameters, we identified 39 active compounds. Since this platform did not exhibit excellent assay quality, we did not consider these results and the strain for further investigations. A comparison of the screening results in the BSL-2 (*Mm* ‘M’) and BSL-3 (three *Mtb* strains) platforms revealed considerable overlap in compounds that reduced bacterial burdens by >50% (**Fig 6B**), notwithstanding strain-specific variations in dependence on host cellular factors for intracellular survival [41]. These ‘top hits’ include 12 drugs with diverse known host-directed effects, and they were effective in all four infection models (**Fig 6B**). We compared the list of ‘hits’ identified using >50% bacterial inhibition with the ‘hits’ discovered using a statistical *p*-value < 0.05 and composite z-scores with a robust cutoff of ≤−4 in all infection models. We found that the list of ‘top hits’ identified using both methods was almost identical (**Fig 6C**). Metixene hydrochloride and clemastine fumarate were exceptions. Yet, their z-scores (<−3) were still below the traditionally used cutoff (z-score <−2). This high degree of agreement (>90%) in the ‘top hits’ identified in *Mm* and *Mtb* infection models reassured us about the performance and validity of the results in our BSL-2 and -3 platforms. Next, we considered the capability of 65 compounds to mitigate immunopathology by reducing granuloma lesions. The 12 ‘top-hit’ compounds also effectively alleviated lesion burdens (**S16 and S17 Figs**), except clemastine fumarate and metixene hydrochloride, which did not efficiently reduce lesions in the Erdman model. Despite consistently reducing the average lesion burdens by >25% in all infection models, the effect was not statistically significant for lansoprazole, dovitinib, 3’4’ dichlorobenzamil HCL, and AT9283 in the Erdman model.

Lymphocytes play a crucial role in the antimycobacterial response in tuberculous granulomas, and the performance of HDT compounds in the context of both innate and adaptive immune cells is paramount. We next investigated 65 compounds in a 3D tuberculoma model of *Mtb* Erdman-infected human CD14^+^ monocytes, supplemented with autologous PBMC subsets comprising lymphocytes and CD14^−^ myeloid cells (**Fig 6D**). There was a substantial agreement in screening results between the THP-1 monocyte and PBMC platforms. Out of 29 effective compounds identified in the Erdman-infected THP-1 model, 23 were also effective in the PBMC model, using a bacterial burden reduction of >25% as a parameter. However, statistical significance compared to DMSO was limited to only 23 compounds in the THP-1 model and 14 in the PBMC model due to variations in individual donor responses (*p* = 0.0063 to <0.0001). Nonetheless, when the percent normalized *Mtb* burdens for individual compounds in two cellular models were compared, no significant difference in efficacy was observed for 64 compounds. One compound, metixene hydrochloride, demonstrated significantly greater effectiveness in the THP-1 compared to the PBMC model (p < 0.05), suggesting that it may not be effective in the context of primary human immunity.

Next, we considered the effectiveness of compounds in five infection models using THP-1 monocytes or PBMCs (**Figs 6**, **S15**, and **S16**). None of the FDA-approved anti-diabetics (metformin, glyburide, sitagliptin) or anti-inflammatory drugs (dexamethasone, prednisone, aspirin, ibuprofen, indomethacin, diclofenac sodium) reduced bacterial burdens by >25% in these models. Additionally, several previously identified potential HDT candidates (**S6 Table**), including 1-methyl-D tryptophan (IDO inhibitor), zileuton (5-lipoxygenase inhibitor), BzATP (P2X7 receptor agonist), verapamil (calcium channel blocker), zoledronic acid (γδ T-cell response inducer), N-Acetyl-L-cysteine (anti-oxidant), sodium valproate (anti-convulsant and sodium channel blocker), L-citrulline (nitric oxide booster), sodium phenylbutyrate (histone deacetylase inhibitor), maraviroc (CCR5 receptor antagonist), and irbesartan (anti-hypertensive and PPARγ activator), were ineffective as standalone treatments, except for indomethacin and L-citrulline, which showed modest inhibition in THP-1 model using Beijing-F2 or Erdman.

Not all compounds identified as ‘hits’ in earlier studies using mycobacterium-infected 2D cell-culture screens (**S6 Table**) proved effective in our 3D model. While previous studies administered test compounds during or shortly after mycobacterial uptake by host cells, our screening focused on a later time window in infection, specifically during the establishment of granuloma milieus. For example, of the nine compounds previously identified amongst ‘top hits’ from a 1260 pharmacologically active compound (LOPAC^®^) library after drug-repurposing 2D screens in H37Rv-infected human MelJuSo cell line or monocyte-derived macrophages [39], only four— dovitinib, H89, 3’4’ dichlorobenzamil HCL, and GW5074—consistently reduced bacterial loads by 25−90% across all infection models (**Figs 6**, **S15**, and **S16**). Three compounds—quinacrine diHCl, tyrphostin AG459, and haloperidol—showed varying efficacy (>25% reduction in one to three models), while SU6656 and SB216763 were ineffective.

Of importance, the previously reported ‘hit’ compounds, which were shown to inhibit mycobacterial burdens in animal models, also demonstrated effectiveness in our infection models. Notably, a gastric proton-pump inhibitor (PPI), lansoprazole, confirmed in the mouse model and identified as a potent ‘hit’ in the *Mtb*-infected MRC-5 fibroblast cultures after screening a Prestwick Chemical Library^®^ [66], showed significant effectiveness across all our infection models. Similarly, four compounds—chlorpromazine hydrochloride, metixene hydrochloride, clemastine fumarate, and zuclopenthixol dihydrochloride—identified amongst the lead HDT candidates after screening a Prestwick Library^®^ of FDA-approved compounds in the *Mm*-infected zebrafish larvae model [67], emerged as ‘top hits’ in our infection models, although metixene hydrochloride was ineffective in the PBMC model. Additionally, antimicrobial drugs nitazoxanide and doxycycline, which exhibit host-directed effects in animal models [68, 69], showed potent anti-mycobacterial activity in our infection models. These results support the capability of our platform to identify *in vivo* effects of known host-directed and pathogen-targeting drugs in 3D *in-vitro* tuberculomas. In a follow-up screening for compounds that more effectively inhibit bacterial growth in a 3D microenvironment, we evaluated compounds in the H37Rv-infected 2D THP-1 culture, focusing on the later stages of infection as in 3D co-culture (**S18 Fig**). While AT9283 and simvastatin demonstrated superior reduction in bacterial burden in the 3D tuberculoma model, imatinib and quinacrine dihydrochloride were more effective in the 2D cultures. Several compounds previously identified as effective in traditional 2D cultures during an early time window in infection were found to be ineffective once the pathogen established a sustainable equilibrium with the host’s intracellular milieu and developed granulomas.

### A chemical screen in the mycobacterial 3D broth culture identifies antimycobacterial compounds

To distinguish between compounds with host-directed versus direct anti-microbial effects, we treated *Mm* ‘M’ or *Mtb* Erdman (tdTomato) broth cultures with compounds in 3D ULA microplates. We monitored bacterial growth as axenic 3D cultures and measured FI on days 6 and 8 post-treatment (**S19 Fig**). Thirteen compounds previously described as having some anti-mycobacterial activity in the broth cultures, in addition to host-directed effects (e.g., antibiotic and MMP inhibitor doxycycline) (**S6 Table**), were also tested. As expected, doxycycline and rifampicin exhibited significant inhibitory activity against *Mm* and *Mtb* in 3D axenic cultures compared to untreated controls (*p* < 0.0002), validating the assay’s effectiveness in reproducing known anti-TB drug effects. Interestingly, auranofin (anti-rheumatic), nitazoxanide (anti-parasitic), and lansoprazole (PPI), drugs with known antimicrobial activity, exhibited significant inhibitory effects against *Mtb* Erdman but were ineffective against *Mm* at 20 µM (**S19 Fig**). Yet, these drugs inhibited both species significantly in the 3D tuberculoma bioplatform. Chlorpromazine hydrochloride (antipsychotic and acid sphingomyelinase inhibitor) and fingolimod hydrochloride (acid sphingosine-1-phosphate receptor modulator), previously reported for direct anti-mycobacterial activity, demonstrated high inhibitory effects in 3D tuberculoma bioplatform, but were ineffective in 3D broth cultures. Other compounds with known direct anti-mycobacterial activity, including N-acetyl-L-cysteine, verapamil, sodium valproate, and irbesartan, were ineffective in both 3D broth culture and the tuberculoma bioplatform. Five compounds previously identified as potential HDT drugs—clementine fumarate, tin protoporphyrin IX, tyrphostin AG 494, metixene hydrochloride, and fluoxetine hydrochloride— exhibited significant growth inhibition of either one or both species in the 3D broth cultures, suggesting that they possess some pathogen-targeting effects. By excluding any direct microbicidal activity in axenic culture, these results indicate that among 12 ‘top-hit’ compounds (**Fig 6B**), AT9283, zuclopenthixol dihydrochloride, dovitinib, 3’4’ dichlorobenzamil HCl, and H89 exert their inhibitory effects on mycobacteria through macrophages in 3D tuberculomas.

### Assessment of compound cytotoxicity in the 3D microenvironment

To verify whether observed effects stem from influences on host-cell viability, we treated uninfected 3D cultures of THP-1 cells or PBMCs with individual compounds and measured cytotoxicity using the CytoTox-Glo^TM^ assay. The results showed strong concordance between the two cell cultures, irrespective of whether immortalized or primary human cells were used. Among the 16 compounds that caused >50% cytotoxicity, 12 were toxic to both cell types at 20 µM (**S20A and S20B Fig**), with 11 having FDA approval or undergone clinical evaluations, indicating that they are safer for human use (**S6 Table**). This highlights a discordance between *in vitro* cytotoxicity and *in vivo* testing results. While toxicity assessments are critical in therapeutic evaluation, the results imply that *in vitro* cytotoxicity methods and doses may not capture the complexities encountered in the human body during *in-vivo* testing. Despite a correlation between cytotoxicity and reduced bacterial loads in the *Mm*-infected THP-1 model (**S20C and S20D Fig**), only eight of the 17 compounds that lowered *Mm* burdens by >50% also reduced THP-1 viability by >50%. Conversely, four compounds that increased *Mm* burdens also exhibited >50% cytotoxicity. To determine the cytotoxic concentration for a 50% reduction in THP-1 cell viability (CC50), we tested a range of concentrations of each cytotoxic compound in 3D cultures of THP-1 cells expressing RFP (**S21 Fig**). Among the 12 ‘top hits’, while six were toxic at 20 µM, effective concentration (EC50) values were obtained for nitazoxanide, doxycycline, dovitinib, and 3’4’ dichlorobenzamil HCl. At nontoxic doses, auranofin reduced *Mm* burdens by <50% and AT9283 did not inhibit *Mm* growth. Given that *Mm* and *Mtb* can grow intracellularly post-macrophage death and *Mm* can quickly overtake the spheroid structure through extracellular proliferation, we inferred that compound-induced cytotoxicity may not necessarily correlate with reductions in intracellular mycobacterial load in *in vitro* assays. We posited that HDT might induce cell death through innate immune mechanisms—such as apoptosis, autophagy, and pyroptosis—which could decrease cellular viability while still inhibiting intracellular pathogen survival. Thus, an ideal bioplatform should characterize these immune mechanisms concurrently with therapeutic efficacy within 3D granulomas, which led us to explore the innate macrophage defenses activated by HDT compounds.

### The readouts of immune mechanisms of HDT can be incorporated *in situ* in the tuberculoma bioplatform

Physiological or pharmacological activation of hypoxia, phagolysosome fusion, autophagy, lysosomal acidification, and inflammasome pathways in infected human macrophages and animal models significantly reduces the survival of intracellular *Mtb* [8, 55, 70, 71]. We explored the incorporation of innate immune readouts into our 3D model of THP-1 monocytes and *Mtb* Erdman (WT), utilizing fluorogenic probes to assess hypoxia and lysosomal acidification. We also used THP-1 reporter cells to monitor autophagic flux and ASC-dependent inflammasomes in the 3D co-cultures, employing high-content imaging to analyze modulation of innate immune responses within 3D tuberculomas by compounds. Despite reducing cell viability and potentially inhibiting reporter-probe activity, we did not exclude cytotoxic drugs in this screen.

Following treatment with six non-cytotoxic ‘top-hit’ compounds at 20 µM (**Fig 7A**), tuberculomas exhibited increased hypoxia in the cores or cellular aggregates (blue arrows) distinct from bacteria-permissive necrotizing lesions (white arrows) on day 6 post-treatment, compared to DMSO or no-drug treatment. Additionally, treated tuberculomas showed cells with increased autophagolysosome formation and acidification, as revealed by increased red over yellow-green fluorescence. While rifampicin-treated tuberculomas exhibited similar hypoxia and autophagy activation features, their scaffolds displayed a less acidic environment, likely due to reduced *Mtb*-induced cell death. In contrast, tuberculomas treated with five representative non-cytotoxic ‘non-hit’ compounds (**Fig 7B**) showed more cellular aggregates with autophagosome accumulation (yellow arrows) despite acidification of tuberculoma cores, indicating a blockade of downstream steps in autophagy in these aggregates. By day 6 post-treatment, these tuberculomas showed less hypoxia staining, mirroring the results of DMSO or untreated controls. Autophagy induction is a dynamic multi-step process, and serial imaging over six days revealed that the autophagy flux progressed relatively swiftly within three days in tuberculomas treated with ‘top-hit’ compounds (**S22A Fig**), especially with lansoprazole and H89. Furthermore, more ASC-GFP-expressing cells persisted on day 6 post-treatment, with reduced *Mtb* growth compared to ‘non-hit’ compound-treated tuberculomas or negative controls (**S22B Fig**), suggesting increased inhibition of *Mtb*-induced host-cell death. In ‘non-hit’ compound-treated tuberculomas, *Mtb* growth occurred unfettered, and autophagy flux developed gradually between three and six days, with several cell clusters lacking autophagosome maturation. In rifampicin-treated tuberculomas, autophagy flux developed slowly by day 6, coinciding with the onset of starvation. Some cellular aggregates lacked autophagosome maturation, suggesting delayed induction of autophagy and incomplete sterilization. Rifampicin-treated tuberculomas contained cellular clusters with *Mtb* persistence and strong inflammasome activation in epithelioid macrophages. Overall, our findings using ’top-hit’ compounds confirm the susceptibility of *Mtb* to pharmacological induction of autophagy and lysosomal acidification in a 3D tuberculoma model.

**Fig. 7.**
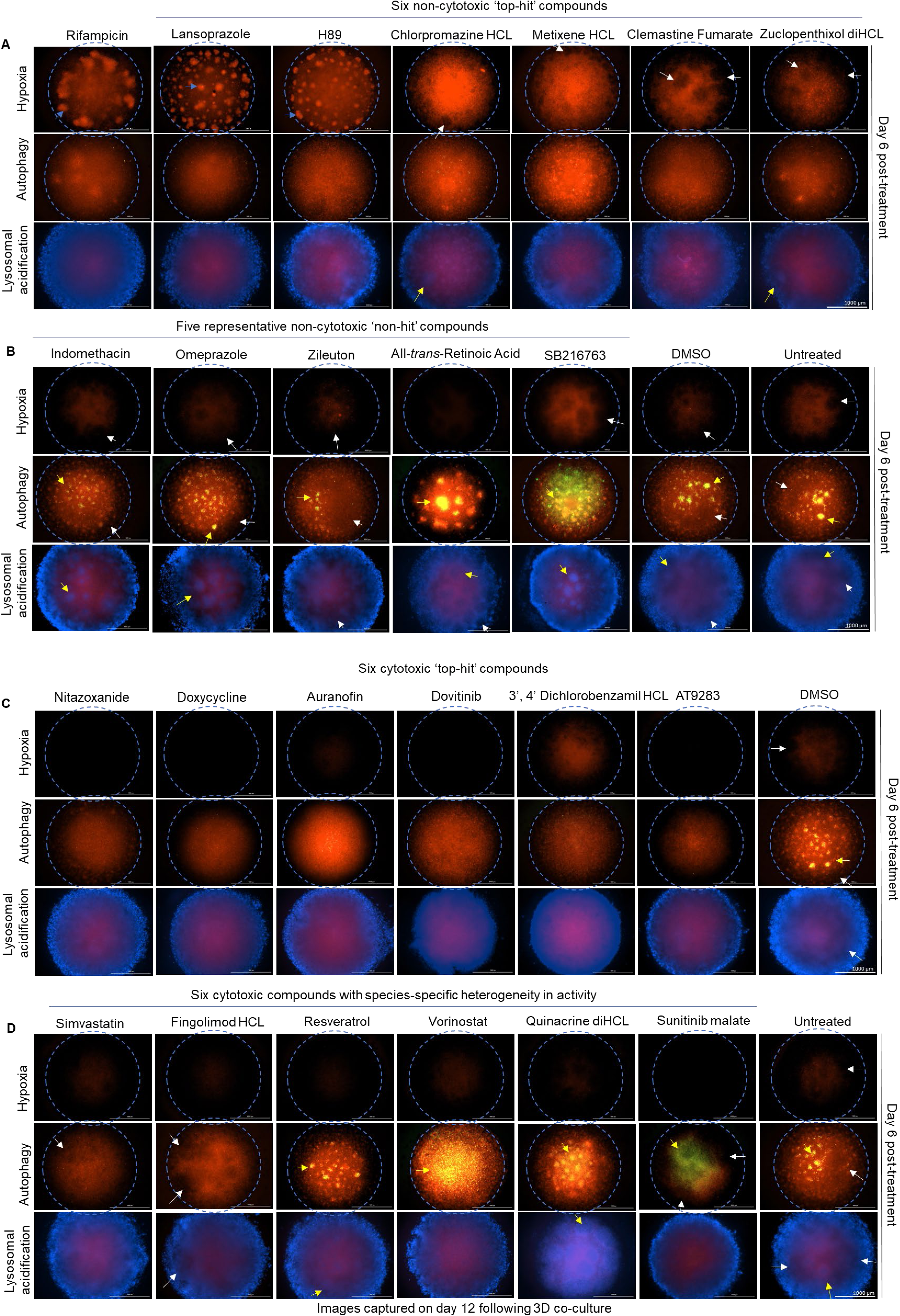
Leading HDT compounds induce crucial innate immune defenses in the 3D tuberculomas. Pharmacological modulation of hypoxia, autophagy, and lysosomal acidification in 3D tuberculomas by (**A**) six non-cytotoxic ‘top-hit’ compounds in comparison with antibiotic rifampicin, (**B**) five representative non-cytotoxic ‘non-hit’ compounds, (**C**) six cytotoxic ‘top-hit’ compounds, and (**D**) six cytotoxic compounds that show *Mycobacterium* species-specific heterogeneity in growth inhibition but reduce 25–50% of *Mtb* Erdman (tdTomato) burdens by day six post-treatment at 20 µM. Hypoxia and lysosomal acidification were investigated using Hypoxia Red and Lyso-ID^®^ Red fluorogenic probes in the Erdman (WT)-infected tuberculomas of THP-1 monocytes. Autophagy was studied in the Erdman (WT)-infected tuberculomas of THP-1 autophagy reporter monocytes. In these cells, the RFP-GFP-tagged LC3 probe emits yellow signals (red plus green) in the cytosol and autophagosomes but only red fluorescence in acidic autophagolysosomes (as acid-sensitive eGFP is quenched or degraded). Following the failure in autophagosome-lysosome fusion, yellow puncta accumulate, while yellow and red puncta are reduced during autophagy inhibition. Images were captured on day 6 post-treatment using Cytation-5. Blue arrows (in **A**) indicate live cellular aggregates with Hypoxia Red staining, distinct from the unstained *Mtb*-permissive lesions (white arrows) in the granuloma zones or cores of tuberculomas. Yellow arrows (in **A**–**D**) indicate cellular aggregates lacking autophagosome maturation (indicated by yellow-green puncta) and lysosomal acidification. These aggregates are readily stained with the nuclear stain Hoechst blue but not with the fluorogenic Lyso-ID^®^ Red dye. (**A**–**D**) The images shown are representative of at least 3–6 tuberculomas treated per compound or control from one of the two experiments that investigated individual innate immune mechanisms, hypoxia, and autophagy, and one experiment that assessed lysosomal acidification.

Tuberculomas treated with six cytotoxic ‘top-hit’ compounds showed less hypoxia on day 6 post-treatment, alongside induction of autophagy and acidification in the centers (**Fig 7C**), likely due to increased cell death. While autophagy flux appeared to progress between one and three days, ASC-GFP-expressing cells decreased, concomitantly inhibiting *Mtb* growth compared to DMSO by day 6 (**S22C** and **S22D Fig**). Similarly, we investigated six remaining cytotoxic compounds that exhibited species-specific heterogeneity in mycobacterial growth inhibition, yet reduced *Mtb* Erdman burden by 25−50% in 3D tuberculomas (**Fig 7D**). Treatment with simvastatin or fingolimod hydrochloride induced autophagy between three and six days (**S22C** and **S22D Fig**). In contrast, histone deacetylase inhibitors, resveratrol and vorinostat, and known autophagy inhibitors, quinacrine dihydrochloride and sunitinib malate, induced incomplete autophagic flux in 3D tuberculomas.

Excessive hypoxia can cause pathogenic necrosis in mycobacterium-infected macrophages, while autophagic and inflammasome activity can lead to non-apoptotic programmed macrophage death and pyroptosis. We examined the results of 65 compounds to understand further the association between these immune processes and *Mtb* control in 3D tuberculomas. When considered individually, no direct association with the inhibition of Erdman growth on day 6 post-treatment was found. For example, KN62, loperamide, and glybenclamide increased hypoxia compared to DMSO (**S23A Fig**), while imatinib, KN62, and loperamide induced autophagy or core acidification but did not significantly inhibit bacterial burdens. Likewise, among nine compounds previously linked to autophagy activation in the *Mtb*-infected 2D cultures (**S23B Fig**), only gefitinib and everolimus induced autophagy with autophagolysosome formation in tuberculomas by day 6. The remaining seven compounds showed incomplete autophagy flux in numerous cellular clusters, highlighting discordance in their ability to induce autophagy in the 3D tuberculoma milieu. None of these nine compounds, except SRT1720, significantly inhibited *Mtb* Erdman loads. Notably, autophagy flux was delayed in 3D tuberculomas treated with these compounds (**S23C**−**F Fig**). These results indicate that the rapid pharmacological induction of complete autophagy flux, rather than mere autophagosome formation, correlates with the inhibition of *Mtb* growth in 3D tuberculomas, highlighting the versatility of the bioplatform in assessing innate immune defenses induced by therapeutics *in situ*.

### HDT compounds can be classified into functional clusters based on therapeutic effects and immune mechanisms in the 3D bioplatform

Several compounds effectively reduced bacterial burdens and lesions across multiple mycobacterial-infection models (**S7 Table**). The six ‘top-hit’ compounds—nitazoxanide, auranofin, doxycycline, dovitinib, 3’4’ dichlorobenzamil HCl, and AT9283—demonstrated significant cytotoxicity at 20 µM in both THP-1 and PBMC cultures. These compounds reduced bacterial burdens by >50% compared to controls in all infection models, except for AT9283, which achieved a 42.2% reduction (*p* = 0.0045) of *Mtb* Erdman burdens in the PBMC model. Classified as Cluster-1, these compounds also effectively reduced lesion burdens by >50% in three out of four infection models using THP-1 monocytes. They induced rapid autophagy flux with lysosomal acidification but did not induce hypoxia at a cytotoxic concentration (**Fig 8A**). By using various concentrations, we confirmed that these compounds induce autophagy at their CC50 or EC50 amounts in THP-1 monocytes transfected with the autophagy sensor LC3B-RFP (BacMam 2.0) and infected with *Mm* (WT) (**S24 Fig**). As the concentrations of auranofin and AT9283 decreased, we observed a decline in autophagy flux and an increase in autophagosome accumulation in 3D tuberculomas of THP-1-Difluo autophagy reporter cells (**S25 Fig**). The reduced autophagy flux aligned with decreased inhibition of Erdman burdens in the 3D tuberculomas of ASC-GFP inflammasome reporter cells. Notably, cellular viability improved at lower compound concentrations, with no corresponding increase in hypoxia induction. This confirms that the compound’s effectiveness correlates with the induction of autophagy rather than hypoxia in 3D tuberculomas.

**Fig. 8.**
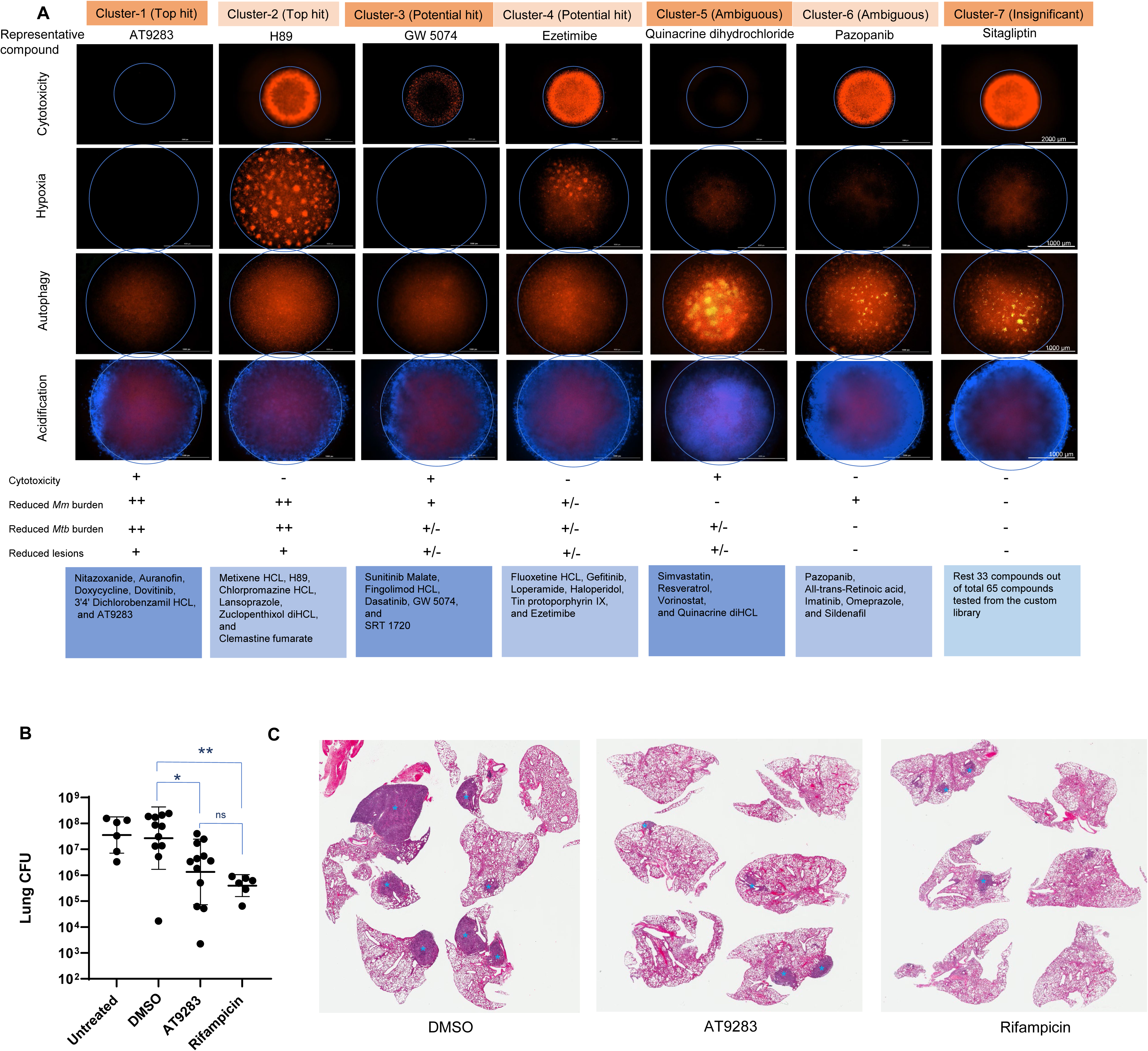
HDT compounds can be classified into functional clusters, and AT9283 enhances host resistance to *Mtb* in aerosol-infected mice. (**A**) The HDT compounds can be classified into seven distinct functional clusters based on the immune mechanisms and therapeutic efficacy in the 3D tuberculoma bioplatform. The key innate immune mechanisms, including hypoxia, autophagy, and lysosomal acidification, induced by representative compounds from seven clusters along with their cytotoxicity profiles, are shown. Compounds belonging to these clusters are listed. Images represent one of the two experiments that assessed cytotoxicity in the THP-1-RFP 3D spheroids, and hypoxia, autophagy, and lysosomal acidification in 3D tuberculomas. n = 3–6 spheroids per compound or controls in each experiment. (**B–C**) The therapeutic effect of AT9283 in C3Heb/FeJ mice infected with *Mtb* Erdman via the aerosol route with 10–15 CFU. Seven days after infection, mice received 16 doses of treatment by oral gavage, as described in the Results and Methods. (**B**) AT9283 treatment inhibits bacterial growth in the lungs of *Mtb*-infected mice. Data from AT9283 treatment compared to DMSO controls from two independent experiments (n = 6 mice/group) are shown. \**p* < 0.05, \*\**p* < 0.01 compared to DMSO using Kruskal-Wallis with Dunn’s post-hoc test. Error bars are the standard deviation, and the horizontal line indicates the Geometric Mean. (**C**) AT9283 treatment decreases granuloma burden in the lungs of *Mtb*-infected mice. Representative lung lobe images are shown (n = 6 mice/group). Blue stars denote granuloma lesions.

The six ‘top-hit’ compounds in Cluster-2—metixene HCl, H89, chlorpromazine HCl, lansoprazole, zuclopenthixol diHCl, and clemastine fumarate—demonstrated <50% toxicity in THP-1 cultures. They effectively reduced bacterial burdens by >50% compared to untreated controls in all mycobacterial-infection models, except for metixene HCl, zuclopenthixol diHCl, and clemastine fumarate, which showed no significant inhibition of Erdman burdens in the PBMC model. Interestingly, lansoprazole, an FDA-approved drug, induced >50% toxicity in PBMC cultures. These six compounds also reduced >50% of lesion burdens in three infection models using THP-1 and induced hypoxia, rapid autophagy flux, and lysosomal acidification (**Fig 8A**). Notably, with decreasing concentrations of lansoprazole and H89, hypoxia, autophagy, and ASC-speck formation decreased by day 6, while Erdman burdens increased. This suggests that these pathways contribute to the efficacy of these compounds in 3D tuberculomas (**S25 Fig**).

Cluster-3 comprised five potential ‘hit’ cytotoxic compounds—sunitinib malate, fingolimod HCl, dasatinib, GW 5074, and SRT 1720—which reduced bacterial burdens by >25% compared to untreated controls in all infection models, although dasatinib did not inhibit Erdman burdens in the THP-1 and PBMC models. They also inhibited >25% of lesion burdens in at least three infection models using THP-1 and induced autophagy flux, albeit slowly, between three and six days, along with lysosomal acidification, but not hypoxia (**Fig 8A**). Cluster-4 included five potentially ‘hit’ noncytotoxic compounds—fluoxetine HCl, gefitinib, loperamide, haloperidol, tin protoporphyrin IX, and ezetimibe—which inhibited bacterial burdens by >25% in at least three infection models using THP-1. However, only fluoxetine and tin protoporphyrin IX reduced Erdman burdens by >25% in the PBMC model. Furthermore, they demonstrated heterogeneous efficacy in reducing lesion burdens and induced autophagy flux slowly between three and six days, with hypoxia and lysosomal acidification.

The comparisons revealed species-specific heterogeneity in bacterial growth inhibition by HDT compounds (**Fig 8A** and **S7 Table**). The four cytotoxic compounds—simvastatin, vorinostat, resveratrol, and quinacrine diHCl—which formed Cluster-5, did not decrease *Mm* burdens but inhibited *Mtb* burdens by >25% in all four or at least one of the four *Mtb*-infection models of THP1 and PBMCs. Conversely, Cluster-6 compounds—pazopanib, all-trans retinoic acid, imatinib, omeprazole, and sildenafil—were effective only in the *Mm*-infection model using this criterion. Compounds in these two clusters also exhibited heterogeneity in their ability to inhibit lesion burdens and induce innate defenses in the THP-1 infection models. These results highlight subtle differences in human macrophage responses to *Mm* and *Mtb* infections.

### AT9283 effectively inhibits *Mtb* burdens and granuloma lesions in mice

One of the ‘top-hit’ compounds and a multi-targeted kinase inhibitor, AT9283, presently in clinical trials for treating malignancies [36, 37], significantly inhibited *Mm* and *Mtb* burdens and granuloma lesions in the THP-1 model. It exhibited anti-*Mtb* activity preferentially in 3D tuberculomas compared to traditional 2D cultures in a host-directed manner without apparent direct anti-mycobacterial activity in axenic 3D cultures. To determine the efficacy of AT9283 against *Mtb in vivo*, we investigated its anti-mycobacterial effects in C3Heb/FeJ mice, targeting the period before the onset of peak adaptive immunity, given that selection was performed in a macrophage-centric 3D model. Mice were infected with *Mtb* Erdman via the aerosol route. Seven days post-infection, groups of mice were treated with AT9283 (35 mg/kg/day), rifampicin (10 mg/kg/day), no drug, or DMSO (<5% v/v) in 150 µl of sterile water, five days per week, for three weeks. On day 28 post-infection, mice were euthanized, and the left lungs were homogenized for CFU analysis while the right middle lung lobes were processed for histopathological evaluation. Mice treated with AT9283 had significantly lower bacterial loads (p = 0.04, average 1.1-log reduction) compared to the DMSO controls (**Fig 8B**). Likewise, AT9283-treated mice exhibited no gross signs of toxicity and had fewer granuloma lesions (**Fig 8C**). These results support the potential of AT9283 for treating early *Mtb* infection. Further research is required to assess its efficacy as an adjunct to standard anti-TB antibiotic therapy for chronic infections and TB disease.

## Discussion

We developed a facile bioplatform that utilizes human cells and pathogenic mycobacteria for high-content analysis of 3D *in vitro* tuberculomas, enabling the screening of anti-TB drugs, including HDT compounds. Although our system lacks the complexities of lung tissues, it forms tuberculoma-like structures and crucial milieus through cellular self-assembly, rather than relying on artificial confinement, commonly employed by contemporary models [25–29, 72]. The formation of tuberculoma-imitative structures is contingent upon specific spatiotemporal factors, including cell-culture-ware, microwell geometry, surface chemistry, culture conditions, and timeline. Despite using 3D spheroid technology and primary or immortalized macrophages with mycobacteria, tuberculomas with organized lesions or cavitary features were not developed in existing models, even though infected 3D spheroid structures exhibited traits, such as central hypoxia and necrosis, shared by control spheroids [28, 29, 73].

We validated the bioplatform by demonstrating that (i) organized collection of granulomas rather than mere macrophage aggregates occur, (ii) classic solid, necrotic, and cavitary forms develop, (iii) crucial features, pathways, and molecular expression analogous to human tuberculomas arise, (iv) the occurrences in the model have parallels in the natural hosts, and (v) known antimicrobials are reliably effective within this system. Our assays establish the effectiveness of test compounds within 14 days by quantifying both bacterial burdens and granuloma lesions *in situ*, offering a more expedited and efficient system for screening extensive compound libraries in tuberculoma milieus for primary ‘hit’ identification and validation. The shortlist of ‘top hits’ from this system can be investigated in mammalian models for lead generation. The bioplatform is highly flexible in terms of cellular composition and allows for scalability in fully automated HTS applications, utilizing standard liquid handlers. It permits prolonged co-cultures for over 50 days (the longest attempted) by replenishing the medium and immortalized monocytes, allowing relatively long-term investigations.

While human primary cells are preferred for generating *in-vitro* granuloma models [30], challenges associated with scalability, reproducibility, and cost during HTS campaigns have prompted the use of immortalized cells [38–40, 66]. Stem cells can generate large quantities of macrophages [74, 75]. However, the self-renewing and proliferative capabilities of macrophages post-differentiation will be essential for maintaining granuloma structures in prolonged experiments. Without self-renewal, repeated incorporation will be necessary for facilitating macrophage influx into developing granulomatous aggregates. Recent hPSC-derived lung organoid models faced challenges in generating adequate macrophages, necessitating the microinjections of THP-1-derived macrophages or co-cultures of induced macrophages with alveolar epithelial organoids [33, 76]. Additionally, the impacts of growth factors used for hPSC-derived macrophage differentiation and maintenance [75] on the efficacy of test HDT are unclear. Cancer cell lines, such as THP-1, are generally not considered because they are deemed to diverge from their primary counterparts and create immunoregulatory milieus that can influence *Mtb* interactions [30]. Despite concerns about their transformed origin, 3D tuberculomas of THP-1 monocytes expressed immunoregulatory markers (e.g., PD-L1, CTLA-4, galectin-9, IDO1, and others) that are also found in human TB lesions and tumors [48]. These similarities suggest that mechanistic overlaps between TB granulomas and tumor microenvironments could be exploited to repurpose cancer immunotherapies for TB. A strong concordance in compound effectiveness or cytotoxicity between the THP-1 and PBMC models in our study indicates that THP-1-derived macrophages, with their self-renewal and recruitment capabilities, can serve as a viable alternative, promoting throughput and reproducibility in early screenings. The 3D cell culture framework provided here, along with streamlined methods for generating hPSC-derived macrophages and multilineage immune cells in the lung tissue [76–78], can further advance the tuberculoma platform.

Many findings in human hosts and animal models are reflected in our model. For instance, granulomas with epithelioid macrophage transformations and cell adhesions were formed [46], illustrated by the increased expression of cadherins, catenins, ICAMs, CEACAMs, claudins, and desmosomal markers. Furthermore, the model demonstrated upregulation of angiogenesis and vascularization markers (e.g., VEGF, VEGFR, VCAM, epiregulin, and angiopoietin), and classic tuberculoma characteristics, including increased hypoxia, necrosis, MMP activity, and matrix remodeling. In pulmonary TB, granuloma heterogeneity, including non-necrotizing leukocyte aggregates and necrotizing forms, has been described [20, 43, 49]. Our model reflected such diversity with bacterial growth-permissive and restrictive granulomas and showed overexpression of previously reported biomarkers, such as IP-10, HO-1, MMP-9, albumin, and RBP, among others [60, 79]. Furthermore, the pathogenic mycobacterial loads corresponded with granuloma formation, and attenuated BCG strains, which lack the ESX-1/RD1 virulence locus, formed smaller granulomas, confirming that granulomas promote mycobacterial growth and that the virulence determinant is essential for effective macrophage recruitment and granuloma development [11, 45]. The observations that TNF blockade at the time of *Mtb* infection or granuloma formation disrupts the structural integrity of nascent granulomas provided validation that TNF is critical for granuloma maintenance. Consistent with the *in vivo* findings [56, 58, 59], TNF blockade in the *Mtb*-infected model did not prevent granuloma formation. In the *Mm*-infected model, it resulted in miliary granuloma formation, associated with pulmonary and extrapulmonary dissemination. These observations reinforce that TNF is not required for granuloma formation but rather preserves granuloma integrity. Many observed features are not invariably replicated in artificially enclosed 3D models [25–27, 72].

Numerous findings in our model provided new insights that merit further *in vivo* validation. While virulent *Mtb* can inhibit autophagy, our findings suggest that this occurs primarily in permissive granuloma macrophages. The cavitary transformation observed suggests that excessive ECM deposition beyond the threshold limit in the tuberculomas, along with increased *Mtb* and granuloma burdens, likely contribute to cavity formation and present a new premise for validation in animal models. Our model thus offers a unique tool for investigating the drivers of cavity formation and identifying therapeutics that modulate cavitation. In recent years, several biotherapeutics have become available for treating various immune-mediated disorders, and our platform identifies biologics that could either mitigate or exacerbate the risk of TB progression. For example, humanized mAb biosimilars targeting α4β7, CD30, and IGF1R delayed granuloma expansion and reduced bacterial burdens. Interestingly, while CD30/TNFRSF8 co-stimulation supports the development of protective T cells [80], our findings suggest that it also regulates *Mtb* growth in myeloid cells independently of T cells in nascent granulomas. Conversely, biosimilar targeting CD52 on myeloid cells increased granuloma size and bacterial load, reflecting the heightened TB incidence observed after alemtuzumab treatment [81], which is thought to be due to rapid depletion of lymphocytes. LFA-1 (CD11a/CD18) is required for protective immunity in mice infected with *Mtb* [82]. In our model, efalizumab biosimilar targeting CD11a on myeloid cells disrupted the structural integrity of granulomas. Additionally, blocking VEGF-VEGFR signaling in myeloid cells, without endothelial involvement, delayed granuloma expansion and reduced *Mtb* burden. Anti-VEGF treatments in animal models have been shown to enhance small molecule delivery, limit mycobacterial growth, and improve host survival [51, 83]. These findings highlight the critical role of innate immunity in granuloma processes, often occurring independently of adaptive immunity or non-hematopoietic cells.

HDTs have the potential to clear persistent and antibiotic-tolerant bacteria, improving the treatment of both drug-susceptible and drug-resistant TB by enhancing the antimicrobial activities of phagocytes and reversing *Mtb*-imposed immune deficits [7–9]. Crucially, our platform demonstrated that many HDT compounds effective in treating early infection in 2D macrophage screens were ineffective once infection progressed and granulomas formed. Many autophagy-inducing drugs failed to activate autophagy within these granulomas. Of the nine previously reported lead compounds in 2D cell cultures [39], only four (dovitinib, H89, 3’4’ dichlorobenzamil HCl, and GW5074) consistently inhibited bacterial loads across our infection models. Drugs like imatinib, pazopanib, and metformin, despite showing promise in reducing *Mm* and *Mtb* burdens in 2D cell-culture or animal models when administered before or during early infection [51, 84–86], were ineffective in *Mtb*-infected 3D tuberculomas of primary or immortalized monocytes. Although our results confirm the effectiveness of imatinib and pazopanib in reducing *Mm* burdens and granuloma lesions by 25−50%, they recommend additional studies in *Mtb*-infected animal models post-granuloma establishment. Notably, imatinib treatment has been shown to increase the risk of TB in persons with cancer [87], and therapeutic concentration of metformin has been found to increase *Mtb* burdens in non-diabetic mice compared to diabetic mice [88], emphasizing context-specific drug evaluation. Nonetheless, our screens identified 12 compounds that consistently inhibited mycobacterial loads by >50% across all infection models, despite the heterogeneity of mycobacterial strains in recruiting host factors for survival. Six of these (AT9283, 3’4’ dichlorobenzamil HCl, dovitinib, H89, and zuclopenthixol dihydrochloride) acted in a host-directed manner without direct antimycobacterial activity. Known HDT compounds, previously identified through animal studies [66–68] and those with both host-directed and pathogen-targeted effects in *Mtb*-infected humans as standalone or adjunct therapies [10, 69, 89], were also effective, validating the platform’s ability to identify clinically relevant candidates. Multi-kinase inhibitors AT9283, dovitinib, and H89 have been evaluated in clinical trials for cancer treatments, while zuclopenthixol is a dopamine antagonist for schizophrenia and could be repurposed for TB treatment. H89 inhibits AKT1, and AT9283 and dovitinib are predicted to target macrophage RTK signaling, thereby controlling intracellular *Mtb* [39]. Importantly, our study demonstrated that AT9283 is effective in the murine TB model. Building on this proof-of-concept study, we have successfully screened two extensive compound libraries (n = 2558) in the THP-1 bioplatform and identified several promising candidates (unpublished work).

Compared to 3D granuloma models, which rely on specialized instrumentation and employ ECM embedding, microencapsulation, or magnetic levitation [25–29], our system is simpler, scalable, and more amenable to HTS. The platform provides comprehensive information on the effectiveness of drugs, including bacterial-burden reduction, granuloma (pathology) resolution, cytotoxicity, and immune mechanisms *in situ*. It has the potential to reduce animal use, offering both ethical and economic benefits. Beyond compound screening, the platform using human primary cells can be adapted for diverse applications, including personalized immunotherapy development, predicting vaccine effectiveness *ex vivo* in clinical trials, studying TB co-infections and co-morbidities, and investigating a wide range of granulomatous diseases, including foreign-body, non-infectious, and infectious granulomas caused by diverse agents. It can accommodate fluorescence-based diverse readouts and integrate solvatochromic dyes [90] for labeling mutant strains or clinical isolates. Moreover, the platform can be mass-produced and cryopreserved for commercial or translational use [91].

Our model and study have limitations. While this model replicates critical features of 3D tuberculomas, it does not capture the full morphological and cellular diversity of human lesions. Tubercles in persons with TB range from a few millimeters to several centimeters [16–18, 92], and larger tuberculomas could be generated by scaling up cell and bacterial inputs to better model drug penetration and diffusion gradients. The absence of accessory cell types, such as fibroblasts, epithelial cells, endothelial cells, and neutrophil subsets, may limit specific applications, though these can be easily incorporated given the system’s tractability. High-end imaging systems used in our study may not be universally accessible; however, the platform is compatible with plate fluorometers for rapid drug screening. The absolute biological activity assessed using fixed drug dosing in our platform does not account for pharmacological variability seen in patients, which could be addressed in future studies using microfluidic-based platforms. Although our study identified several promising HDT candidates that activate innate immune mechanisms, more in-depth evaluations are needed to describe specific pathways activated and understand the host-directed effects. Finally, all candidates identified in the platform require further validation in animal models and clinical studies. Toward this goal, we are currently testing several ‘hits’ in murine TB models.

## Materials and methods

### Ethics statement

This activity was reviewed by Centers for Disease Control and Prevention (CDC), Atlanta, and Emory University, Atlanta, Georgia, USA, and was conducted consistent with applicable federal law, 45 Code of Federal Regulations, Part 46. The peripheral blood was drawn from healthy human participants after obtaining written consent (CDC Institutional Review Board [IRB] Protocol Number 1652; Emory University IRB Protocol Number 00045947). CDC reviewed the use of peripheral blood for this activity as Project Number 7294. All animal experiments were performed strictly following the U.S. Public Health Service Policy on the Humane Care and Use of Animals and the Guide for the Humane Care and Use of Laboratory Animals. The Institutional Animal Care and Use Committee, CDC, approved the animal protocol (SABMOUC3110).

### Reagents

Critical reagents and resources used in the study are listed in the **S8 Table**. Biosimilars were purchased from R&D Systems, and biologics and control antibodies were obtained from Sigma-Aldrich. The custom-built library of known potential HDT compounds prepared as a 10 mM solution in DMSO was purchased from the Small Molecule Screening Library Service, Aldrich Market Select, Millipore Sigma, USA (**S6 Table**).

### Mycobacterial culture

Mycobacteria were cultured in Difco Middlebrook 7H9 medium (Becton Dickinson) supplemented with 10% ADC (Becton Dickinson), 0.05% Tween-80, and 0.4% glycerol (Sigma-Aldrich) at 30°C (*Mm*) or 37°C (*Mtb* and BCG). The mycobacterial strains and the antibiotics used in 7H9 broth for recombinant mycobacterial selection are listed in the **S8 Table**. The frozen stocks were stored at -80°C.

### Cell culture

Immortalized THP-1, U937, AMJ2, RAW 264.7, and J774A.1 cell lines (American Type Culture Collection) and THP-1-RFP, THP-1-GFP (Applied Biological Materials, Inc), THP-1-ASC-GFP, and THP-1-Difluo-hLC3 (InvivoGen) reporter cell lines were grown and maintained as per the supplier’s instructions (**S8 Table**). Venous blood (250 ml) was collected in BD Vacutainer Cell Preparation Tubes from healthy human donors, and PBMCs were isolated by centrifugation at 1500−1800g for 30 minutes. Primary CD14^+^ monocytes were purified from PBMCs by CD14 Magnetic-Activated Cell-Sorting (MACS) Microbeads Technology using LS columns (Miltenyi Biotec). Unlabeled PBMCs, which contained lymphocytes and mononuclear cells other than CD14^+^ monocytes, were collected as CD14^−^ cells. Immortalized or human primary cells were grown or maintained in Gibco RPMI 1640 medium containing L-glutamine (1mM) and supplemented with heat-inactivated FBS (9.63%), 100 mM sodium pyruvate (0.87%), 1M HEPES (0.87%), and 10,000 units/ml penicillin-streptomycin solution (1%) (Life Technologies-Invitrogen) before use in co-cultures. See Supplementary Methods for the growth of immortalized cell lines.

### 3D ‘mycobacteria-in-spheroid’ co-culture

Mycobacterial frozen stocks were thawed, washed with complete RPMI 1640 medium without antibiotics (co-culture medium), and pelleted by centrifugation at 5000 g. The mycobacterial suspension was passed through a 25*–*27G needle fitted to a syringe 15 times to break bacterial clumps. The suspension was diluted in the co-culture medium to achieve an MOI of 0.008 for *Mm* strains and 0.025*–*0.05 for *Mtb* strains (unless otherwise indicated). The actual CFU count in the mycobacterial suspension was ascertained by plating dilutions onto Middlebrook 7H10 agar plates. Cell cultures were washed with a prewarmed (37°C) co-culture medium to remove antibiotics from the growth medium and then pelleted by centrifugation at 250 g. A total of 10^5^ immortalized monocytes-macrophages, reporter cells, or human PBMCs were seeded in 100 µl co-culture medium per well in 96-well 3D cell culture plates (**S8 Table**) and inoculated with 100 µl of a mycobacterial suspension. Cells were mixed by pipetting without touching the bottom of the microwell. Microplates were incubated at 37°C, 5% CO2, and 100% humidity to continue the co-culture until readouts were obtained using Cytation 5 or a CFU assay. For 3D co-cultures using purified PBMC subsets, 2×10^5^ CD14^+^ monocytes were infected with *Mtb* Erdman (MOI 0.025) in a 96-well 3D cell culture plate (Corning). The cell suspension (200 µl/well) was mixed by pipetting before incubation. On day three post-co-culture, 100 µl of cell suspension containing 4×10^5^ autologous lymphocyte-rich CD14^−^ PBMC subsets in the co-culture medium was added to each microwell without disturbing the monocyte-macrophage spheroids. In the workflow incorporating ECM, 3D co-cultures of THP-1 monocytes and *Mm* 1218 GFP were generated as described above, and 5−50 µl of the ECM solution was carefully added to each microwell on day 3. The preparation of the ECM solution is described in Supplementary Methods. The protocol for generating 3D tuberculoma bioplatform workflows is detailed in the Methods article [91].

### Biologic or chemical compound treatment

As described above, co-cultures of THP-1 monocytes, reporter cells, or purified PBMC subsets with *Mm* or *Mtb* strains were developed in 96-well 2D or 3D cell culture plates. Biologics, biosimilars, and control antibodies were used at the indicated doses for treatment immediately before infection or on day 6 post-co-culture. The co-cultures (200 µl/well) were treated on day 6 with chemical compounds at the indicated doses, DMSO at equal v/v in the medium, or medium alone. All exposures were performed in an additional 50 µl fresh co-culture medium per well. No media changes were performed unless indicated otherwise. Mycobacterial growth in co-cultures was monitored longitudinally in real time using fluorescence (Cytation-5 Cell Imaging Multimode Reader, Agilent-BioTek). The fluorescence readings were taken on days 12 and 14 post-co-culture, and images were captured.

### Mycobacterial 3D broth culture

*Mm* or *Mtb* (tdTomato) culture (OD 600 nm, 0.05–0.06) was plated in a 100 µl volume of 7H9 broth containing hygromycin (50 µg/ml) in a 96-well U-bottom, ULA 3D cell culture microplate (Corning) containing 100 µl of fresh 7H9 broth per microwell. The plates were incubated at 30°C or 37°C for *Mm* and *Mtb* in an incubator containing 5% CO2 and 100% humidity, resulting in 3D bacterial growth in microwells. Cultures were treated after 24 hours with compounds in an additional 50 µl of 7H9 broth containing hygromycin. Fluorescence was measured using Cytation-5 on day 6 post-treatment.

### Cell toxicity assays

The CytoTox-Glo^TM^ Cytotoxicity Assay (Promega) measures dead cells in cultures by detecting protease activity from cells with compromised membrane integrity using a luminogenic substrate (AAF-amino luciferin). 3D cell cultures were treated with compounds on day 6, and cytotoxicity was measured on day 12. Spheroids were dissociated to obtain a single-cell suspension in the medium. To obtain the denominator, total cell death was induced by digitonin in equal numbers of cells in the wells. After adding the cytotoxicity assay reagent, luminescence was measured using Cytation-5. As an alternative measure of cytotoxicity, 3D cell cultures of THP-RFP cells were treated with chemical compounds and analyzed for fluorescence reduction (excitation 588 and emission 633nm), indicating cell death. Cytotoxicity (%) caused by the test compounds was calculated relative to cultures treated with DMSO or medium alone. See Supplementary Methods for additional details.

### Immunofluorescence

Whole-mount 3D spheroid immunostaining was performed *in situ* in cell culture plates. The medium was carefully aspirated, and the spheroids were washed twice by carefully adding 75 µl of PBS and incubating for 10 minutes. After aspirating the PBS, spheroids were fixed using 4% paraformaldehyde (75 µl) for 60 minutes. PFA was removed, 0.05% Triton X-100 in PBS (75 µl) was added for 30–60 minutes for permeabilization, and spheroids were washed once with PBS. The spheroids were incubated at room temperature for 1–2 hours with a blocking buffer containing 1% BSA and 5% FBS in PBS. The blocking buffer was removed, and fluorescently labeled antibodies diluted in the blocking buffer (100 µl) were added and incubated overnight at 4°C. Antibodies were used according to the manufacturer’s specifications. Optimal staining concentration was determined by titration (**S8 Table**). Spheroids were washed three times with PBS, counterstained with Hoechst 33342 (1µl/ml), and imaged using a Cytation-5. Samples were stored in the dark at 4°C until microscopic analysis.

### Fluorescent probes and microscopy

Hypoxia, lysosomal acidification, apoptosis, necrosis, biofilm formation, and live and dead cells in 3D cell cultures were probed using commercial probes (**S8 Table**) according to the manufacturer’s instructions and with some modifications after titrations. See Supplementary Methods for details.

### Flow cytometry

Multicolor flow cytometry was conducted on single-cell suspensions of cell cultures. Cell viability was assessed using the trypan blue exclusion method. Two million cells were incubated with an Fc receptor-blocking solution (human TruStain FcXTM, BioLegend) for 15 minutes, followed by staining with a live/dead fixable stain and a pre-titrated anti-human cell-surface CD antibody cocktail. CD68 staining was performed intracellularly. The fluorophore-labeled antibodies and the amount used for staining are provided in the **S8 Table** and the Supplementary Methods. Cells were fixed with 0.25% paraformaldehyde and analyzed on a BD LSRFortessa^TM^ flow cytometer using FlowJo v10.10 software, with a minimum of 55,000 events analyzed. Single-color UltraComp eBeads^TM^ (Invitrogen) and fluorescence minus one (FMO)-stained cells were used as positive and negative controls, respectively.

### Protein array

A Proteomics array was performed using the Quantibody^®^ Human Kiloplex Quantitative Proteomics Array (RayBiotech) to measure the concentrations of 1000 proteins in the cell culture supernatants and lysates of the 3D cell cultures generated with or without ECM (30 µl). Culture supernatants and cells were isolated on day 16 from the control and *Mm* 1218-infected THP-1 co-cultures. Cell lysates were prepared using the lysis buffer supplemented with a protease inhibitor cocktail (RayBiotech), and protein concentrations in lysates were determined using the BCA Protein Assay kit (Sigma-Aldrich). Background levels in blank media control were subtracted from the supernatant samples to calculate array protein concentrations.

### ELISPOT assay

ELISPOT assays were performed using human IP-10/CXCL-10, IL-8/CXCL-8, MMP-1, and MMP-9 ELISpot Development Modules (R&D Systems) and TNF-α ELISPOT pair (BD Biosciences) following the manufacturer’s protocol. See Supplementary Methods for details.

### RNA sequencing

Control and infected 3D spheroids (n = 30/time point) were dissociated to obtain single-cell suspensions. Cells were pelleted and resuspended in 1 ml RNA Protect Cell Reagent (Qiagen). Total RNA was extracted using the RNeasy Plus Mini Kit (Qiagen), and integrity was assessed using RNA Screen-Tape and 4200 TapeStation (Agilent); samples with an RNA integrity number >8.0 were used. RNA was stored at −80°C, and its concentration was determined using the Qubit^®^ RNA BR Assay Kit (Invitrogen) and a Qubit fluorometer. RNA sequencing libraries were prepared with the TruSeq Stranded mRNA Library Preparation Kit and indexed using TruSeq RNA UD Indexes (Illumina). Library quality was evaluated using the D1000 Screen-Tape (Agilent) and quantified using the Qubit^®^ DNA BR Assay Kit (Invitrogen). The sequencing was performed on a NextSeq 500 platform using the 75 × 2-cycle Medium Output Kit (Illumina), yielding about 28 million paired-end reads per sample. RNA-seq analyses are detailed in Supplementary Methods, and data are deposited in the National Center for Biotechnology Information (NCBI) Gene Expression Omnibus (GEO, record number GSE269193).

### Autophagy and inflammasome assays

3D co-cultures of *Mtb* Erdman and THP-1-Difluo (RFP-GFP) hLC3 autophagy or THP-1-ASC-GFP inflammasome reporter cells (InvivoGen) were generated in 3D cell culture plates. Autophagy and inflammasome induction in 3D tuberculomas treated with or without chemical compounds were investigated using longitudinal imaging in a Cytation-5. See Supplementary Methods for additional details.

### Mouse infection

Female C3Heb/FeJ mice (6−8 weeks old) were infected in a Glas-Col aerosol chamber. Six mice were euthanized 1-day post-infection for CFU analysis of bacteria deposited in the lungs. In the remaining mice, the infection was allowed to progress for 7 days before treatment initiation. Mice were subsequently administered an oral gavage of AT9283, rifampicin, or DMSO in sterile water five days per week. Fresh dilutions of the drugs, prepared daily in distilled water from frozen stocks (dissolved in DMSO), were used for oral gavage. On day 28 post-infection, six mice from each group were euthanized, and the left lungs were homogenized and plated for CFUs. CFUs were enumerated 4 and 8 weeks after incubation. The right lung lobes were fixed in 4% paraformaldehyde in PBS, followed by 80% ethanol. Tissues were processed, embedded in paraffin, sectioned, and stained with hematoxylin and eosin or anti-mycobacterial antibodies.

### Statistical analysis

The statistical analyses included Welch’s t-test, 2-way ANOVA (with Tukey’s or Šidák’s post-hoc tests), Brown-Forsythe and Welch ANOVA, and the Kruskal-Wallis test with Dunn’s post-hoc test. Details about technical and biological replicates, experiment numbers, and post-hoc tests are provided in the figure legends. GraphPad Prism software (v9) was used for analyses, with *p*-values < 0.05 considered significant. Further statistical methods for the compound screening assay and RNA-seq are described in the Supplementary Methods.

## Supporting information

Supplemental Methods, Supplementary Figures and Legends for Supplementary Tables

## Acknowledgments

We thank David Tobin, Duke University, for providing the *Mm* ‘M’ wildtype and tdTomato strains, and the CDC Comparative Medicine Branch staff and veterinarians for help with animal husbandry and oral gavage. We acknowledge the Applied Research Team at the Laboratory Branch, Division of Tuberculosis Elimination, for assistance with CFU enumeration, and Jennifer Oosthuizen, Creative Services, CDC, for creating the bioplatform workflow schematic.

The findings and conclusions in this publication are those of the authors and do not necessarily represent the official position of the CDC. References in this manuscript to any specific commercial products, process, service, manufacturer, or company do not constitute their endorsement or recommendation by the U.S. Government or CDC.

## Author contributions

Conceptualization: SBS. Methodology: SBS, WL, AK, SG, VV, and JEP. Investigation: SBS, WL, AK, and SG. Data analyses and graphs/display items: SBS, WL, AK, SG, and VV. Project supervision: SBS and JEP. Writing—Original draft: SBS and WL. Writing—Review and editing: All authors.

## Data and materials availability

All data necessary for evaluating the paper’s conclusions are present in the Main Text and Supplementary Materials. RNA-Seq data can be found in the NCBI GEO database under record number GSE269193. CDC can provide recombinant *Mtb* strains upon scientific review and completion of a material transfer agreement. Requests for the strains should be submitted to the corresponding author.

## References

1. World Health Organization, Global Tuberculosis Report 2024. Geneva: WHO (2024).

2. Trajman A, Campbell JR, Kunor T, Ruslami R, Amanullah F, Behr MA, et al. Tuberculosis. Lancet. 2025;405(10481):850–66. doi: 10.1016/S0140-6736(24)02479-6. PubMed PMID: 40057344.

3. Dartois VA, Rubin EJ. Anti-tuberculosis treatment strategies and drug development: challenges and priorities. Nat Rev Microbiol. 2022. Epub 20220427. doi: 10.1038/s41579-022-00731-y. PubMed PMID: 35478222; PubMed Central PMCID: PMCPMC9045034.

4. MD JL, Boshoff HI, Barry CE, 3rd. The present state of the tuberculosis drug development pipeline. Curr Opin Pharmacol. 2018;42:81–94. Epub 20180823. doi: 10.1016/j.coph.2018.08.001. PubMed PMID: 30144650; PubMed Central PMCID: PMCPMC6204086.

5. Meghji J, Auld SC, Bisson GP, Khosa C, Masekela R, Navuluri N, et al. Post-tuberculosis lung disease: towards prevention, diagnosis, and care. Lancet Respir Med. 2025;13(5):460–72. Epub 20250322. doi: 10.1016/S2213-2600(24)00429-6. PubMed PMID: 40127662.

6. Dartois VA, Mizrahi V, Savic RM, Silverman JA, Hermann D, Barry CE, 3rd. Strategies for shortening tuberculosis therapy. Nat Med. 2025. Epub 20250613. doi: 10.1038/s41591-025-03742-3. PubMed PMID: 40514466.

7. Hawn TR, Shah JA, Kalman D. New tricks for old dogs: countering antibiotic resistance in tuberculosis with host-directed therapeutics. Immunol Rev. 2015;264(1):344–62. doi: 10.1111/imr.12255. PubMed PMID: 25703571; PubMed Central PMCID: PMCPMC4571192.

8. Kilinc G, Saris A, Ottenhoff THM, Haks MC. Host-directed therapy to combat mycobacterial infections. Immunol Rev. 2021;301(1):62–83. Epub 20210209. doi: 10.1111/imr.12951. PubMed PMID: 33565103; PubMed Central PMCID: PMCPMC8248113.

9. Wallis RS, O’Garra A, Sher A, Wack A. Host-directed immunotherapy of viral and bacterial infections: past, present and future. Nat Rev Immunol. 2023;23(2):121–33. Epub 20220607. doi: 10.1038/s41577-022-00734-z. PubMed PMID: 35672482; PubMed Central PMCID: PMCPMC9171745.

10. Wallis RS, Ginindza S, Beattie T, Arjun N, Likoti M, Edward VA, et al. Adjunctive host-directed therapies for pulmonary tuberculosis: a prospective, open-label, phase 2, randomised controlled trial. Lancet Respir Med. 2021;9(8):897–908. Epub 20210316. doi: 10.1016/S2213-2600(20)30448-3. PubMed PMID: 33740465; PubMed Central PMCID: PMCPMC8332197.

11. Davis JM, Ramakrishnan L. The role of the granuloma in expansion and dissemination of early tuberculous infection. Cell. 2009;136(1):37–49. doi: 10.1016/j.cell.2008.11.014. PubMed PMID: 19135887; PubMed Central PMCID: PMCPMC3134310.

12. Ramakrishnan L. Revisiting the role of the granuloma in tuberculosis. Nat Rev Immunol. 2012;12(5):352–66. Epub 20120420. doi: 10.1038/nri3211. PubMed PMID: 22517424.

13. Pagan AJ, Ramakrishnan L. Immunity and Immunopathology in the Tuberculous Granuloma. Cold Spring Harb Perspect Med. 2014;5(9). Epub 20141106. doi: 10.1101/cshperspect.a018499. PubMed PMID: 25377142; PubMed Central PMCID: PMCPMC4561401.

14. Cronan MR. In the Thick of It: Formation of the Tuberculous Granuloma and Its Effects on Host and Therapeutic Responses. Front Immunol. 2022;13:820134. Epub 20220307. doi: 10.3389/fimmu.2022.820134. PubMed PMID: 35320930; PubMed Central PMCID: PMCPMC8934850.

15. Sarathy JP, Dartois V. Caseum: a Niche for Mycobacterium tuberculosis Drug-Tolerant Persisters. Clin Microbiol Rev. 2020;33(3). Epub 20200401. doi: 10.1128/CMR.00159-19. PubMed PMID: 32238365; PubMed Central PMCID: PMCPMC7117546.

16. Sochocky S. Tuberculoma of the lung. Am Rev Tuberc. 1958;78(3):403–10. doi: 10.1164/artpd.1958.78.3.403. PubMed PMID: 13571599.

17. Monteiro R, Carneiro JC, Costa C, Duarte R. Cerebral tuberculomas - A clinical challenge. Respir Med Case Rep. 2013;9:34–7. Epub 20130603. doi: 10.1016/j.rmcr.2013.04.003. PubMed PMID: 26029627; PubMed Central PMCID: PMCPMC3949551.

18. Lee HS, Oh JY, Lee JH, Yoo CG, Lee CT, Kim YW, et al. Response of pulmonary tuberculomas to anti-tuberculous treatment. Eur Respir J. 2004;23(3):452–5. doi: 10.1183/09031936.04.00087304. PubMed PMID: 15065838.

19. Urbanowski ME, Ordonez AA, Ruiz-Bedoya CA, Jain SK, Bishai WR. Cavitary tuberculosis: the gateway of disease transmission. Lancet Infect Dis. 2020;20(6):e117–e28. Epub 20200505. doi: 10.1016/S1473-3099(20)30148-1. PubMed PMID: 32482293; PubMed Central PMCID: PMCPMC7357333.

20. Lenaerts A, Barry CE, 3rd, Dartois V. Heterogeneity in tuberculosis pathology, microenvironments and therapeutic responses. Immunol Rev. 2015;264(1):288–307. doi: 10.1111/imr.12252. PubMed PMID: 25703567; PubMed Central PMCID: PMCPMC4368385.

21. Parish T. In vitro drug discovery models for Mycobacterium tuberculosis relevant for host infection. Expert Opin Drug Discov. 2020;15(3):349–58. Epub 20200103. doi: 10.1080/17460441.2020.1707801. PubMed PMID: 31899974.

22. Puissegur MP, Botanch C, Duteyrat JL, Delsol G, Caratero C, Altare F. An in vitro dual model of mycobacterial granulomas to investigate the molecular interactions between mycobacteria and human host cells. Cell Microbiol. 2004;6(5):423–33. doi: 10.1111/j.1462-5822.2004.00371.x. PubMed PMID: 15056213.

23. Birkness KA, Guarner J, Sable SB, Tripp RA, Kellar KL, Bartlett J, et al. An in vitro model of the leukocyte interactions associated with granuloma formation in Mycobacterium tuberculosis infection. Immunol Cell Biol. 2007;85(2):160–8. Epub 20070102. doi: 10.1038/sj.icb.7100019. PubMed PMID: 17199112.

24. Guirado E, Mbawuike U, Keiser TL, Arcos J, Azad AK, Wang SH, et al. Characterization of host and microbial determinants in individuals with latent tuberculosis infection using a human granuloma model. mBio. 2015;6(1):e02537–14. Epub 20150217. doi: 10.1128/mBio.02537-14. PubMed PMID: 25691598; PubMed Central PMCID: PMCPMC4337582.

25. Kapoor N, Pawar S, Sirakova TD, Deb C, Warren WL, Kolattukudy PE. Human granuloma in vitro model, for TB dormancy and resuscitation. PLoS One. 2013;8(1):e53657. Epub 20130107. doi: 10.1371/journal.pone.0053657. PubMed PMID: 23308269; PubMed Central PMCID: PMCPMC3538642.

26. Bielecka MK, Tezera LB, Zmijan R, Drobniewski F, Zhang X, Jayasinghe S, et al. A Bioengineered Three-Dimensional Cell Culture Platform Integrated with Microfluidics To Address Antimicrobial Resistance in Tuberculosis. mBio. 2017;8(1). Epub 20170207. doi: 10.1128/mBio.02073-16. PubMed PMID: 28174307; PubMed Central PMCID: PMCPMC5296599.

27. Tezera LB, Bielecka MK, Chancellor A, Reichmann MT, Shammari BA, Brace P, et al. Dissection of the host-pathogen interaction in human tuberculosis using a bioengineered 3-dimensional model. Elife. 2017;6. Epub 20170107. doi: 10.7554/eLife.21283. PubMed PMID: 28063256; PubMed Central PMCID: PMCPMC5238961.

28. Kotze LA, Beltran CGG, Lang D, Loxton AG, Cooper S, Meiring M, et al. Establishment of a Patient-Derived, Magnetic Levitation-Based, Three-Dimensional Spheroid Granuloma Model for Human Tuberculosis. mSphere. 2021;6(4):e0055221. Epub 20210721. doi: 10.1128/mSphere.00552-21. PubMed PMID: 34287004; PubMed Central PMCID: PMCPMC8386456.

29. Arbues A, Schmidiger S, Kammuller M, Portevin D. Extracellular Matrix-Induced GM-CSF and Hypoxia Promote Immune Control of Mycobacterium tuberculosis in Human In Vitro Granulomas. Front Immunol. 2021;12:727508. Epub 20210917. doi: 10.3389/fimmu.2021.727508. PubMed PMID: 34603299; PubMed Central PMCID: PMCPMC8486295.

30. Elkington P, Lerm M, Kapoor N, Mahon R, Pienaar E, Huh D, et al. In Vitro Granuloma Models of Tuberculosis: Potential and Challenges. J Infect Dis. 2019;219(12):1858–66. doi: 10.1093/infdis/jiz020. PubMed PMID: 30929010; PubMed Central PMCID: PMCPMC6534193.

31. Silva-Miranda M, Ekaza E, Breiman A, Asehnoune K, Barros-Aguirre D, Pethe K, et al. High-content screening technology combined with a human granuloma model as a new approach to evaluate the activities of drugs against Mycobacterium tuberculosis. Antimicrob Agents Chemother. 2015;59(1):693–7. Epub 20141027. doi: 10.1128/AAC.03705-14. PubMed PMID: 25348525; PubMed Central PMCID: PMCPMC4291390.

32. Thacker VV, Dhar N, Sharma K, Barrile R, Karalis K, McKinney JD. A lung-on-chip model of early Mycobacterium tuberculosis infection reveals an essential role for alveolar epithelial cells in controlling bacterial growth. Elife. 2020;9. Epub 20201124. doi: 10.7554/eLife.59961. PubMed PMID: 33228849; PubMed Central PMCID: PMCPMC7735758.

33. Kim SY, Choi JA, Choi S, Kim KK, Song CH, Kim EM. Advances in an In Vitro Tuberculosis Infection Model Using Human Lung Organoids for Host-Directed Therapies. PLoS Pathog. 2024;20(7):e1012295. Epub 20240725. doi: 10.1371/journal.ppat.1012295. PubMed PMID: 39052544; PubMed Central PMCID: PMCPMC11271890.

34. Cronan MR, Matty MA, Rosenberg AF, Blanc L, Pyle CJ, Espenschied ST, et al. An explant technique for high-resolution imaging and manipulation of mycobacterial granulomas. Nat Methods. 2018;15(12):1098–107. Epub 20181130. doi: 10.1038/s41592-018-0215-8. PubMed PMID: 30504889; PubMed Central PMCID: PMCPMC6312189.

35. Bryk R, Mundhra S, Jiang X, Wood M, Pfau D, Weber E, et al. Potentiation of rifampin activity in a mouse model of tuberculosis by activation of host transcription factor EB. PLoS Pathog. 2020;16(6):e1008567. Epub 20200623. doi: 10.1371/journal.ppat.1008567. PubMed PMID: 32574211; PubMed Central PMCID: PMCPMC7337396.

36. Arkenau HT, Plummer R, Molife LR, Olmos D, Yap TA, Squires M, et al. A phase I dose escalation study of AT9283, a small molecule inhibitor of aurora kinases, in patients with advanced solid malignancies. Ann Oncol. 2012;23(5):1307–13. Epub 20111019. doi: 10.1093/annonc/mdr451. PubMed PMID: 22015452.

37. Hay AE, Murugesan A, DiPasquale AM, Kouroukis T, Sandhu I, Kukreti V, et al. A phase II study of AT9283, an aurora kinase inhibitor, in patients with relapsed or refractory multiple myeloma: NCIC clinical trials group IND.191. Leuk Lymphoma. 2016;57(6):1463–6. Epub 20151015. doi: 10.3109/10428194.2015.1091927. PubMed PMID: 26376958.

38. Stanley SA, Barczak AK, Silvis MR, Luo SS, Sogi K, Vokes M, et al. Identification of host-targeted small molecules that restrict intracellular Mycobacterium tuberculosis growth. PLoS Pathog. 2014;10(2):e1003946. Epub 20140220. doi: 10.1371/journal.ppat.1003946. PubMed PMID: 24586159; PubMed Central PMCID: PMCPMC3930586.

39. Korbee CJ, Heemskerk MT, Kocev D, van Strijen E, Rabiee O, Franken K, et al. Combined chemical genetics and data-driven bioinformatics approach identifies receptor tyrosine kinase inhibitors as host-directed antimicrobials. Nat Commun. 2018;9(1):358. Epub 20180124. doi: 10.1038/s41467-017-02777-6. PubMed PMID: 29367740; PubMed Central PMCID: PMCPMC5783939.

40. Shapira T, Rankine-Wilson L, Chao JD, Pichler V, Rens C, Pfeifer T, et al. High-Content Screening of Eukaryotic Kinase Inhibitors Identify CHK2 Inhibitor Activity Against Mycobacterium tuberculosis. Front Microbiol. 2020;11:553962. Epub 20200918. doi: 10.3389/fmicb.2020.553962. PubMed PMID: 33042061; PubMed Central PMCID: PMCPMC7530171.

41. Kumar D, Nath L, Kamal MA, Varshney A, Jain A, Singh S, et al. Genome-wide analysis of the host intracellular network that regulates survival of Mycobacterium tuberculosis. Cell. 2010;140(5):731–43. doi: 10.1016/j.cell.2010.02.012. PubMed PMID: 20211141.

42. Lai Y, Babunovic GH, Cui L, Dedon PC, Doench JG, Fortune SM, et al. Illuminating Host-Mycobacterial Interactions with Genome-wide CRISPR Knockout and CRISPRi Screens. Cell Syst. 2020;11(3):239–51 e7. doi: 10.1016/j.cels.2020.08.010. PubMed PMID: 32970993.

43. Lin PL, Ford CB, Coleman MT, Myers AJ, Gawande R, Ioerger T, et al. Sterilization of granulomas is common in active and latent tuberculosis despite within-host variability in bacterial killing. Nat Med. 2014;20(1):75–9. Epub 20131215. doi: 10.1038/nm.3412. PubMed PMID: 24336248; PubMed Central PMCID: PMCPMC3947310.

44. Behr MA, Wilson MA, Gill WP, Salamon H, Schoolnik GK, Rane S, et al. Comparative genomics of BCG vaccines by whole-genome DNA microarray. Science. 1999;284(5419):1520-3. doi: 10.1126/science.284.5419.1520. PubMed PMID: 10348738.

45. Volkman HE, Clay H, Beery D, Chang JC, Sherman DR, Ramakrishnan L. Tuberculous granuloma formation is enhanced by a mycobacterium virulence determinant. PLoS Biol. 2004;2(11):e367. Epub 20041026. doi: 10.1371/journal.pbio.0020367. PubMed PMID: 15510227; PubMed Central PMCID: PMCPMC524251.

46. Cronan MR, Beerman RW, Rosenberg AF, Saelens JW, Johnson MG, Oehlers SH, et al. Macrophage Epithelial Reprogramming Underlies Mycobacterial Granuloma Formation and Promotes Infection. Immunity. 2016;45(4):861–76. doi: 10.1016/j.immuni.2016.09.014. PubMed PMID: 27760340; PubMed Central PMCID: PMCPMC5268069.

47. Mattila JT, Ojo OO, Kepka-Lenhart D, Marino S, Kim JH, Eum SY, et al. Microenvironments in tuberculous granulomas are delineated by distinct populations of macrophage subsets and expression of nitric oxide synthase and arginase isoforms. J Immunol. 2013;191(2):773–84. Epub 20130607. doi: 10.4049/jimmunol.1300113. PubMed PMID: 23749634; PubMed Central PMCID: PMCPMC3746594.

48. McCaffrey EF, Donato M, Keren L, Chen Z, Delmastro A, Fitzpatrick MB, et al. The immunoregulatory landscape of human tuberculosis granulomas. Nat Immunol. 2022;23(2):318–29. Epub 20220120. doi: 10.1038/s41590-021-01121-x. PubMed PMID: 35058616; PubMed Central PMCID: PMCPMC8810384.

49. Sawyer AJ, Patrick E, Edwards J, Wilmott JS, Fielder T, Yang Q, et al. Spatial mapping reveals granuloma diversity and histopathological superstructure in human tuberculosis. J Exp Med. 2023;220(6). Epub 20230315. doi: 10.1084/jem.20221392. PubMed PMID: 36920308.

50. Hillman H, Khan N, Singhania A, Dubelko P, Soldevila F, Tippalagama R, et al. Single-cell profiling reveals distinct subsets of CD14+ monocytes drive blood immune signatures of active tuberculosis. Front Immunol. 2022;13:1087010. Epub 20230111. doi: 10.3389/fimmu.2022.1087010. PubMed PMID: 36713384; PubMed Central PMCID: PMCPMC9874319.

51. Oehlers SH, Cronan MR, Scott NR, Thomas MI, Okuda KS, Walton EM, et al. Interception of host angiogenic signalling limits mycobacterial growth. Nature. 2015;517(7536):612-5. Epub 20141124. doi: 10.1038/nature13967. PubMed PMID: 25470057; PubMed Central PMCID: PMCPMC4312197.

52. Chandra P, Grigsby SJ, Philips JA. Immune evasion and provocation by Mycobacterium tuberculosis. Nat Rev Microbiol. 2022;20(12):750–66. Epub 20220725. doi: 10.1038/s41579-022-00763-4. PubMed PMID: 35879556; PubMed Central PMCID: PMCPMC9310001.

53. Kimura S, Noda T, Yoshimori T. Dissection of the autophagosome maturation process by a novel reporter protein, tandem fluorescent-tagged LC3. Autophagy. 2007;3(5):452–60. Epub 20070521. doi: 10.4161/auto.4451. PubMed PMID: 17534139.

54. Beckwith KS, Beckwith MS, Ullmann S, Saetra RS, Kim H, Marstad A, et al. Plasma membrane damage causes NLRP3 activation and pyroptosis during Mycobacterium tuberculosis infection. Nat Commun. 2020;11(1):2270. Epub 20200508. doi: 10.1038/s41467-020-16143-6. PubMed PMID: 32385301; PubMed Central PMCID: PMCPMC7210277.

55. Zenk SF, Hauck S, Mayer D, Grieshober M, Stenger S. Stabilization of Hypoxia-Inducible Factor Promotes Antimicrobial Activity of Human Macrophages Against Mycobacterium tuberculosis. Front Immunol. 2021;12:678354. Epub 20210602. doi: 10.3389/fimmu.2021.678354. PubMed PMID: 34149713; PubMed Central PMCID: PMCPMC8206807.

56. Flynn JL, Goldstein MM, Chan J, Triebold KJ, Pfeffer K, Lowenstein CJ, et al. Tumor necrosis factor-alpha is required in the protective immune response against Mycobacterium tuberculosis in mice. Immunity. 1995;2(6):561–72. doi: 10.1016/1074-7613(95)90001-2. PubMed PMID: 7540941.

57. Roca FJ, Whitworth LJ, Prag HA, Murphy MP, Ramakrishnan L. Tumor necrosis factor induces pathogenic mitochondrial ROS in tuberculosis through reverse electron transport. Science. 2022;376(6600):eabh2841. Epub 20220624. doi: 10.1126/science.abh2841. PubMed PMID: 35737799; PubMed Central PMCID: PMCPMC7612974.

58. Lin PL, Myers A, Smith L, Bigbee C, Bigbee M, Fuhrman C, et al. Tumor necrosis factor neutralization results in disseminated disease in acute and latent Mycobacterium tuberculosis infection with normal granuloma structure in a cynomolgus macaque model. Arthritis Rheum. 2010;62(2):340–50. doi: 10.1002/art.27271. PubMed PMID: 20112395; PubMed Central PMCID: PMCPMC3047004.

59. Iliopoulos A, Psathakis K, Aslanidis S, Skagias L, Sfikakis PP. Tuberculosis and granuloma formation in patients receiving anti-TNF therapy. Int J Tuberc Lung Dis. 2006;10(5):588–90. PubMed PMID: 16704045.

60. Lu Q, Liu J, Yu Y, Liang HF, Zhang SQ, Li ZB, et al. ALB, HP, OAF and RBP4 as novel protein biomarkers for identifying cured patients with pulmonary tuberculosis by DIA. Clin Chim Acta. 2022;535:82–91. Epub 20220812. doi: 10.1016/j.cca.2022.08.002. PubMed PMID: 35964702.

61. Reichmann MT, Tezera LB, Vallejo AF, Vukmirovic M, Xiao R, Reynolds J, et al. Integrated transcriptomic analysis of human tuberculosis granulomas and a biomimetic model identifies therapeutic targets. J Clin Invest. 2021;131(15). doi: 10.1172/JCI148136. PubMed PMID: 34128839; PubMed Central PMCID: PMCPMC8321576.

62. Ahmed M, Thirunavukkarasu S, Rosa BA, Thomas KA, Das S, Rangel-Moreno J, et al. Immune correlates of tuberculosis disease and risk translate across species. Sci Transl Med. 2020;12(528). doi: 10.1126/scitranslmed.aay0233. PubMed PMID: 31996462; PubMed Central PMCID: PMCPMC7354419.

63. Singhania A, Verma R, Graham CM, Lee J, Tran T, Richardson M, et al. A modular transcriptional signature identifies phenotypic heterogeneity of human tuberculosis infection. Nat Commun. 2018;9(1):2308. Epub 20180619. doi: 10.1038/s41467-018-04579-w. PubMed PMID: 29921861; PubMed Central PMCID: PMCPMC6008327.

64. Prideaux B, Via LE, Zimmerman MD, Eum S, Sarathy J, O’Brien P, et al. The association between sterilizing activity and drug distribution into tuberculosis lesions. Nat Med. 2015;21(10):1223–7. Epub 20150907. doi: 10.1038/nm.3937. PubMed PMID: 26343800; PubMed Central PMCID: PMCPMC4598290.

65. Manca C, Tsenova L, Barry CE, 3rd, Bergtold A, Freeman S, Haslett PA, et al. Mycobacterium tuberculosis CDC1551 induces a more vigorous host response in vivo and in vitro, but is not more virulent than other clinical isolates. J Immunol. 1999;162(11):6740–6. PubMed PMID: 10352293.

66. Rybniker J, Vocat A, Sala C, Busso P, Pojer F, Benjak A, et al. Lansoprazole is an antituberculous prodrug targeting cytochrome bc1. Nat Commun. 2015;6:7659. Epub 20150709. doi: 10.1038/ncomms8659. PubMed PMID: 26158909; PubMed Central PMCID: PMCPMC4510652.

67. Matty MA, Knudsen DR, Walton EM, Beerman RW, Cronan MR, Pyle CJ, et al. Potentiation of P2RX7 as a host-directed strategy for control of mycobacterial infection. Elife. 2019;8. Epub 20190129. doi: 10.7554/eLife.39123. PubMed PMID: 30693866; PubMed Central PMCID: PMCPMC6351102.

68. Bailey MA, Na H, Duthie MS, Gillis TP, Lahiri R, Parish T. Nitazoxanide is active against Mycobacterium leprae. PLoS One. 2017;12(8):e0184107. Epub 20170829. doi: 10.1371/journal.pone.0184107. PubMed PMID: 28850614; PubMed Central PMCID: PMCPMC5574600.

69. Miow QH, Vallejo AF, Wang Y, Hong JM, Bai C, Teo FS, et al. Doxycycline host-directed therapy in human pulmonary tuberculosis. J Clin Invest. 2021;131(15). doi: 10.1172/JCI141895. PubMed PMID: 34128838; PubMed Central PMCID: PMCPMC8321570.

70. Golovkine GR, Roberts AW, Morrison HM, Rivera-Lugo R, McCall RM, Nilsson H, et al. Autophagy restricts Mycobacterium tuberculosis during acute infection in mice. Nat Microbiol. 2023;8(5):819–32. Epub 20230410. doi: 10.1038/s41564-023-01354-6. PubMed PMID: 37037941.

71. Rastogi S, Briken V. Interaction of Mycobacteria With Host Cell Inflammasomes. Front Immunol. 2022;13:791136. Epub 20220214. doi: 10.3389/fimmu.2022.791136. PubMed PMID: 35237260; PubMed Central PMCID: PMCPMC8882646.

72. Arbues A, Brees D, Chibout SD, Fox T, Kammuller M, Portevin D. TNF-alpha antagonists differentially induce TGF-beta1-dependent resuscitation of dormant-like Mycobacterium tuberculosis. PLoS Pathog. 2020;16(2):e1008312. Epub 20200218. doi: 10.1371/journal.ppat.1008312. PubMed PMID: 32069329; PubMed Central PMCID: PMCPMC7048311.

73. Mukundan S, Bhatt R, Lucas J, Tereyek M, Chang TL, Subbian S, et al. 3D host cell and pathogen-based bioassay development for testing anti-tuberculosis (TB) drug response and modeling immunodeficiency. Biomol Concepts. 2021;12(1):117–28. Epub 20210902. doi: 10.1515/bmc-2021-0013. PubMed PMID: 34473918.

74. Armesilla-Diaz A, Pilar Arenaz M, Ashby C, Blanco D, D’Oria E, Garuti H, et al. High-throughput screening of small molecules targeting Mycobacterium tuberculosis in human iPSC macrophages. Antimicrob Agents Chemother. 2025:e0161324. Epub 20250527. doi: 10.1128/aac.01613-24. PubMed PMID: 40423030.

75. Lyadova I, Gerasimova T, Nenasheva T. Macrophages Derived From Human Induced Pluripotent Stem Cells: The Diversity of Protocols, Future Prospects, and Outstanding Questions. Front Cell Dev Biol. 2021;9:640703. Epub 20210602. doi: 10.3389/fcell.2021.640703. PubMed PMID: 34150747; PubMed Central PMCID: PMCPMC8207294.

76. Kang JS, Lee Y, Lee Y, Gil D, Kim MJ, Wood C, et al. Generation of induced alveolar assembloids with functional alveolar-like macrophages. Nat Commun. 2025;16(1):3346. Epub 20250409. doi: 10.1038/s41467-025-58450-w. PubMed PMID: 40199883; PubMed Central PMCID: PMCPMC11978882.

77. Choi SS, van Unen V, Zhang H, Rustagi A, Alwahabi SA, Santos AJM, et al. Organoid modeling of lung-resident immune responses to SARS-CoV-2 infection. Res Sq. 2023. Epub 20230505. doi: 10.21203/rs.3.rs-2870695/v1. PubMed PMID: 37205380; PubMed Central PMCID: PMCPMC10187413.

78. Tiwari SK, Wong WJ, Moreira M, Pasqualini C, Ginhoux F. Induced pluripotent stem cell-derived macrophages as a platform for modelling human disease. Nat Rev Immunol. 2025;25(2):108–24. Epub 20240927. doi: 10.1038/s41577-024-01081-x. PubMed PMID: 39333753.

79. Gaeddert M, Glaser K, Chendi BH, Sultanli A, Koeppel L, MacLean EL, et al. Host blood protein biomarkers to screen for tuberculosis disease: a systematic review and meta-analysis. J Clin Microbiol. 2024;62(11):e0078624. Epub 20241024. doi: 10.1128/jcm.00786-24. PubMed PMID: 39445833; PubMed Central PMCID: PMCPMC11559064.

80. Foreman TW, Nelson CE, Sallin MA, Kauffman KD, Sakai S, Otaizo-Carrasquero F, et al. CD30 co-stimulation drives differentiation of protective T cells during Mycobacterium tuberculosis infection. J Exp Med. 2023;220(8). Epub 20230425. doi: 10.1084/jem.20222090. PubMed PMID: 37097292; PubMed Central PMCID: PMCPMC10130742.

81. Au WY, Leung AY, Tse EW, Cheung WW, Shek TW, Kwong YL. High incidence of tuberculosis after alemtuzumab treatment in Hong Kong Chinese patients. Leuk Res. 2008;32(4):547–51. Epub 20070821. doi: 10.1016/j.leukres.2007.06.010. PubMed PMID: 17714782.

82. Ghosh S, Chackerian AA, Parker CM, Ballantyne CM, Behar SM. The LFA-1 adhesion molecule is required for protective immunity during pulmonary Mycobacterium tuberculosis infection. J Immunol. 2006;176(8):4914–22. doi: 10.4049/jimmunol.176.8.4914. PubMed PMID: 16585587.

83. Datta M, Via LE, Kamoun WS, Liu C, Chen W, Seano G, et al. Anti-vascular endothelial growth factor treatment normalizes tuberculosis granuloma vasculature and improves small molecule delivery. Proc Natl Acad Sci U S A. 2015;112(6):1827–32. Epub 20150126. doi: 10.1073/pnas.1424563112. PubMed PMID: 25624495; PubMed Central PMCID: PMCPMC4330784.

84. Napier RJ, Rafi W, Cheruvu M, Powell KR, Zaunbrecher MA, Bornmann W, et al. Imatinib-sensitive tyrosine kinases regulate mycobacterial pathogenesis and represent therapeutic targets against tuberculosis. Cell Host Microbe. 2011;10(5):475–85. doi: 10.1016/j.chom.2011.09.010. PubMed PMID: 22100163; PubMed Central PMCID: PMCPMC3222875.

85. Singhal A, Jie L, Kumar P, Hong GS, Leow MK, Paleja B, et al. Metformin as adjunct antituberculosis therapy. Sci Transl Med. 2014;6(263):263ra159. doi: 10.1126/scitranslmed.3009885. PubMed PMID: 25411472.

86. Frenkel JDH, Ackart DF, Todd AK, DiLisio JE, Hoffman S, Tanner S, et al. Metformin enhances protection in guinea pigs chronically infected with Mycobacterium tuberculosis. Sci Rep. 2020;10(1):16257. Epub 20201001. doi: 10.1038/s41598-020-73212-y. PubMed PMID: 33004826; PubMed Central PMCID: PMCPMC7530990.

87. Daniels JM, Vonk-Noordegraaf A, Janssen JJ, Postmus PE, van Altena R. Tuberculosis complicating imatinib treatment for chronic myeloid leukaemia. Eur Respir J. 2009;33(3):670–2. doi: 10.1183/09031936.00025408. PubMed PMID: 19251803.

88. Sathkumara HD, Hansen K, Miranda-Hernandez S, Govan B, Rush CM, Henning L, et al. Disparate Effects of Metformin on Mycobacterium tuberculosis Infection in Diabetic and Nondiabetic Mice. Antimicrob Agents Chemother. 2020;65(1). Epub 20201216. doi: 10.1128/AAC.01422-20. PubMed PMID: 33046495; PubMed Central PMCID: PMCPMC7927854.

89. Yates TA, Tomlinson LA, Bhaskaran K, Langan S, Thomas S, Smeeth L, et al. Lansoprazole use and tuberculosis incidence in the United Kingdom Clinical Practice Research Datalink: A population based cohort. PLoS Med. 2017;14(11):e1002457. Epub 20171121. doi: 10.1371/journal.pmed.1002457. PubMed PMID: 29161254; PubMed Central PMCID: PMCPMC5697821.

90. Kamariza M, Shieh P, Ealand CS, Peters JS, Chu B, Rodriguez-Rivera FP, et al. Rapid detection of Mycobacterium tuberculosis in sputum with a solvatochromic trehalose probe. Sci Transl Med. 2018;10(430). doi: 10.1126/scitranslmed.aam6310. PubMed PMID: 29491187; PubMed Central PMCID: PMCPMC5985656.

91. Sable SB, Kline A, Li W, Posey JE. Methodology for a freshly engineered or cryo-preserved 3D tuberculoma bioplatform for studying tuberculosis biology and high-content screening of therapeutics. bioRxiv. 2025.

92. Wells G, Glasgow JN, Nargan K, Lumamba K, Madansein R, Maharaj K, et al. Micro-Computed Tomography Analysis of the Human Tuberculous Lung Reveals Remarkable Heterogeneity in Three-dimensional Granuloma Morphology. Am J Respir Crit Care Med. 2021;204(5):583–95. doi: 10.1164/rccm.202101-0032OC. PubMed PMID: 34015247; PubMed Central PMCID: PMCPMC8491258.

