## Supplemental Methods, Supplementary Figures and Legends for Supplementary Tables for "Identification and characterization of host-directed therapeutics for tuberculosis using a versatile human 3D tuberculoma bioplatform"

Supplementary information includes Supplementary Materials and Methods (S1 Text), Supplementary Figures (S1−S25 Figs), and legends for Supplementary Figures and Tables (S1−S9 Tables).

### Supplementary materials and methods

#### Human donors

Blood from eight healthy adult donors (four males and four females, aged 23−55 years) of diverse ethnicities was used in different aspects of the study. All donors were HIV and hepatitis-negative, with no clinical signs of TB. Exclusions were made for those on medication or with metabolic disorders. All donors were BCG vaccination-negative and had negative tuberculin skin test results, with no known history of contact with TB patients. The exposure of donors to environmental mycobacteria is unknown; all resided in the Atlanta area, which has a very low incidence of TB or atypical mycobacterial infections. Total peripheral blood mononuclear cells (PBMCs) or purified PBMC subsets were used for 3D cell culture. Specifically, PBMCs from four healthy donors were used to purify $CD14^+$ monocytes and $CD14^−$ cell subsets and to generate 3D tuberculomas in 3D cell culture plates (Corning, 4515). For screening assays using the tuberculoma bioplatform, 250 mL of peripheral blood from each donor was used per assay to purify $CD14^+$ monocytes and $CD14^−$ cell subsets. Blood from the same four donors was used to purify cell subsets and generate 3D tuberculomas for screening 65 HDT compounds in assays investigating bacterial burden inhibition, granuloma lesion resolution, and cytotoxicity.

#### Growth of cell lines

Immortalized cell lines THP-1, AMJ2, U937, RAW264.7, and J774A.1 were maintained and grown according to ATCC's instructions at the CDC cell culture core facility. Briefly, AMJ2, a mouse alveolar macrophage cell line, was cultured in complete Dulbecco's modified Eagle's medium (DMEM) with 4 mM L-glutamine adjusted to contain 1.5 g/L sodium bicarbonate and 4.5 g/L glucose and supplemented with 5 mM HEPES and fetal bovine serum (FBS, 5%). Mouse ascites macrophage cell lines RAW264.7 and J774A.1 were cultured in complete DMEM containing 4 mM L-glutamine, 4.5 g/L glucose, 1 mM sodium pyruvate, and 1.5 g/L sodium bicarbonate and supplemented with 10% FBS. Human pleural histiocyte and blood monocytic cell lines U937 and THP-1 were cultured in complete RPMI-1640 medium modified to contain 2 mM L-glutamine, 10 mM HEPES, 1 mM sodium pyruvate, 4.5 g/L glucose, and 1.5 g/L sodium bicarbonate and supplemented with 10% FBS. For the THP-1 cell line, the above-described complete RPMI-1640 medium was supplemented with 2-mercaptoethanol to a final concentration of 0.05 mM. Reporter THP-1 monocytes stably expressing copGFP or RFP (Applied Biological Materials) were cultured in complete RPMI medium containing L-glutamine (1mM), 100 mM sodium pyruvate (1%), 1M HEPES (1%), 10,000 units/ml penicillin-streptomycin solution (1%), and 10% FBS, with puromycin (1.2 µg/ml) for selection. Reporter THP-1 ASC-GFP and THP-1-Difluo hLC3 cell lines (InvivoGen) were grown in complete RPMI-1640 medium (described above) supplemented with Normocin (100 µg/ml). Zeocin (200 µg/ml) was used for the selection of ASC-GFP-expressing cells. All cell lines were cultured in a cell culture incubator at 37 °C, 5% $CO_2$, and 100% humidity.

### Cell encapsulation

THP-1 monocytes, pre-infected or mixed with *Mycobacterium marinum* (*Mm*) 'M' (tdTomato), were encapsulated using the Austrianova 'Cell-in-a-Box®' kit (Sigma-Aldrich), which employs a biocompatible cellulose-based polymer for cell encapsulation. This method is advantageous over traditional encapsulation techniques that require large, expensive, and complex machinery. Briefly, THP-1 monocytes were washed with a prewarmed (37 ºC) complete RPMI-1640 medium without antibiotics (a co-culture medium) to remove antibiotics from the growth medium. Cells were pelleted by centrifugation at 250 g for 6 minutes and resuspended in a co-culture medium. They were infected with *Mm* 'M' (tdTomato) (MOI 0.008) in a Corning 75 $cm^2$ tissue culture flask and incubated overnight at 37 °C in a $CO_2$ incubator. The next day, after washing the infected cells three times and pelleting them using low-speed centrifugation at 50 g for 5 minutes, they were resuspended in a co-culture medium and prepared for encapsulation. Infected monocytes ($2\times10^6$ live cells/ml) were placed in a sterile 1.5 ml microcentrifuge tube and pelleted by centrifugation at 250 g for 6 minutes. Alternatively, THP-1 monocytes ($2\times10^6$ live cells/ml) were mixed with *Mm* 'M' (tdTomato) in a microcentrifuge tube (MOI 0.008) and pelleted by centrifugation for 30 minutes. After discarding the supernatant, cells were mixed with Solution 1 from the kit and added dropwise with constant stirring into Solution 2, the hardening bath, using a 1 ml Luer lock syringe fitted with a filling needle (G18½, blunt end) to draw up the suspension from the microcentrifuge tube and a droplet needle (G34, blunt end) to dispense the droplets. The droplets were dispensed into Solution 2 at a moderate speed of 1–2 drops per second, maintaining the same drop height. After the droplets hardened into beads, they underwent several wash steps and were then transferred to the co-culture medium. The beads, approximately 2 mm in diameter with pores allowing nutrient exchange while retaining cells, were cultured in a 96-well flat-bottom (2D) ULA microplate (Corning) for 12 days. Cells proliferated over 12 days and colonized within the beads. On day 12, images of the beads encapsulating the 3D co-culture were captured using the Cytation-5 cell imager.

#### Preparation of extracellular matrix solution

To prepare the ECM solution for 3D cell culture, human VitroCol type I collagen (3 mg/ml, Advanced Biomatrix) and human fibronectin (0.1 %, Sigma-Aldrich) solutions were used. VitroCol human atelocollagen comprises approximately 97% type I collagen, with the remainder consisting of type III collagen. Gentle yet thorough mixing and pH monitoring are critical while preparing the ECM mixture. Keep the mixture at 4 °C in the refrigerator or ice to prevent gelation. To prepare the mix, 1 part of chilled 10× PBS was added to 8 parts of chilled VitroCol collagen solution with gentle swirling. The pH of the mixture was adjusted to 7.2–7.6 using sterile 0.1M NaOH, and the final volume was adjusted to 10 parts with cell culture-grade water. Finally, 20 µl of human fibronectin solution was added per 5 ml of collagen mixture, mixing gently by pipetting.

#### ELISPOT assay

The 96-well ELISPOT plates (Millipore, S2EM004M99) were coated overnight at 4°C with 100 µl of cytokine- or protein-specific capture antibodies diluted in PBS (pH 7.2) according to the manufacturer's instructions. The next day, after washing twice with RPMI-1640 to remove unbound antibodies, the microwell membranes were blocked with RPMI-1640 containing 10% FBS (Atlas Biologicals) for 2 hours at room temperature. A single-cell suspension was prepared from dissociated 3D spheroid cultures, and cells from multiple infected or control spheroids from the plate were combined at individual time points. Live cells were counted using the Trypan blue dye exclusion method and plated at a concentration of $1\times10^5$ live cells per well in complete RPMI-1640 medium without additional mycobacterial antigen stimulation. The *E. coli* LPS (10 µg/ml; Sigma-Aldrich) was used as a positive control for cell reactivity. After incubation at 37°C with 5% CO2 and 100% humidity for 16 hours, the wells were washed twice with cell culture-grade water and three times with PBS-Tween-20 wash buffer. A 100 µl per well of biotin-labeled detection antibodies diluted in dilution buffer, PBS with 10% FBS or 1% BSA, was added. The plates were incubated at room temperature for 2 hours. Following three washes with wash buffer, 100 µl per well of the diluted horseradish peroxidase-conjugated streptavidin in the dilution buffer was added. The reaction was developed using a 3-amino-9-ethylcarbazole (AEC) substrate reagent set (BD Biosciences). The spot-forming units (SFUs) were counted using an ELISPOT reader (Cellular Technology Limited).

#### CytoTox-Glo™ cytotoxicity assay

The assay was performed using the CytoTox-Glo™ Cytotoxicity Assay kit (Promega) according to the manufacturer's instructions. Briefly, 3D spheroids in cell culture media were dissociated by pipetting, and single-cell suspensions were prepared in the microwells. The cell suspension in each well was divided into equal volumes (100 µl each) and transferred into two wells (e.g., wells A1 and A2) in a luminescence plate (Corning Costar, Cat #3917) with a solid white, flat bottom. In well A1, 50 µl of Lysis Reagent (digitonin, 30 µg/ml, final concentration) was added, and in well A2, 50 µl of complete RPMI-1640 medium was added. CytoTox-Glo™ Assay Reagent was prepared by transferring the contents of one bottle of Assay Buffer to the AAF-Glo™ Substrate bottle. A prewarmed (37 ºC) CytoTox-Glo™ Assay Reagent (50 µl) was added to all wells (1:4 dilution), followed by a 15-minute incubation at room temperature with orbital shaking at 700−900 rpm. Luminescence was measured using a Cytation-5 Multimode Reader (Agilent-BioTek). Control samples were included in each plate, and a 2-fold serial dilution of freshly grown THP-1 cells (starting from $5\times10^6$ cells per well) was used to create a standard curve. The total number of cells was determined from well A1, and the number of dead cells was determined from well A2. The data analysis was performed using Microsoft Excel. The cytotoxicity was calculated using the following formula.

$$\text{Cell cytotoxicity} = \frac{(\text{Total number of cells from well A2})\times 100\%}{\text{Total number of cells from well A1}}$$

The CytoTox-Glo™ assay is highly sensitive, capable of detecting as few as 200−500 dead cells in a population of 10,000 and less than 10 dead cells in a limiting dilution series.

#### Determining $CC_{50}$ and $EC_{50}$ values

To determine the 50% cytotoxic concentration ($CC_{50}$) of chemical compounds, 3D spheroid cultures of THP-1-RFP reporter cells (Applied Biological Materials) generated in 96-well plates (Corning, 4515) were exposed to a range of doses of individual compounds, starting at 20 µM on day 6. Exposures were performed in 50 µl of complete RPMI-1640 medium. The cytotoxicity was determined by measuring the inhibition of red fluorescence (excitation at 588 and emission at 633 nm) resulting from cell death on day 12 using a Cytation-5. $CC_{50}$ values were calculated using GraphPad Prism v9.3. To assess efficacy against mycobacteria in tuberculomas, 3D co-cultures of *Mm* ‘M’ (tdTomato) and THP-1 cells were treated with a range of doses of each compound, like those used for determining $CC_{50}$ concentrations. Mycobacterial growth was monitored in real-time by measuring the tdTomato RFUs, a reliable surrogate for CFUs. Six different concentrations of each compound were tested to determine the concentration required to inhibit *Mm* (tdTomato) growth by 50%. A half-maximal effective concentration ($EC_{50}$) was determined for each of the tested compounds. The ability of a compound to inhibit ≥50% *Mm* growth at non-toxic doses was ascertained. A favorable therapeutic index, as measured by the $CC_{50}/EC_{50}$, was established.

#### Flow cytometry analysis

Cells were acquired on a BD LSRFortessaTM flow cytometer equipped with four lasers (405 nm, 488 nm, 532 nm, and 633 nm) and analyzed using FlowJo v10.10 software (Tree Star Inc.). At least 55,000 events were acquired for each sample, as described previously [1]. One infected or uninfected control 3D sample per time point consisted of cells isolated and combined from six 3D spheroids to capture these events. One THP-1 control sample consisted of a 2D suspension cell culture pooled from three tissue culture flasks. The Uniform Manifold Approximation and Projection (UMAP) method, a dimensionality reduction technique to visualize high-dimensional data, was used to visualize the cells in a 2D embedding [2]. We used human canonical marker knowledge to classify cells into different cell subtypes.

#### RNA sequencing analyses

Raw FASTQ files underwent preprocessing for RNA sequencing analysis using the FastQC tool to assess QC score, GC content, and adapter contaminations. Before proceeding to downstream analyses, low-quality reads and adapter sequences were trimmed with Trimmomatic [3], discarding sequence reads shorter than 35 base pairs. The quality of the trimmed reads was re-evaluated using FastQC to retain sequences with high-quality scores. The reference genome and corresponding GTF files were sourced from the GENCODE website using the primary assembly version (Reference genome: GRCh38.primary_assembly.genome.fa.gz, GTF file: gencode.v38.primary_assembly.annoation.gtf.gz) [4]. Stranded forward reads were aligned to the reference genome using the STAR aligner and annotated by the GTF file [5]. The output SAM files were sorted and converted to BAM files, and MultiQC provided a comprehensive report for trimming and mapping quality [6]. Downstream analyses were conducted in R. To count reads mapped to the genes, sorted BAM files were processed with the featureCounts function from the RSubread package [7]. The generated count table was normalized based on the sequencing depth using DESeq2 [8]. A preliminary data evaluation included a principal component analysis (PCA) plot and a hierarchical clustering heatmap derived from a sample distance matrix, using DESeq2's built-in functions with 'regularized log-transformed' count reads.

Raw count reads were subjected to differential gene expression analysis using the DESeq2 package. Differential gene expression analysis was performed between 3D-infected and control samples using the DESeq2 package. The fold-change in gene expression of *Mm* 'M'-infected samples over corresponding uninfected control samples isolated at a given time-point in culture was determined. Genes were considered differentially expressed if they had a Benjamin-Hochberg adjusted $p$-value < 0.05 and an absolute $\log_2$ fold-change >1.0 [9]. Significantly up-regulated and down-regulated genes ($p < 0.05$) were visualized using volcano plots and MA plots, utilizing DESeq2's built-in function. Gene Set Enrichment Analysis (GSEA) was performed using two platforms: (1) Integrity Pathway Analysis (IPA; Qiagen) and (2) clusterProfiler with the Kyoto Encyclopedia of Genes and Genomes (KEGG) database [10]. In IPA, significantly differentially expressed genes (DEGs) at 3, 6, 9, and 12 days in 3D cell cultures were analyzed for core pathways. The most significant canonical pathways, functions, and gene networks were identified. Z-scores of enriched canonical pathways were plotted to visualize the dynamic changes in gene expression before, during, and after granuloma development in the 3D model. In a second approach, a list of significant DEGs was subjected to KEGG pathway enrichment using the *gseKEGG* function included in the clusterProfiler package in R. Output of significantly enriched KEGG pathways was plotted as bar graphs (ggplot2 package in R) for samples at four different time points. The interesting KEGG pathways were further identified using the Pathview package in R [11].

GSEA results were presented via bar graphs for both IPA and KEGG, with separate panels for each time point in 3D cell culture. The x-axis of the bar plot represents the z-score for IPA and the normalized enrichment score (NES) for KEGG. Heatmaps of gene expression were generated using the pheatmap.2 function of the gplots package (v3.1.3) in R and Venn diagrams were generated using the online program (Draw Venn Diagram (ugent.be)). Gene expressions in the top interesting pathways were plotted as heatmaps using the pheatmap.2 function of the gplots package (v3.1.3) in R. Shared DEGs identified by Venn diagrams were subjected to GSEA using online software ShinyGo 0.80 (ShinyGO 0.80 (sdstate.edu)) [12].

Additionally, we compared our 3D tuberculoma model with the previously published 3D *in-vitro* granuloma model by Elkington and colleagues [13], which involved encapsulating *Mtb*-infected human PBMCs and collagen ECM in polymer microparticles, and with clinical human TB granulomas at the transcriptomic level. We downloaded raw RNA sequencing reads from their study [13] from the Gene Expression Omnibus (GEO) database, using accession codes GSE174566 (*Mtb*-infected PBMC 3D cell culture model) and GSE174443 (clinical samples of human lymph node TB with granulomas). We also compared data from the study by Khader and colleagues [14] regarding the *Mtb*-infected diversity outbred (DO) mice and macaque models. Raw RNA sequencing reads of mouse and macaque lung samples with tuberculous granulomas from their study [14] were downloaded from the Bioproject PRJNA523820. We also compared 16 signature genes that were differentially expressed between progressors and non-progressors in the *Mtb*-infected South African adolescent cohort that predicted the onset of TB disease across species [14]. Furthermore, we examined 70 signature genes that were consistently upregulated in the blood of active TB patients compared to individuals with latent *Mtb* infection, as described in cohorts from the UK and South Africa by O'Garra and colleagues [15]. The $\log_2$-fold change in gene expression in patients' blood samples (as shown in Table S3 of their published research) [15] was used for the comparative transcriptomic analysis. The raw reads were downloaded from the NCBI GEO database with the accession code GSE107995. All sample identifiers and descriptions are provided in **S9 Table**. Raw FASTQ files were downloaded using the SRA toolkit/2.11.3. All raw FASTQ files underwent the same transcriptomic analysis workflow described in this study (**S10 Fig**). DEGs with absolute $\log_2$ fold-change > 1 and adjusted *p*-value < 0.05 were used for the comparison analysis.

#### Fluorescent probes and microscopy

**(A) Hypoxia Probes:** Hypoxia induction was probed using Image-iT™ Red Hypoxia Reagent (ThermoFisher Scientific) or Hypoxia Red fluorogenic probe (ROS-ID® Hypoxia/Oxidative Stress Detection Kit, Enzo Life Sciences). Image-iT™ Red Hypoxia Reagent is a live cell-permeable fluorogenic compound that fluoresces when atmospheric oxygen levels are as low as 5% and reverses when the cells return to normal oxygen levels. This reagent is considered more sensitive than pimonidazole, which forms adducts in hypoxic cells in granulomas [16] and only responds to very low oxygen levels. It has an excitation/emission maxima of 490/610 nm. The 3D cell cultures of THP-1 monocytes, with or without *Mm* (WT 1218 or 'M') or *Mtb* (WT Erdman), were developed to investigate hypoxia induction. 3D tuberculomas or control spheroids were exposed to pre-titrated Image-iT Hypoxia Red Reagent stock solution at a final concentration of 1−10 μM in 3D cell culture medium and incubated in a cell culture incubator at 37 °C with 5% CO2, 20% $O_2$, and >90% humidity for 1 hour, followed by additional 3.5 hours after reagent exchange with a fresh 3D cell culture medium. Imaging was performed using Cytation-5 with a Texas Red filter. The Enzo Hypoxia Red Detection Reagent is a non-fluorescent or weakly fluorescent aromatic compound containing a nitro (NO2) moiety. Due to the presence of nitro-reductase activity in hypoxic cells, the nitro group is converted in a series of chemical steps to a hydroxylamine (NHOH) and an amino (NH2) group; the original molecule then degrades, releasing the fluorescent probe, which stains hypoxic cells red. For hypoxia staining, the medium was exchanged with fresh 3D cell culture medium. 3D tuberculomas or control spheroids, immersed in 50 μl of medium per well, were exposed to a 50 μl pre-titrated Hypoxia Red probe in 3D cell culture medium and cultured for 24 or 48 hours in a cell culture incubator. After incubation, the 3D cell culture medium in the well was carefully exchanged twice with fresh prewarmed (37 °C) medium to remove the reagent. The working stock was prepared by diluting 7 μl of Hypoxia Red reagent in 21 ml of 3D cell culture medium, which was found to be optimal for staining. The reagent also facilitated the characterization of hypoxia induction in 3D tuberculomas following treatment with the HDT compound. After the treatment period, spheroids were stained with hypoxia probe as described above and cultured for 16−24 hours. The stained spheroids were incubated for 1 hour after media exchange and imaged using Cytation-5 with a Texas Red filter. Hypoxia inducer deferoxamine and hypoxia inhibitor nitazoxanide were used as positive and negative controls, respectively.

**(B) Biofilm staining:** *Mtb* biofilm formation in 3D tuberculomas of THP-1 monocytes and *Mtb* Erdman (WT) was probed using FUN-1 and Calcofluor white (CW) M2R staining kit (ThermoFisher Scientific). FUN-1 is a unique two-color fluorescent probe that readily diffuses into cells and fluoresces in red (Texas red) and green (GFP) channels, while CW labels chitin and cellulose (β (1,4)-D-glucopyranosyl units) with blue fluorescence. To prepare a staining solution, 1 μl of Component A (FUN 1 stain) and 5 μl of Component B (CW stain) were added to each ml of prewarmed (37 °C) 3D cell culture medium, achieving final concentrations of 10 μM for FUN-1 and 25 μM for CW. 3D tuberculomas or control spheroids, immersed in 50 μl medium per well, were exposed to 100 μl of biofilm staining solution and cultured for 1 hour in a cell culture incubator. After incubation, the medium was exchanged with fresh, prewarmed (37 °C) 3D cell culture medium, and the cultures were imaged using Cytation-5 with appropriate filters for brightfield (3D spheroids), Texas Red and GFP (FUN-1), and DAPI (CW).

**(C) Live/dead probes:** To probe cell death, 3D infected and control spheroids were stained using a Fixable Red Dead Cell Stain kit (Thermo Fisher Scientific) according to the manufacturer's instructions. The stain reacts with free amines in the interior and on the surface of cells with compromised membranes, yielding intense red fluorescent staining. The stain was prepared by adding 50 µl of DMSO to the vial and then mixing 1 µl of the reconstituted dye with 1 ml of PBS. For staining, the medium was exchanged with PBS, and 3D tuberculomas or control spheroids in 50 µl of PBS per well were exposed to 100 µl of staining solution and incubated for 30 minutes to 1 hour at room temperature. The spheroids were then carefully washed twice with 100 µl of PBS containing 1% bovine serum albumin (BSA) and resuspended in 100 µl of PBS with 1% BSA before imaging in a Cytation-5 using a Texas Red filter. Additional live/dead cell staining was performed using a Calcein AM and BOBO-3 iodide kit (Thermo Fisher Scientific) according to the manufacturer's instructions. The presence of intracellular esterase activity distinguishes live cells. Esterase staining with Calcein (green) indicates live cells, while BOBO-3 iodide (red) marks free DNA or dead/damaged cells in the 3D cultures.

**(D) Apoptosis and necrosis probes:** The apoptotic and necrotic cell staining was performed using the Apoptosis/Necrosis kit (Abcam), which includes Apopxin (red) for detecting apoptotic cells via phosphatidylserine (PS), Nuclear Green DCS1 (green), a membrane-impermeable dye, for staining nuclei of damaged or necrotic cells, and CytoCalcein dye (blue/violet) for live cells. The cell culture medium in wells with 3D tuberculomas or control spheroids was carefully exchanged with 100 µl assay buffer. After aspirating the assay buffer, the spheroids were resuspended in 200 µl of fresh assay buffer containing 2 µl of Apopxin Deep Red Indicator (100×), 1 µl of Nuclear Green (200×), and 1 µl of CytoCalcein 450 (200×). Following incubation for 1 hour at room temperature, spheroids were washed twice with 100 µl of assay buffer, resuspended in fresh assay buffer, and imaged using a Cytation-5 with Texas Red, GFP, and DAPI filters.

**(E) Lysosomal acidification probe:** Lysosomal acidification was probed using a unique cationic amphiphilic tracer (CAT) dye, part of the LYSO-ID® Red cytotoxicity kit (Enzo Life Sciences), which rapidly partitions into cells and labels acidic organelles (red) and enables the long-term monitoring of lysosome and lysosome-like organelle accumulation and cytotoxic effects resulting from drug treatment. The dye fluoresces in acidic environments generated by increased cell death and the release of acidic organelles. Compounds causing phospholipidosis or the accumulation of autophagosomes by blocking the downstream lysosomal pathways and intracellular trafficking of autophagosomes also lead to increased intracellular LYSO-ID® Red fluorescence, indicating probe accumulation. The red fluorescent probe is selectively sequestered in acidic organelles by a mechanism involving protonation and retention within the organelles. 3D spheroids treated overnight with a final concentration of 100 μM verapamil in the medium were used as a positive control. The medium was carefully aspirated for 3D tuberculomas and control spheroids staining, and 100 μl of 1× assay buffer was dispensed to each well. Likewise, after incubating 3D tuberculomas with the compound of interest, the medium was carefully aspirated, and 100 μl of 1× assay buffer was dispensed. The buffer was carefully aspirated, and 100 μl of the 1× Dual Color Detection Reagent was dispensed to each well. The detection reagent was prepared by diluting 1 ml of 10× Dual Color Detection Reagent in 8.8 ml of detection buffer and 0.2 ml of FBS. The spheroids were incubated in the dark at room temperature for 30 minutes to 1 hour and washed twice with 200 μl of 1× assay buffer. Excess buffer was aspirated, and 50 μl of fresh 1× assay buffer was added to each well. Red lysosome staining was imaged using Cytation-5 with a Texas Red filter, and blue nuclear counterstain was imaged with a DAPI filter. Increases in red lysosome signal, without significant loss of blue signal, indicate probe accumulation within cells, resulting from an increase in lysosome or lysosome-like vesicle size and number.

#### Autophagy assays

Autophagy induction in 3D *in-vitro* tuberculomas was investigated using 3D co-cultures of THP-1-Difluo hLC3 reporter cells [17] and *Mtb* Erdman (WT). Control groups included untreated, DMSO-treated, rapamycin-treated (an mTOR inhibitor and autophagy inducer), chloroquine-treated (a lysosomal inhibitor), and rapamycin plus chloroquine-treated tuberculomas. Notably, while chloroquine can inhibit autophagy by blocking autophagosome-lysosome fusion and slowing down lysosomal acidification, chloroquine-induced lysosomal inhibition can also inhibit mTORC1 and secondarily induce autophagy. It can also induce LC3-II formation independently of autophagy.

For secondary autophagy measure, THP-1 monocytes were transduced with the Premo Autophagy Sensor LC3B-RFP (BacMam 2.0, Invitrogen) in a T75 flask following the kit supplier's protocol. BacMam technology utilizes a modified baculovirus (an insect virus) to deliver genes into mammalian cells efficiently, ensuring safety and minimizing cytopathic effects. BacMam 2.0

incorporates a pseudotyped capsid protein for more efficient cell entry and genetic elements (enhanced CMV promoter and Woodchuck Post-transcriptional Regulatory Element) that boost expression levels. After 16 hours of transduction, THP-1 cells ($1\times10^5$ cells) were infected with WT *Mm* M (MOI 0.008) in flat clear-bottom black 96-well 2D cell culture plates (Costar, 3603) in complete RPMI-1640 medium without antibiotics (200 µl/well) and incubated at 37°C with 5% CO2 and 100% humidity. On day 6 post-infection, 2D co-cultures were exposed to selected individual compounds at 20 µM, $CC_{50}$, $EC_{50}$, or 2-fold dilutions. Cultures treated with DMSO at equal v/v in the medium or medium alone served as controls. 2D cultures were imaged at 20× on day three post-treatment using a Cytation-5 cell imager.

Additionally, autophagy was investigated using the Enzo Life Sciences CYTO-ID® Autophagy Detection Kit. The kit measures autophagic vacuoles and monitors autophagic flux in lysosomally-inhibited live cells using a green dye that selectively labels accumulated autophagic vacuoles. The 488 nm-excitable dye has been optimized through the identification of titratable functional moieties that allow for minimal staining of lysosomes while exhibiting bright fluorescence (green) upon incorporation into pre-autophagosomes, autophagosomes, and autophagolysosomes. The kit also includes the Hoechst 33342 dye for nuclear staining, an autophagy inducer (rapamycin), and a lysosomal inhibitor (chloroquine). To monitor autophagy, co-cultures of THP-1 monocytes and WT *Mm* 'M' were exposed to candidate molecules or their serial dilutions on day six post-co-culture. However, we found that several chemical compounds induce autofluorescence in the GFP channel in 3D tuberculomas or 2D cell cultures, which interfered with autophagy results using this dye and rendered the kit unsuitable for autophagy studies of autofluorescing compounds. Consequently, those results were not presented.

#### Screening assay statistics and validation

A chemical compound screening assay using *Mm* or *Mtb* (tdTomato)-infected human PBMC or THP-1 monocyte 3D tuberculoma bioplatform was developed following HTS assay guidelines [18, 19]. The heat map generated using Gen-5 v3.08 software in Cytation-5 was used to assess the reproducibility of fluorescent intensity data between individual replicate wells in 3D cell culture microplates (Corning 3415). No outliers were discarded during data analysis. The normalized reduction of bacterial burden (%) in test compound-treated (n = 3) or DMSO-treated (n = 3–6) individual wells was calculated using the formula below, where $F_t$ is the fluorescence intensity of test compound- or DMSO-treated wells. Mean $F_u$ is the average fluorescence intensity of untreated tuberculoma wells (n = 6).

$$\text{Percent bacterial burden reduction} = \frac{(F_t - \text{Mean } F_u)}{\text{Mean } F_u}$$

The normalized bacterial burden reduction (%) in compound-treated wells was compared to that in DMSO-treated control wells. Statistical significance was determined using the Kruskal-Wallis test and Dunn's post hoc test. The Z'-factor statistic assessed the impact of the MOI of mycobacterial strains (tdTomato) on the screening assay quality in the THP-1 tuberculoma platform. The Z'-factor describes how well separated the positive and negative controls are and indicates the probability of false positives or negatives. The assay was investigated using assay controls without the intervention of test compounds in a 96-well plate format. HTS assay optimization requires performance evaluation in multiple plates and determination of plate-to-plate variation. Therefore, the Z'-factor for the screening assay was calculated by performing the assay in a batch of 10 microplates and using the formula below.

$$Z' = 1 - \frac{(3\sigma_{c-} + 3\sigma_{c+})}{|\mu_{c-} - \mu_{c+}|}$$

In the Z'-factor formula, $\sigma_{c-}$ is the standard deviation of the untreated or DMSO control wells, $\sigma_{c+}$ is the standard deviation of the rifampicin or nitazoxanide control wells, $\mu_{c-}$ is the mean of the untreated or DMSO control wells, and $\mu_{c+}$ is the mean of the rifampicin or nitazoxanide control wells. A Z' value between 0.5 and 1 is considered excellent, a value between 0 and 0.5 is acceptable, and a value less than 0 indicates that the assay is unlikely to be suitable for HTS applications.

The data from screenings of compounds in the 3D tuberculoma bioplatform, investigating the effects on *Mm* and *Mtb* (tdTomato) strain burdens, were also analyzed using z-scores. The z-score was calculated per plate and for each well using the formula below.

$$z = \frac{(x - \mu)}{\sigma}$$

Where x = the fluorescent intensity of the compound treated well, μ = the mean of the fluorescent intensity of the negative control wells, and σ = the standard deviation of the fluorescent intensity of the negative control wells. The average z-score and standard error for each compound screened in 2–4 experiments per mycobacterial strain are presented. A robust z-score cutoff of −4 was used.

The uniformity and reproducibility of 3D tuberculomas generated in the platform were also determined. The platform generated highly uniform 3D tuberculomas and exhibited consistent screening assay results, yielding excellent to acceptable Z'-factors. A detailed step-by-step protocol for chemical compound screening using the 3D tuberculoma bioplatform, data processing, analyses, and suitability for HTS applications is described in the Methods article [20].

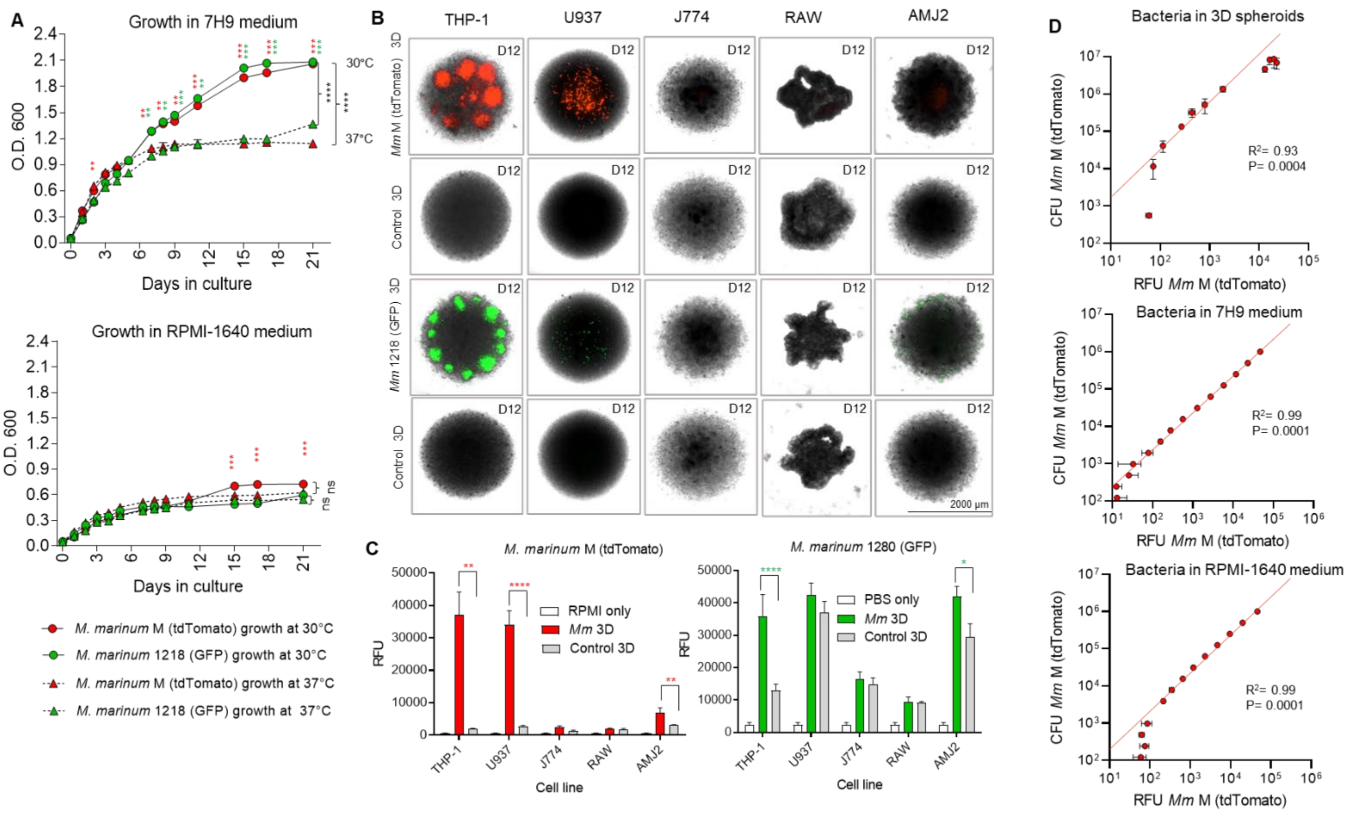


**S1 Fig. 'Mycobacteria-in-spheroid' 3D co-cultures using human or mouse macrophage cell lines and fluorescent *M. marinum* strains.**

(**A**) Growth of *Mm* 'M' (tdTomato) and 1218 (GFP) strains in Middlebrook 7H9 or RPMI-1640 medium at 30 or 37 °C measured by spectrophotometry. Data are the optical density (O.D.) at 600 nm of axenic cultures (n = 3) grown in 5 ml medium in culture tubes and initiated with a 100 µl inoculum ($1\times10^7$ CFU/ml) from frozen stocks. Error bars indicate SD. **$p < 0.01$, ***$p < 0.001$, and ****$p < 0.0001$ comparing growths at two temperatures by 2-way ANOVA with Tukey's test for growth curves (black asterisks) or Welch's *t*-test for indicated time points (colored asterisks). (**B** and **C**) Growth of *Mm* 'M' (tdTomato) and 1218 (GFP) in 3D co-cultures of monocyte-macrophage cell lines at 37 °C in RPMI-1640 medium in 96-well 3D cell culture microplates (Corning 4515). Five different cell lines, THP-1 monocytes (human leukemia), U937 pleural-fluid macrophages (human histiocytic-lymphoma), J774A.1 ascites macrophages (BALB/c mouse reticulum-cell-sarcoma), RAW-264.7 macrophages (BALB/c mouse leukemia virus-transformed), and AMJ2-C11 alveolar macrophages (C57BL/6J mouse, *in vitro* J2 retrovirus-transformed), were investigated. (**B**) Representative images of *Mm*-infected and uninfected control 3D spheroids at day 12. (**C**) Fluorescence intensity, measured as relative fluorescent units (RFU), demonstrated background autofluorescence in uninfected spheroids in the GFP channel despite exchanging phenol red RPMI-1640 medium with PBS (pH 7.2). Data in **B** and **C** are of 5–8 infected and four control spheroids per cell line. Error bars indicate SD. *$p < 0.05$, **$p < 0.01$, and ****$p < 0.0001$ using Brown-Forsythe and Welch ANOVA multiple comparison tests. (**D**) A relationship between the number of red-fluorescent bacteria and RFUs in the 3D spheroids, 7H9 medium, or RPMI 1640 medium. n = 6. Error bars indicate SD. Pearson correlation coefficients $R^2$ and two-tailed *p*-values are shown.

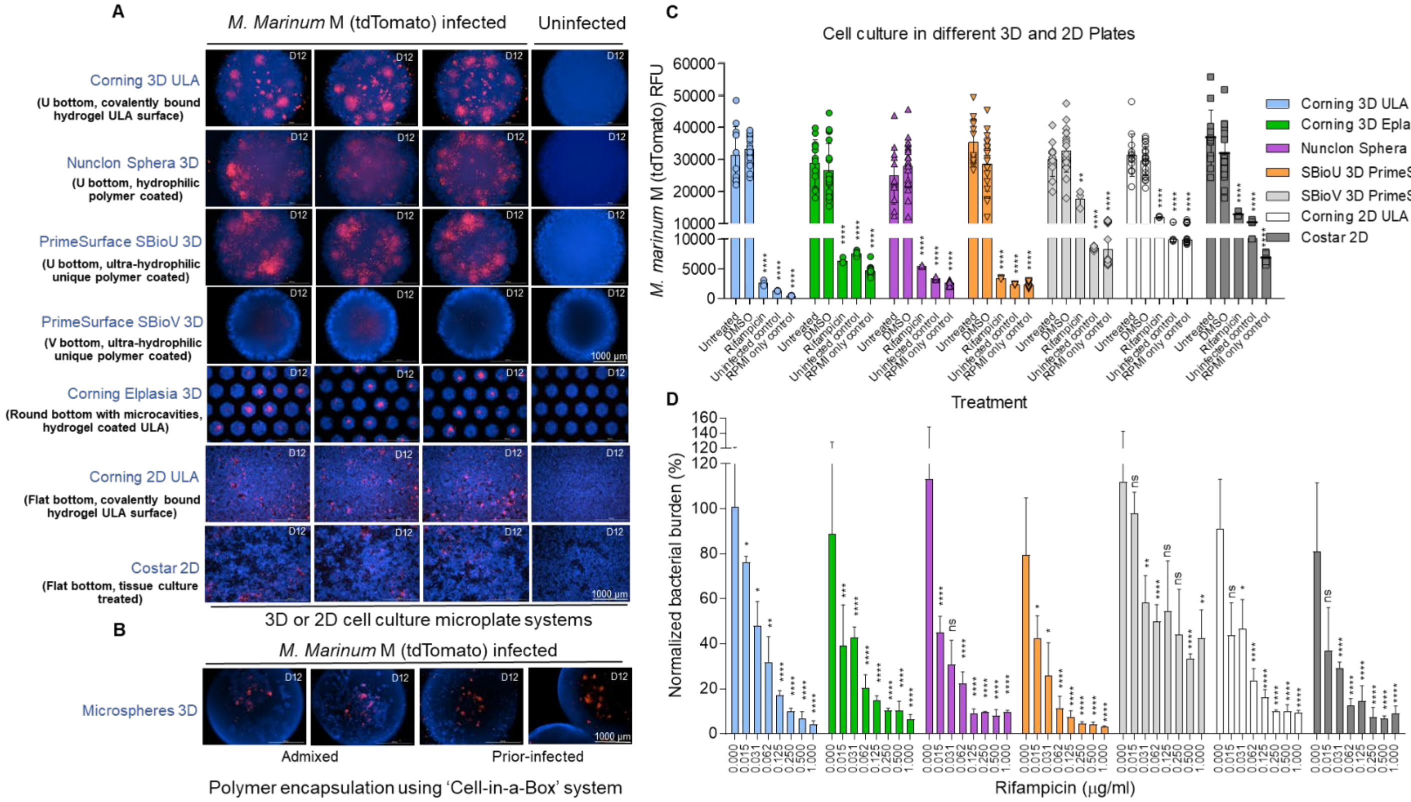


**S2 Fig. Generating tuberculoma-like structures with conglomerating lesions in the co-cultures depends on the cell-culture-ware and growth conditions.**

**(A)** THP-1 monocyte culture with or without *Mm* 'M' (tdTomato) in 96-well 2D and 3D cell culture plates with distinct microwell geometries and cell-culture surfaces. (**B**) THP-1 monocyte and *Mm* 'M' (tdTomato) co-culture in a polymer encapsulated 'Cell-in-a-Box®' 3D system. Representative images show the formation of miniature granulomatous aggregates rather than well-organized lesions in the encapsulated 3D system, regardless of whether THP-1 cells were admixed or infected overnight with bacteria before encapsulation in cellulose polymer particles. (**C**) The fluorescence intensity readings in the 2D and 3D cell culture plates. In all plate types, a significant reduction in fluorescence was found in rifampicin (1 µg/ml)-treated co-cultures relative to untreated or DMSO-treated controls. However, the lowest background noise was found in control wells containing RPMI-1640 medium only or uninfected THP-1 cultures using a Corning 3D ULA plate. RFU, relative fluorescence units. Data are from 3–21 replicates per treatment or control. (**D**) The bacterial burden reduction in different 2D and 3D cell culture plates. The results show a gradual reduction in bacterial burdens with increasing rifampicin dose in the Corning 3D ULA plate. Data are from 21 control (0 µg/ml) and three rifampicin-treated cultures per dose. Error bars in **C** and **D** indicate SD. $*p < 0.05$, $**p < 0.01$, $***p < 0.001$, and $****p < 0.0001$ using Brown-Forsythe and Welch ANOVA multiple comparison tests. The low MOI (0.008) was used in all experiments, treatments were carried out on day 6, and representative images (**A** and **B)** and fluorescence intensity readings (**C** and **D**) were captured on day 12 post-co-culture. Images (in **A** and **B**) represent 6–12 infected or uninfected control cultures per plate or system type.

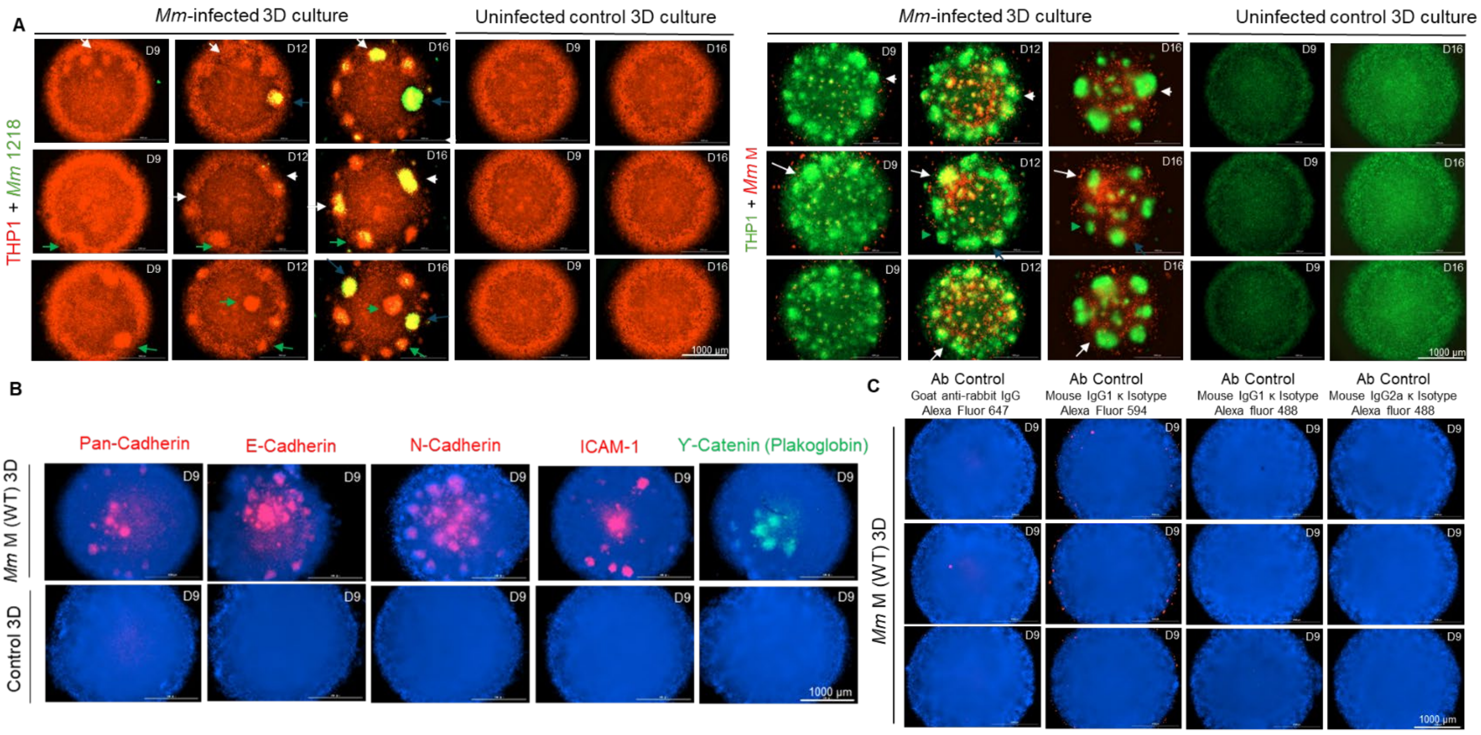


**S3 Fig. Dynamic granuloma lesions develop in the 3D model and express characteristic markers.**

**(A)** Spatiotemporal host and bacterial growth dynamics and granulomatous lesion development in the 3D co-cultures generated using either THP-1-RFP monocytes with *Mm* 1218 GFP or THP-1-copGFP monocytes with *Mm* 'M' tdTomato. Examples of coalescing granulomatous lesions over time (white arrows), lesions that progressed quickly (blue arrows), and those that remained relatively stable (green arrows) are indicated using arrowheads. The images shown are representative of 54 infected spheroids and 6 control spheroids. (**B**) The macrophage epithelization and adherence junction markers Pan-cadherin, E-cadherin, N-cadherin, ICAM-1 (all red), and γ-catenin (green), detected in the nascent granulomatous aggregates in the 3D co-cultures of THP-1 cells and WT *Mm* 'M'. The cell surface cadherins, catenins, and ICAM-1 are known to be shed or secreted and can potentially accumulate in the necrotic cores over time. Whole-mount immunostaining was performed using titrated antibodies on day 9 in the co-cultures generated with a very low MOI (0.004), to minimize mycobacterium-induced extensive cell death. These co-cultures contained fewer early developing granulomatous aggregates, with some forming in the centers of the structures. (**C**) 3D co-cultures stained with control antibodies (Ab) demonstrating the absence of non-specific staining in the developing granuloma lesions. Representative images (**B** and **C**) are from one of the two experiments performed with three technical replicates.

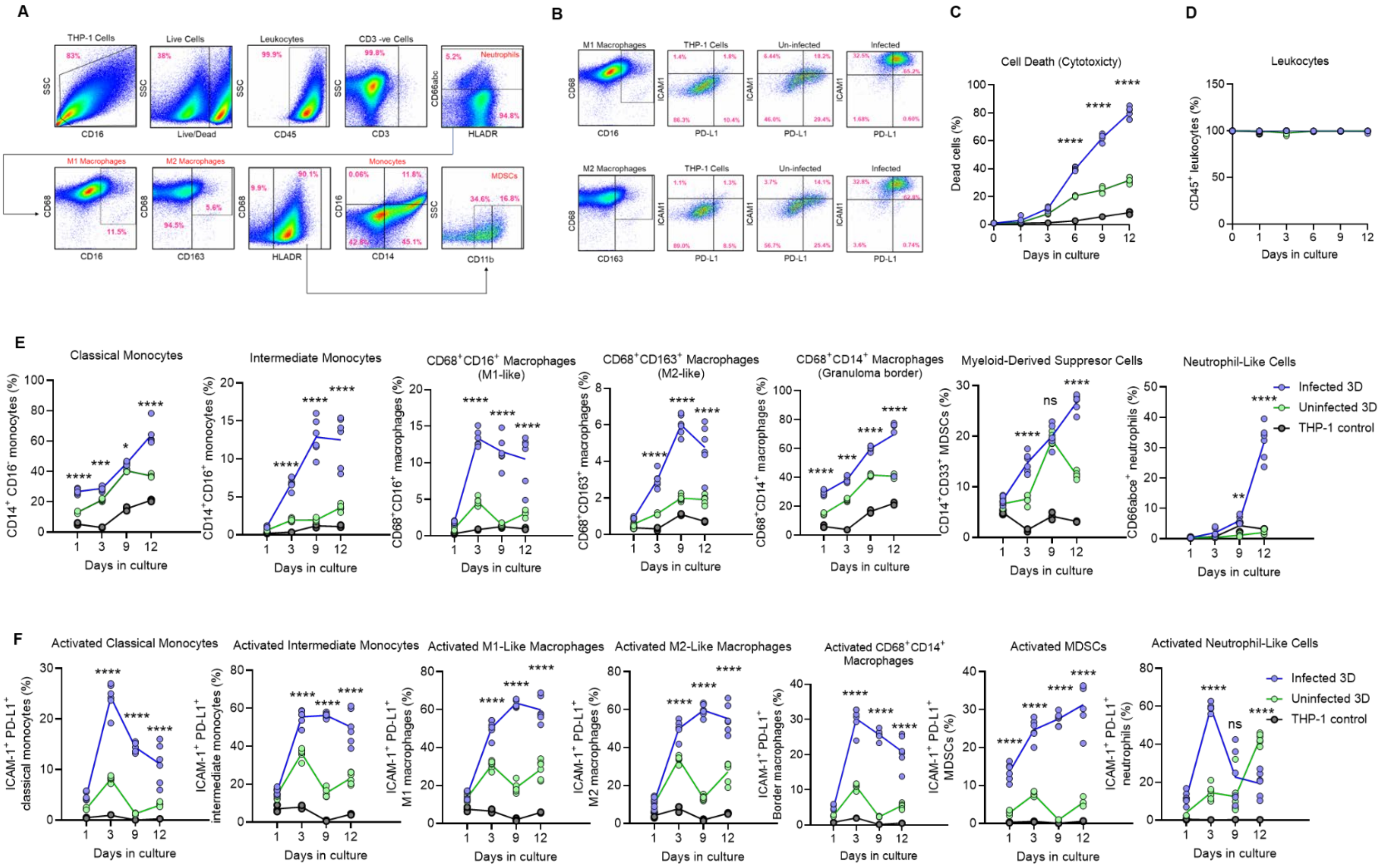

**S4 Fig. Myeloid cells in the 3D model express distinct cell subset markers**

**(A)** The multicolor flow cytometry dot plots of one representative 3D *Mm* 'M'-infected cell culture sample (day 9), illustrating the gating strategy to identify different cell subsets. (**B**) The dot plots of one representative THP-1 control (2D suspension), 3D uninfected control, and 3D *Mm* 'M'-infected sample (day 9), illustrating the gating strategy to identify ICAM$^{+}$PD-L1$^{+}$ M1 and M2-like macrophages. (**C**) The magnitude of dead cells detected in three cell culture types using fixable live/dead cell stain. (**D**) The frequencies of CD45$^{+}$ leukocytes in three cell culture types over 12 days. (**E**) The magnitude of different cell subsets in 3D-infected, 3D-uninfected controls, and THP-1 controls. (**F**) The magnitude of ICAM$^{+}$PD-L1$^{+}$ activated cell subsets in 3D infected, 3D-uninfected controls, and THP-1 controls. One 3D infected or uninfected sample per time point consisted of cells pooled from six spheroids. n = 6 infected and uninfected samples collected from six individual 3D cell culture plates and n = 3 THP-1 control samples (2D suspension culture grown in tissue culture flasks). $^{*}p < 0.05$, $^{**}p < 0.01$, $^{***}p < 0.001$, $^{****}p < 0.0001$ using two-way ANOVA with Šidák's multiple comparison test.

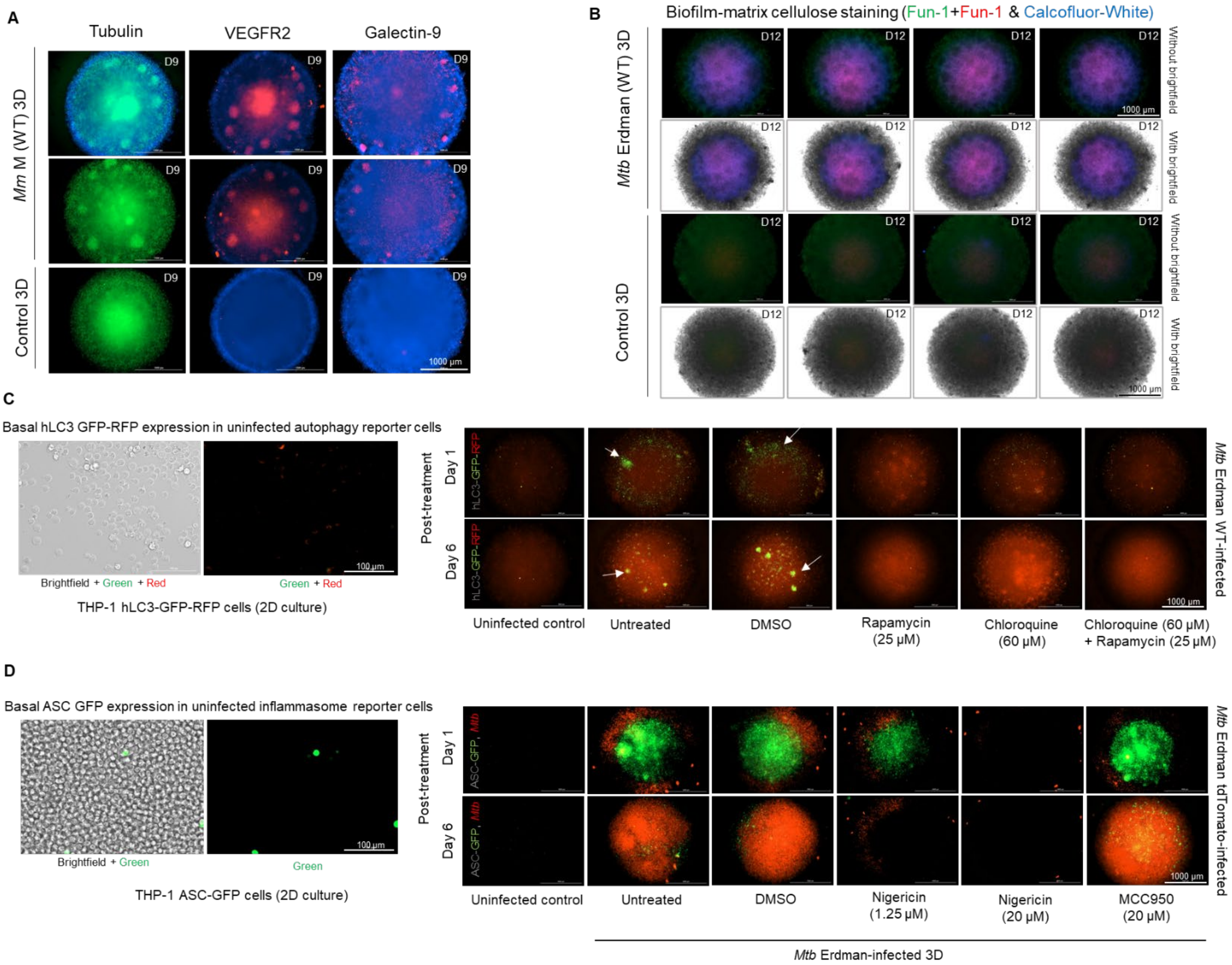


**S5 Fig. The 3D model expresses key features and microenvironments within human tuberculomas, including mycobacterial biofilm formation.**

(**A**) Expression of α-tubulin (green), VEGFR-2, and galectin-9 (red) determined by immunostaining in the 3D tuberculomas of THP-1 monocytes and *Mm* 'M' WT and uninfected spheroids. Nuclei are stained using Hoechst 33342 dye (blue). The α-tubulin is conserved in eukaryotic cells, while VEGFR-2 and galectin-9 are known to be shed and secreted from cells and may accumulate in the necrotic core over time. Representative images are from one of the two experiments, with three technical replicates, and show early developing granulomas on day 9 (MOI 0.004). (**B**) *Mtb*-biofilm formation in the 3D tuberculomas of THP-1 monocytes and *Mtb* Erdman (WT) as detected by FUN-1 and Calcofluor white (CW) staining of biofilm-matrix component cellulose. Composite images from red and green (Fun-1) and blue (CW) fluorescence channels, overlaid with or without one from the brightfield channel, are shown. Except for a diffuse green fluorescence (background), no fluorescence was detected in the uninfected 3D spheroids. Images shown are representatives of at least 6 infected and control spheroids each. (**C**) 3D tuberculomas of THP-1-Difluo hLC3 autophagy reporter cells infected with *Mtb* Erdman (WT). The left panel shows the basal expression of hLC3-GFP-RFP in the uninfected reporter cells grown in the Corning ULA flat-bottom microwell as a 2D cell culture. Uninfected 3D cell cultures and co-cultures treated with DMSO (drug carrier), rapamycin (mTOR inhibitor and autophagy inducer), chloroquine (lysosomal inhibitor), and rapamycin plus chloroquine are shown on the right. Untreated or DMSO-treated 3D co-cultures exhibit accumulation of yellow-green puncta (white arrows) in the granuloma-zone macrophages and numerous cellular clusters on day 1 and day 6 post-treatment (days 7 and 13 post-infection), indicating a block in autophagolysosome formation and incomplete autophagy flux. Increased red fluorescence on day 6 post-treatment indicates delayed induction of autophagy (*Mtb* infection- and starvation-induced) in these 3D co-cultures. On the contrary, rapamycin-treated 3D co-cultures exhibit relatively rapid induction of autophagy (red fluorescence) throughout the 3D co-culture (≤ 1-day post-treatment). Chloroquine-treated co-cultures show increased autophagy on day 6 compared to day 1. Images are representative of 3–6 spheroids per treatment.

(**D**) Inflammasome and pyroptosis induction in the 3D tuberculomas of THP-1-ASC-GFP reporter cells infected with *Mtb* Erdman (tdTomato). *Mtb* infection leads to ASC-GFP expression and ASC-speck formation following inflammasome activation in these reporter cells. The left panel shows the basal expression of ASC-GFP in the uninfected inflammasome reporter cells grown in Corning ULA flat-bottom microwells as a 2D cell culture. An NF-κB-inducible promoter drives the ASC-GFP expression, and little GFP signal is detected in resting cells. Uninfected 3D cultures and 3D co-cultures treated with DMSO (drug carrier), nigericin (NLRP3 inflammasome inducer), and MCC950 (NLRP3 inhibitor) are shown on the right. Untreated or DMSO-treated 3D co-cultures demonstrate that *Mtb* (red) induces the assembly of ASC-dependent inflammasomes (green) in THP-1 cells and, eventually, cell death. Nigericin-treated 3D co-cultures exhibit ASC-speck (green) formation, relatively rapid macrophage death (pyroptosis), and inhibition of *Mtb* (red) growth. Conversely, MCC950 treatment, which is known to inhibit inflammasomes and ASC speck formation, fails to reduce the *Mtb* burden in 3D co-cultures by day 6 post-treatment, relative to DMSO. Images are representative of 3–6 spheroids per treatment. *Mtb* MOI 0.05 (**B**–**D**).

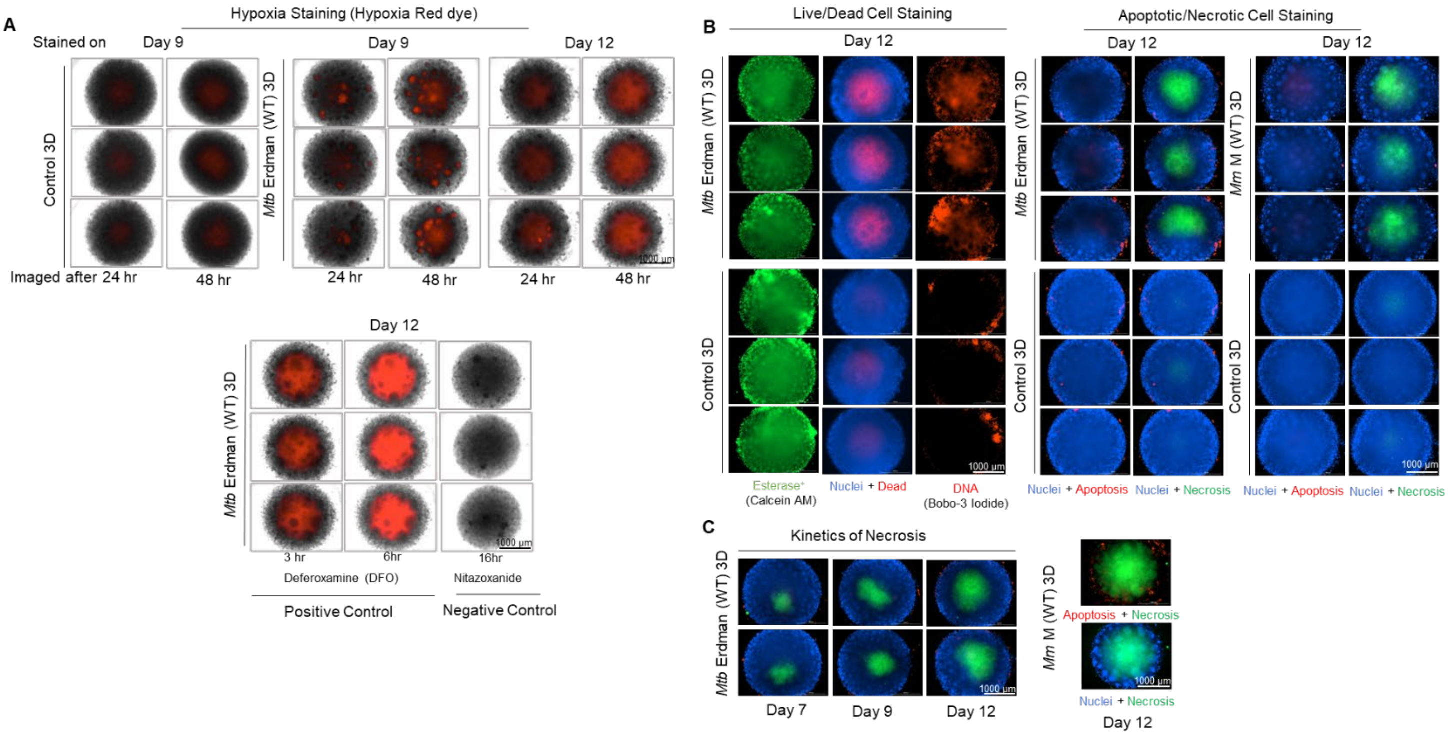


**S6 Fig. Hypoxia and necrosis develop in the 3D tuberculomas.**
(**A**) Increased hypoxia staining in the centers of 3D tuberculomas of THP-1 monocytes and WT *Mtb* Erdman (MOI 0.05). Images are representative of 3D infected and control spheroids exposed to a fluorogenic Hypoxia Red probe (Enzo Life Sciences) at indicated time points and cultured for an additional 24 or 48 hours in the $CO_2$ incubator before image capture. Representative images of 3D tuberculomas exposed to hypoxia inducer deferoxamine (200 µM for 3 or 6 hours) and hypoxia inhibitor nitazoxanide (20 µM for 16 hours) are presented as positive and negative controls, respectively. (**B**) Increased cell death (red) in the centers of 3D tuberculomas compared to control 3D spheroids. Representative images of 3D infected and control spheroids stained using a Fixable Red Dead Cell Stain Kit (Thermo Fisher Scientific) are shown (See middle column in the 'Live/Dead Cell Staining' panel). Nuclei were counterstained using Hoechst 33342 dye (blue). As a second measure of live/dead cell staining, representative images of 3D infected and control spheroids stained using a Live/Dead Cell Imaging Kit containing esterase and BOBO-3 iodide are also shown. Esterase staining (green) indicates live cells, and BOBO-3 iodide (red) staining denotes free DNA or dead/damaged cells. The 'Apoptotic/Necrotic Cell Staining' panel includes representative images of 3D infected and control spheroids following apoptotic and necrotic cell staining using an Abcam Apoptosis/Necrosis Kit. Images reveal increased cell death in the centers of 3D-infected spheroids compared to control spheroids due to increased macrophage necrosis (green) rather than apoptosis (red). Apopxin, a phosphatidylserine (PS) sensor in the kit, stains apoptotic cells red. Nuclear Green DCS1, a membrane-impermeable dye, stains the nucleus of damaged cells or those undergoing necrosis green, and CytoCalcein dye stains live cells blue/violet. (**C**) Longitudinal imaging performed shows a gradual increase in cell necrosis in the cores of 3D tuberculomas over a 12-day period. (**A**–**C**) Images are from one of the two experiments, which contained 3–6 replicates.

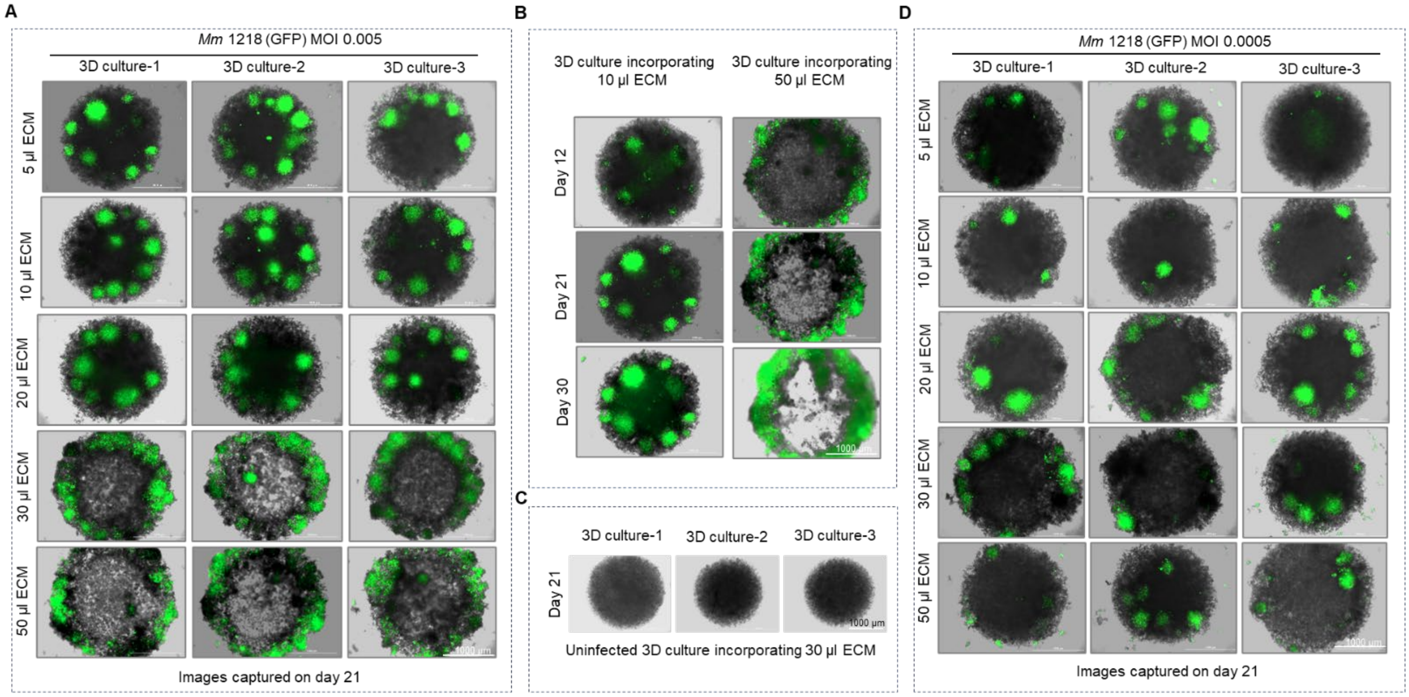


**S7 Fig. Cavity formation occurs following increased ECM deposition in the 3D tuberculomas.**

(**A**) Development of cavity-like features on day 21 in the 3D co-cultures of THP-1 monocytes and *Mm* 1218 GFP (MOI 0.005) following a deposition of an increasing amount (5−50 µl) of the ECM mixture containing human collagen type-1 and fibronectin. (**B**) The kinetics of cavity formation in the 3D co-cultures of THP-1 monocytes and *Mm* 1218 GFP (MOI 0.005) deposited with 10 or 50 µl of ECM are shown. Images show that cavitary transformation starts around day 12, and features grow over time in the co-cultures deposited with 50 µl of ECM. (**C**) Absence of cavitary transformation in the uninfected control 3D spheroids despite the deposition of a high dose of ECM. Representative 3D cell cultures, deposited with 30 µl of ECM, are shown. (**D**) Despite the deposition of a high dose (≥30 µl) of ECM, the absence of cavitary transformation in the 3D co-cultures generated using a 10-fold lower MOI (0.0005) of *Mm* 1218 GFP, which develops lesser bacterial and granuloma burdens. Images are from one of the two experiments performed with 3–6 technical replicates. The day of image capture following 3D cell culture is indicated in each panel.

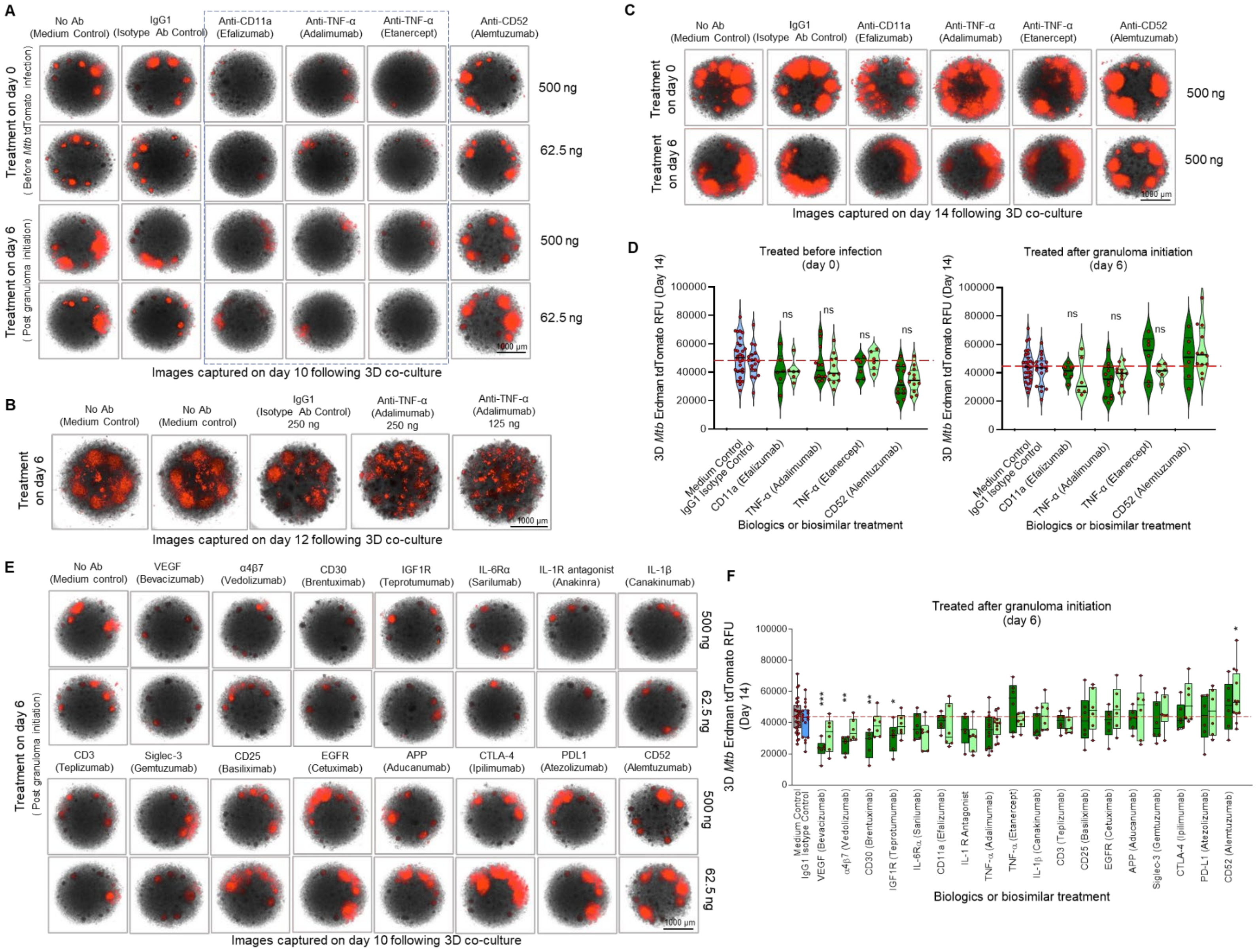


**S8 Fig. Screenings of biologics and biosimilars in the 3D model identify immunotherapeutics that influence the structural organization and bacterial burdens in developing granulomas.** (**A**) Representative images of 3D co-cultures of THP-1 and *Mtb* Erdman (tdTomato) (MOI 0.01) show a loss of structural integrity in nascent granulomas by day 10, after anti-CD11a or anti-TNF but not anti-CD52 or isotype Ab treatments, regardless of the dose (62.5 or 500 ng) or timing of treatment (day 0 or 6) used. The names of clinical counterparts of humanized mAb biosimilars or biologics used are shown in brackets. (**B**) Representative images of 3D co-cultures of THP-1 and *Mm* M (tdTomato) (MOI 0.006) show the formation of numerous miniature (miliary-type) rather than well-organized and compact granulomas by day 12, after anti-TNF adalimumab biosimilar treatment. Despite early interference with the granuloma's structural integrity, anti-CD11a or anti-TNF-treated 3D co-cultures of THP-1 cells and *Mtb* Erdman (tdTomato) eventually form mature granulomas (**C**) and develop bacterial loads (**D**) comparable to those of isotype Ab-treated or medium-alone (no Ab) controls by day 14. ns, nonsignificant using one-way ANOVA after comparing RFU levels (bacterial burdens) in anti-CD11a or anti-TNF-treated co-cultures with isotype Ab-treated or medium alone (no Ab) controls. (**E**) Effects of biologics and biosimilars (n = 15), other than anti-CD11a or anti-TNF, on the structural organization of nascent granulomas in the 3D co-cultures of THP-1 and *Mtb* Erdman (tdTomato) by day 10. (**F**) Effects of biologics and biosimilars (n = 18) on the bacterial burdens in the 3D co-cultures of THP-1 and *Mtb* Erdman (tdTomato) as measured on day 14. $^{*}p < 0.05$, $^{**}p < 0.01$, $^{***}p < 0.001$ using one-way ANOVA with Šidák's post-hoc test. Representative images and data are from one of two experiments performed, containing 6 to 30 replicates. Symbols (red circles) in plots (**D** and **F**) depict the number of 3D co-cultures evaluated per treatment, while dark and light green columns indicate treatments with 500 ng and 62.5 ng, respectively. IgG1 isotype antibody control data is using 500 ng.

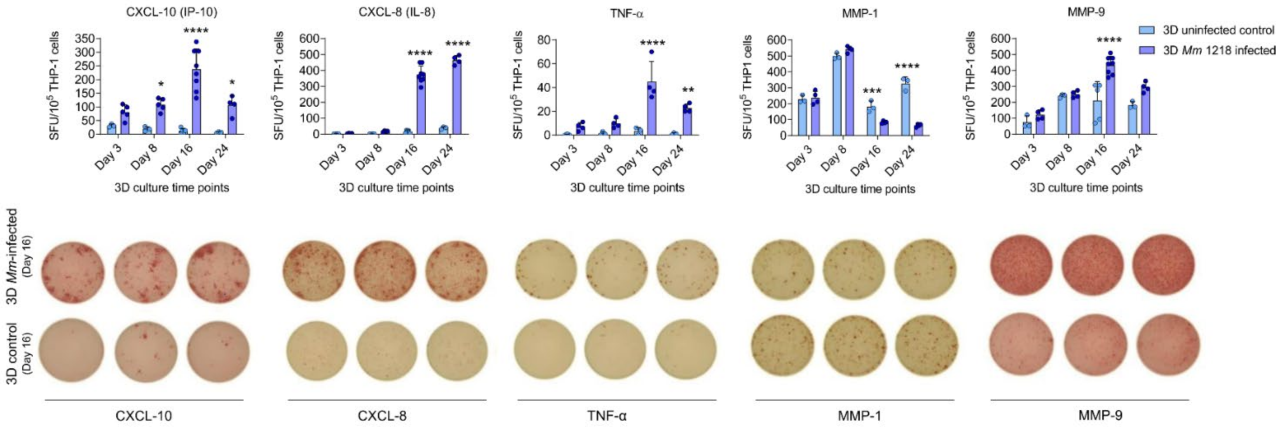


**S9 Fig. Selected immunological protein biomarker-secreting cells in the 3D tuberculomas.** The magnitudes of five selected chemokine, cytokine, or matrix metalloproteinase (MMP)-secreting cells in 3D cell cultures of THP-1 monocytes with or without *Mm* 1218 infection as determined using the enzyme-linked immuno-spot (ELISPOT) assay. The spot-forming units (SFU) per $10^5$ live cells are plotted, and representative images of triplicate wells from the ELISPOT assay using cells isolated on day 16 are shown. Data are from one of the two experiments performed. Cells isolated from 30 infected or control spheroids from the 3D cell culture plate were combined and used in the assay. If necessary, spheroids from multiple plates were combined, and $1\times10^5$ live cells per ELISPOT well were added (n = 3–8 wells/time point). *$p < 0.05$, **$p < 0.01$, ***$p < 0.001$, ****$p < 0.0001$ using one-way ANOVA with Šidák's multiple comparison test. CXCL-10, C-X-C motif chemokine ligand 10 or IP-10, interferon-γ-induced protein 10; CXCL-8, C-X-C motif chemokine ligand 8 or IL-8, interleukin 8; TNF-α, tumor necrosis factor-alpha, MMP-1, matrix metalloproteinase 1; and MMP-9, matrix metalloproteinase 9.

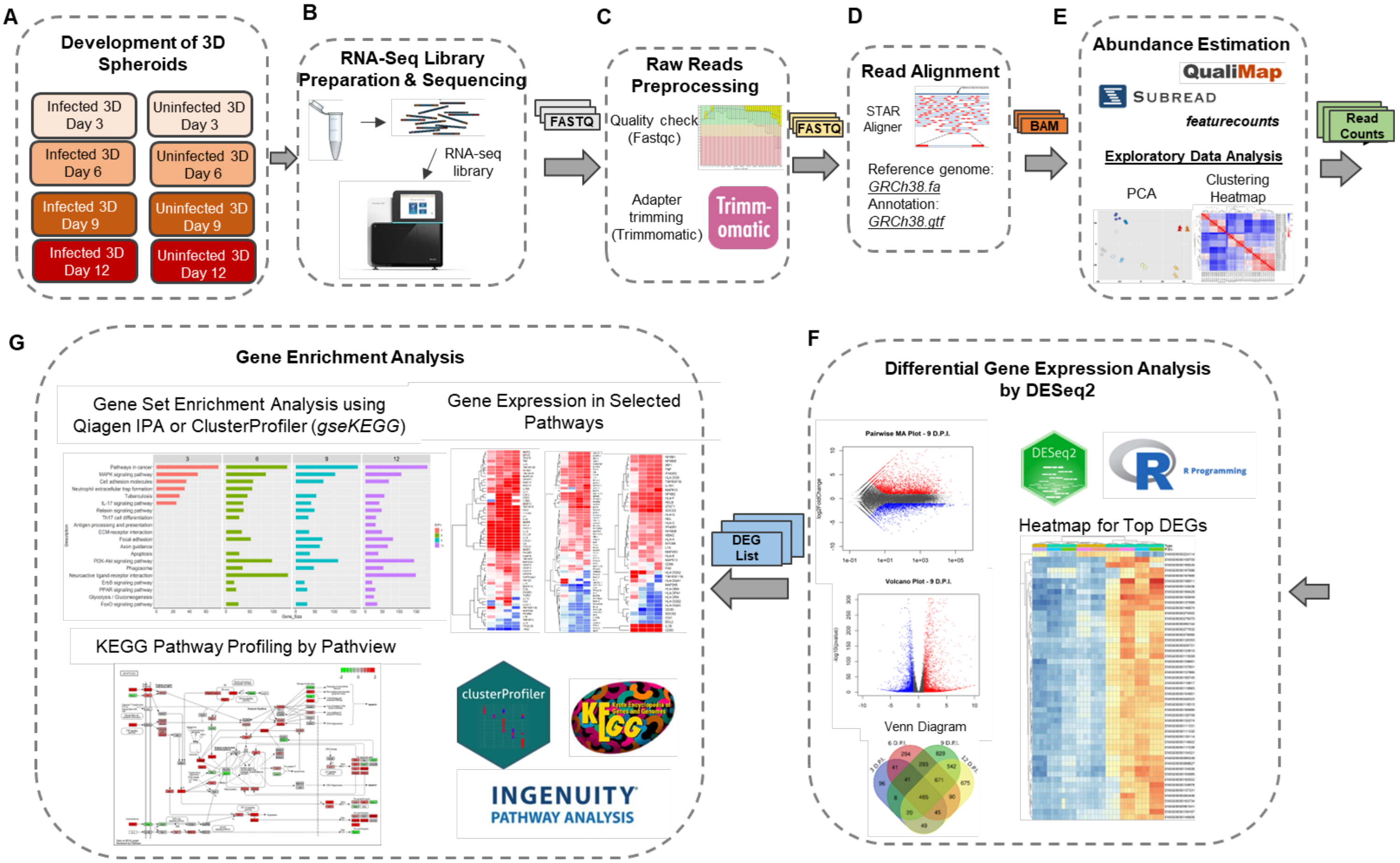


**S10 Fig. The schematic workflow of the transcriptomic characterization of the 3D tuberculoma model.**

(**A**) Development of 3D cell cultures of THP-1 monocytes with or without *Mm* 'M' tdTomato infection (MOI 0.005), and the extraction of total RNA from THP-1 cells on days 0, 3, 6, 9, and 12. (**B**) Preparation of RNA sequencing libraries using Illumina TruSeq Stranded mRNA Library Preparation Kit, and sequencing of the resulting libraries using Illumina NextSeq Sequencing Kit (75 × 2 cycles). (**C**) Quality-check of the RNA-sequencing data using FastQC (v.0.11.5) and trimming using Trimmomatic (v.0.39). (**D**) Mapping of the high-quality reads to the human reference genome (GRCh38.p13) using the STAR aligner software (v.2.5.2b) and annotation using the GRCh38.gtf file. (**E**) Counting of the reads mapped to the genes using the featureCounts function in the Rsubread package in R. Normalization of the generated read count table based on the sequencing depth using the DESeq2 package. PCA plot and generation of the sample distance matrix to visualize the clustering of samples. (**F**) The identification of DEGs using the DESeq2 package and visualization by MA and volcano plots. Determination of core DEGs shared by different time points in 3D cell culture using a Venn diagram. (**G**) The GSEA using IPA software (Qiagen Inc.) or the clusterProfiler package in R for the KEGG pathways. Presentation of changes (log2 fold-change) in the expression of core genes in selected pathways as a heatmap.

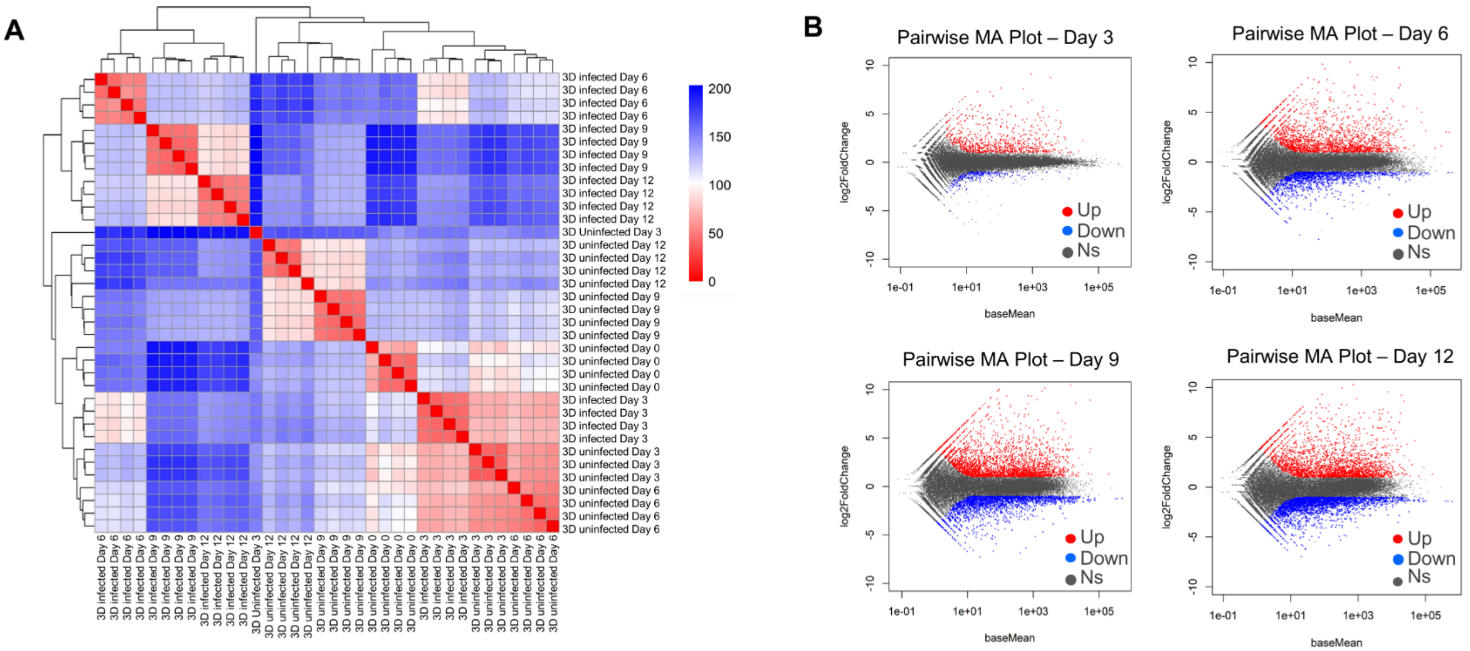


**S11 Fig. Additional RNA sequencing data analyses of the 3D cell culture samples.**

(**A**) The heatmap (clustergram) to visualize the hierarchical clustering of gene expression across 3D co-culture and uninfected control RNA-Seq samples on days 0, 3, 6, 9, and 12 post-3D cell culture. (**B**) The MA plots show the log2 fold-change (log2FC) versus the base mean expression for each gene in 3D co-cultures compared to 3D controls. Red dots show up-regulated genes with log2FC >1 and adjusted $p$-value < 0.05, blue dots indicate down-regulated genes with log2FC < −1 and adjusted $p$-value <0.05, and grey dots indicate genes with nonsignificant (ns) change in expression with adjusted $p$-value ≥ 0.05.

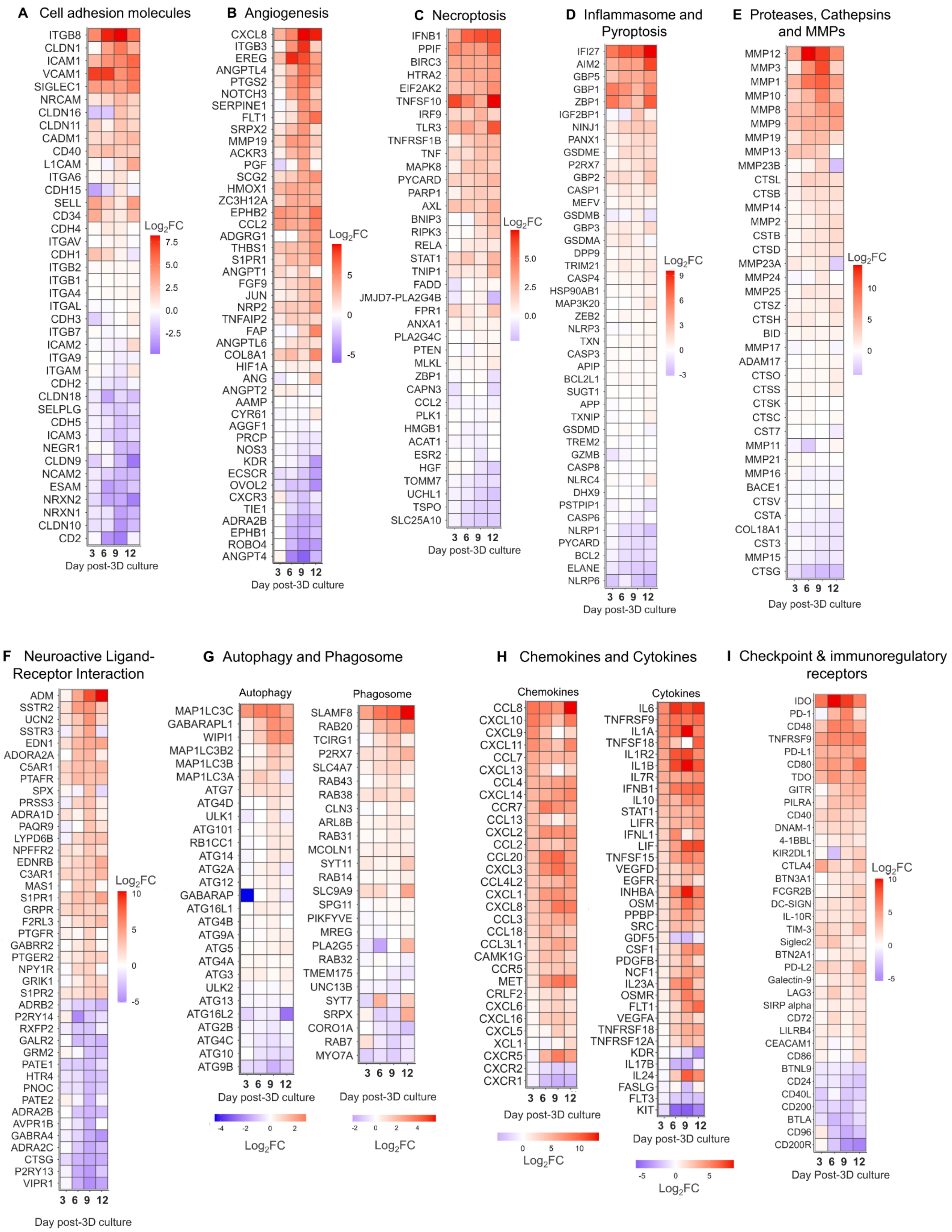

A Cell adhesion molecules
ITGB8
CLDN1
ICAM1
VCAM1
SIGLEC1
NRCAM
CLDN16
CLDN11
CADM1
CD40
L1CAM
ITGA6
CDH15
SELL
CD34
CDH4
ITGAV
CDH1
ITGB2
ITGB1
ITGA4
ITGAL
CDH3
ITGB7
ICAM2
ITGA9
ITGAM
CDH2
CLDN18
SELPLG
CDH5
ICAM3
NEGR1
CLDN9
NCAM2
ESAM
NRXN2
NRXN1
CLDN10
CD2
3 6 9 12
Day post-3D culture
Log2FC
7.5
5.0
2.5
0.0
-2.5
B Angiogenesis
CXCL8
ITGB3
EREG
ANGPTL4
PTGS2
NOTCH3
SERPINE1
FLT1
SRPX2
MMP19
ACKR3
PGF
SCG2
HMOX1
ZC3H12A
EPHB2
CCL2
ADGRG1
THBS1
S1PR1
ANGPT1
FGF9
JUN
NRP2
TNFAIP2
FAP
ANGPTL6
COL8A1
HIF1A
ANG
ANGPT2
AAMP
CYR61
AGGF1
PRCP
NOS3
KDR
ECSCR
OVOL2
CXCR3
TIE1
ADRA2B
EPHB1
ROBO4
ANGPT4
3 6 9 12
Day post-3D culture
Log2FC
5
0
-5
C Necroptosis
IFNB1
PPIF
BIRC3
HTRA2
EIF2AK2
TNFSF10
IRF9
TLR3
TNFRSF1B
TNF
MAPK8
PYCARD
PARP1
AXL
BNIP3
RIPK3
RELA
STAT1
TNIP1
FADD
JMJD7-PLA2G4B
FPR1
ANXA1
PLA2G4C
PTEN
MLKL
ZBP1
CAPN3
CCL2
PLK1
HMGB1
ACAT1
ESR2
HGF
TOMM7
UCHL1
TSPO
SLC25A10
3 6 9 12
Day post-3D culture
Log2FC
5.0
2.5
0.0
D Inflammasome and Pyroptosis
IFI27
AIM2
GBP5
GBP1
ZBP1
IGF2BP1
NINJ1
PANX1
GSDME
P2RX7
GBP2
CASP1
MEFV
GSDMB
GBP3
GSDMA
DPP9
TRIM21
CASP4
HSP90AB1
MAP3K20
ZEB2
NLRP3
TXN
CASP3
APIP
BCL2L1
SUGT1
APP
TXNIP
GSDMD
TREM2
GZMB
CASP8
NLRC4
DHX9
PSTPIP1
CASP6
NLRP1
PYCARD
BCL2
ELANE
NLRP6
3 6 9 12
Day post-3D culture
Log2FC
9
6
3
0
-3
E Proteases, Cathepsins and MMPs
MMP12
MMP3
MMP1
MMP10
MMP8
MMP9
MMP19
MMP13
MMP23B
CTSL
CTSB
MMP14
MMP2
CSTB
CTSD
MMP23A
MMP24
MMP25
CTSZ
CTSH
BID
MMP17
ADAM17
CTSO
CTSS
CTSK
CTSC
CST7
MMP11
MMP21
MMP16
BACE1
CTSV
CSTA
COL18A1
CST3
MMP15
CTSG
3 6 9 12
Day post-3D culture
Log2FC
10
5
0
F Neuroactive Ligand-Receptor Interaction
ADM
SSTR2
UCN2
SSTR3
EDN1
ADORA2A
C5AR1
PTAFR
SPX
PRSS3
ADRA1D
PAQR9
LYPD6B
NPFFR2
EDNRB
C3AR1
MAS1
S1PR1
GRPR
F2RL3
PTGFR
GABRR2
PTGER2
NPY1R
GRIK1
S1PR2
ADRB2
P2RY14
RXFP2
GALR2
GRM2
PATE1
HTR4
PNOC
PATE2
ADRA2B
AVPR1B
GABRA4
ADRA2C
CTSG
P2RY13
VIPR1
3 6 9 12
Day post-3D culture
Log2FC
10
5
0
-5
G Autophagy and Phagosome
Autophagy
MAP1LC3C
GABARAPL1
WIPI1
MAP1LC3B2
MAP1LC3B
MAP1LC3A
ATG7
ATG4D
ULK1
ATG101
RB1CC1
ATG14
ATG2A
ATG12
GABARAP
ATG16L1
ATG4B
ATG9A
ATG5
ATG4A
ATG3
ULK2
ATG13
ATG16L2
ATG2B
ATG4C
ATG10
ATG9B
3 6 9 12
Day post-3D culture
-4 -2 0 2
Log2FC
Phagosome
SLAMF8
RAB20
TCIRG1
P2RX7
SLC4A7
RAB43
RAB38
CLN3
ARL8B
RAB31
MCOLN1
SYT11
RAB14
SLC9A9
SPG11
PIKFYVE
MREG
PLA2G5
RAB32
TMEM175
UNC13B
SYT7
SRPX
CORO1A
RAB7
MYO7A
3 6 9 12
Day post-3D culture
-2 0 2 4
Log2FC
H Chemokines and Cytokines
Chemokines
CCL8
CXCL10
CXCL9
CXCL11
CCL7
CXCL13
CCL4
CXCL14
CCR7
CCL13
CXCL2
CCL2
CCL20
CXCL3
CCL4L2
CXCL1
CXCL8
CCL3
CCL18
CCL3L1
CAMK1G
CCR5
MET
CRLF2
CXCL6
CXCL16
CXCL5
XCL1
CXCR5
CXCR2
CXCR1
3 6 9 12
Day post-3D culture
0 5 10
Log2FC
Cytokines
IL6
TNFRSF9
IL1A
TNFSF18
IL1R2
IL1B
IL7R
IFNB1
IL10
STAT1
LIFR
IFNL1
LIF
TNFSF15
VEGFD
EGFR
INHBA
OSM
PPBP
SRC
GDF5
CSF1
PDGFB
NCF1
IL23A
OSMR
FLT1
VEGFA
TNFRSF18
TNFRSF12A
KDR
IL17B
IL24
FASLG
FLT3
KIT
3 6 9 12
Day post-3D culture
-5 0 5
Log2FC
I Checkpoint & immunoregulatory receptors
IDO
PD-1
CD48
TNFRSF9
PD-L1
CD80
TDO
GITR
PILRA
CD40
DNAM-1
4-1BBL
KIR2DL1
CTLA4
BTN3A1
FCGR2B
DC-SIGN
IL-10R
TIM-3
Siglec2
BTN2A1
PD-L2
Galectin-9
LAG3
SIRP alpha
CD72
LILRB4
CEACAM1
CD86
BTNL9
CD24
CD40L
CD200
BTLA
CD96
CD200R
3 6 9 12
Day Post-3D culture
Log2FC
10
5
0
-5

**S12 Fig. Differentially expressed genes in the selected enriched pathways.**

The log2 fold-change in the expression of core genes involved in the pathway is plotted as a heatmap using the ggplot2 package in the R programming environment. The horizontal four blocks denote the gene expression levels on days 3, 6, 9, and 12 post-3D cell culture. The block color indicates $log_2$ fold-change ($log_2FC$); red indicates upregulation, and blue denotes downregulation. Selected pathways shown include (**A**) cell adhesion molecules, (**B**) angiogenesis, (**C**) necroptosis, (**D**) inflammasome and pyroptosis, (**E**) proteases, cathepsins, and matrix metalloproteinases (MMPs), (**F**) neuroactive ligand-receptor interactions, (**G**) autophagy and phagosome, (**H**) chemokine and cytokine pathway, and (**I**) checkpoint and immunoregulatory receptors.

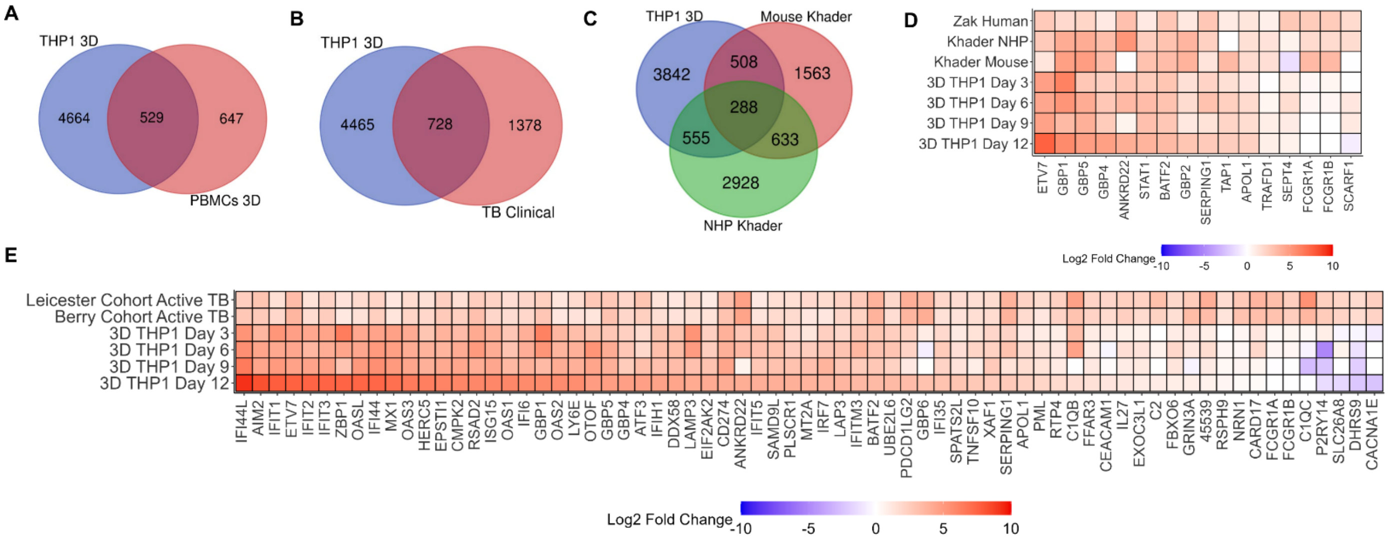


**S13 Fig. Transcriptomic comparison of the 3D tuberculoma model with published 3D human *in-vitro* granuloma model and human- or animal-derived TB tissue specimens.**

(**A**) The Venn diagram of shared DEGs between the 3D tuberculoma model of human THP-1 monocytes and *Mm* 'M' (day 9) and the published 3D *in-vitro* granuloma model of *Mtb*-infected human PBMCs and collagen ECM, microencapsulated in polymer particles, described by Elkington and colleagues. The published study isolated control uninfected and bioluminescent *Mtb* H37Rv-infected human PBMC 3D cell culture samples after microsphere decapsulation and cell lysis. (**B**) The Venn diagram of shared DEGs between the 3D tuberculoma model (day 9) and the human lymph node TB specimens described by the Elkington group. Control and mediastinal or neck lymph node TB biopsy clinical samples were used, and TB lymph node samples were culture-positive for drug-sensitive *Mtb* and exhibited caseating granulomas. (**C**) The Venn diagram shows shared DEGs between the 3D tuberculoma model (day 9) and *Mtb*-infected mouse and non-human primate lung specimens with granulomas described by Khader and colleagues. *Mtb* HN878 strain-infected DO mice and *Mtb* CDC1551-infected Indian rhesus macaques were classified as progressors in the study. (**D**) A heatmap displays the log2 fold-change in selected 16 signature genes that predict the onset of TB disease in humans, non-human primates, and mice, compared with the 3D tuberculoma model at four different time points post-co-culture. (**E**) A heatmap displays the $log_2$ fold-change in selected 70 signature genes upregulated in active TB patients relative to individuals with latent TB infection (from the UK and South African cohorts described in the study by O'Garra and colleagues) and compared to the 3D tuberculoma model at four different time points. See **S9 Table** for additional description and identification of specimens in published studies.

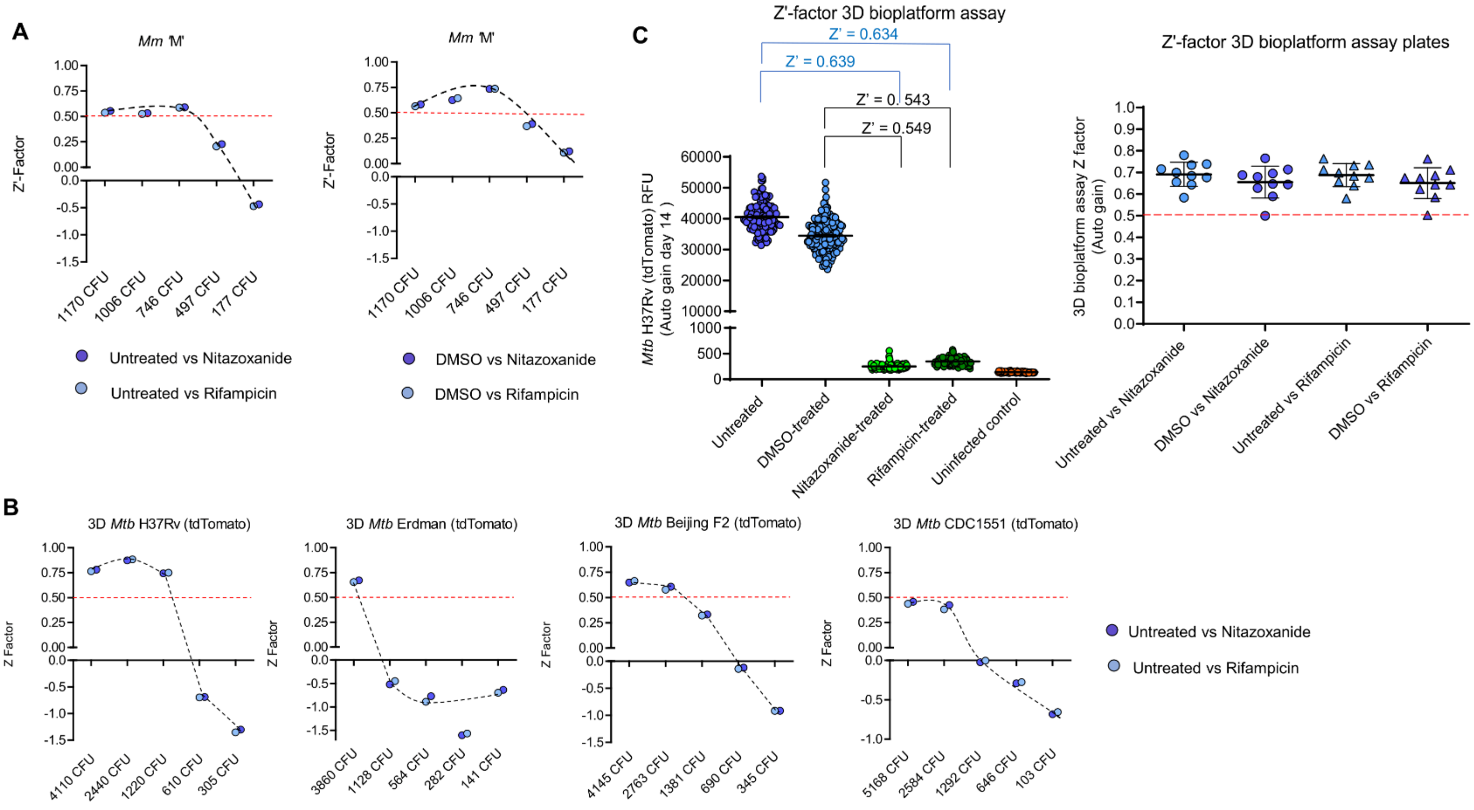


**S14 Fig. Chemical compound screening assay using 3D tuberculoma bioplatform exhibits excellent quality and performance.**

(**A**) The impact of *Mm* 'M' (tdTomato) MOI on the screening assay quality in the 3Dtuberculoma bioplatform of THP-1 as assessed by the Z'-factor statistic. The Z'-factor describes how well separated the positive and negative controls are and indicates the probability of false positives or negatives. The investigation was conducted using assay controls without the intervention of test compounds in a 96-well plate format. One infection dose per plate (Corning 3D) and five doses (MOI range 0.001 to 0.012) were investigated. Nitazoxanide (20 µM/well) and rifampicin (1 µg/ml) were used as positive controls for bacterial reduction, while DMSO and medium only (no drug or untreated) served as negative controls in the assay. FI was measured on day 12, and the data shown are from one of the two experiments performed (n = 12 tuberculomas/positive control and 12–24 tuberculomas/negative control per assay plate). The dashed line denotes a cutoff at a Z'-factor of 0.5. The Z'-factor value of 1.0 indicates an ideal assay, values between 0.5 and 1.0 indicate an excellent assay, and values between 0 and 0.5 indicate an acceptable assay. Values below 0 indicate that the assay conditions have not been optimized, and the assay is unlikely to generate valuable data. The optimal MOI range identified is 0.007 to 0.012 for the assay using *Mm* 'M' to exhibit an excellent Z'-factor and quality. (**B**) Determining the optimal MOI of *Mtb* (tdTomato) strains in the THP-1 bioplatform for an excellent screening assay quality. FI measurements on day 14 were considered for slow-growing *Mtb* strains. The optimal MOI range identified for *Mtb* strains H37Rv, Erdman, and Beijing is 0.025 to 0.05. Excellent assay quality could not be achieved for the *Mtb* CDC1551 strain, even with the higher MOI of 0.05 tested. Increased *Mtb*-induced THP-1 cell death was observed over 14 days in an assay investigating the

MOI of 0.1 for virulent *Mtb* strains; hence, the MOI doses >0.05 were not considered. (**C**) The performance and quality of the screening assay using 3D tuberculomas of THP-1 cells and *Mtb* H37Rv (tdTomato) in several 96-well plates. The assay was performed in 10 plates using an optimal MOI of 0.025, and FI was measured on day 14. Filled circles represent fluorescence in individual 3D tuberculomas or uninfected spheroids. RFU, relative fluorescent units. n = 126 untreated, 180 DMSO-treated, and 120 nitazoxanide or rifampicin-treated tuberculomas. To determine background fluorescence, 54 uninfected spheroids were included in the analysis. The assay demonstrated excellent quality and performance (Z'-factor >0.5). The performance in the individual plates was also assessed, and the Z' factor ≥0.5 in all ten plates further confirms the suitability of the assay for HTS applications.

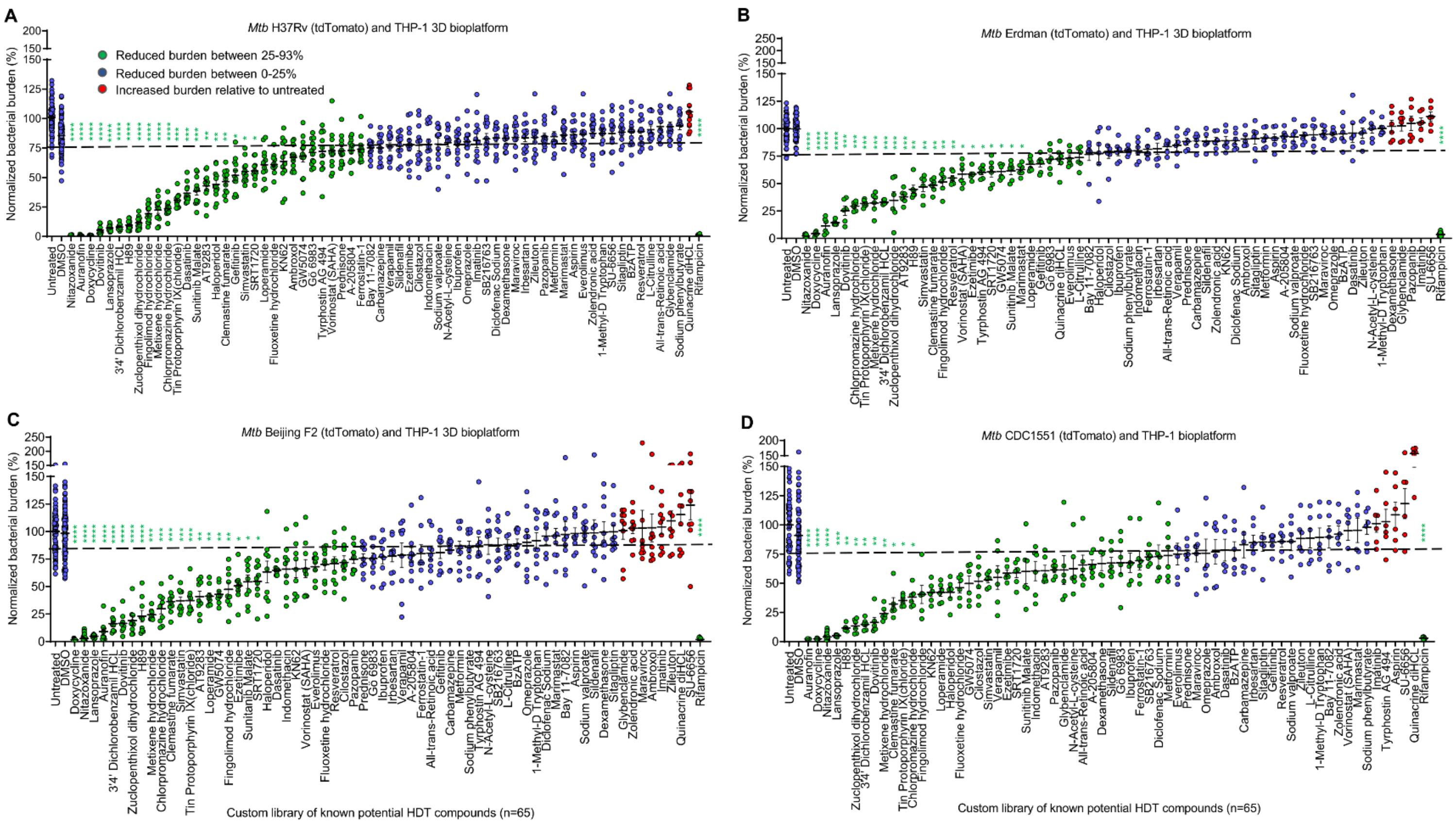


**S15 Fig. The HDT compound screens for the inhibition of mycobacterial burdens in the 3D tuberculoma bioplatform of THP-1 cells infected with four different *Mtb* strains.**
(**A**–**D**) Results of the HDT compound screens at 20 μM for inhibition of mycobacterial burdens in 3D tuberculoma bioplatform of THP-1 cells infected individually with (**A**) *Mtb* H37Rv, (**B**) *Mtb* Erdman, (**C**) *Mtb* Beijing F2, or (**D**) *Mtb* CDC 1551 constitutively expressing tdTomato, presented as a normalized bacterial burden (%) relative to untreated controls. Rifampicin was used as a positive control for reducing the bacterial burden. Tuberculomas were treated on day 6, and FI was measured on day 14 (day 8 post-treatment). Filled circles represent bacterial burden in individual 3D tuberculomas, and the data are from four (**A**), three (**C**), and two (**B** and **D**) independent experiments performed per strain (5 plates/experiment). Each plate contained tuberculomas treated with no drug (untreated, n = 6), DMSO (n = 6), rifampicin (n = 3–6), and test compounds (n = 3/compound). The dashed line represents a 25% reduction of bacterial burden. A horizontal line with error bars denotes mean ± SEM. *$p < 0.05$, **$p < 0.01$, ***$p < 0.001$, and ****$p < 0.0001$ compared to DMSO control by Kruskal-Wallis with Dunn's post-hoc test.

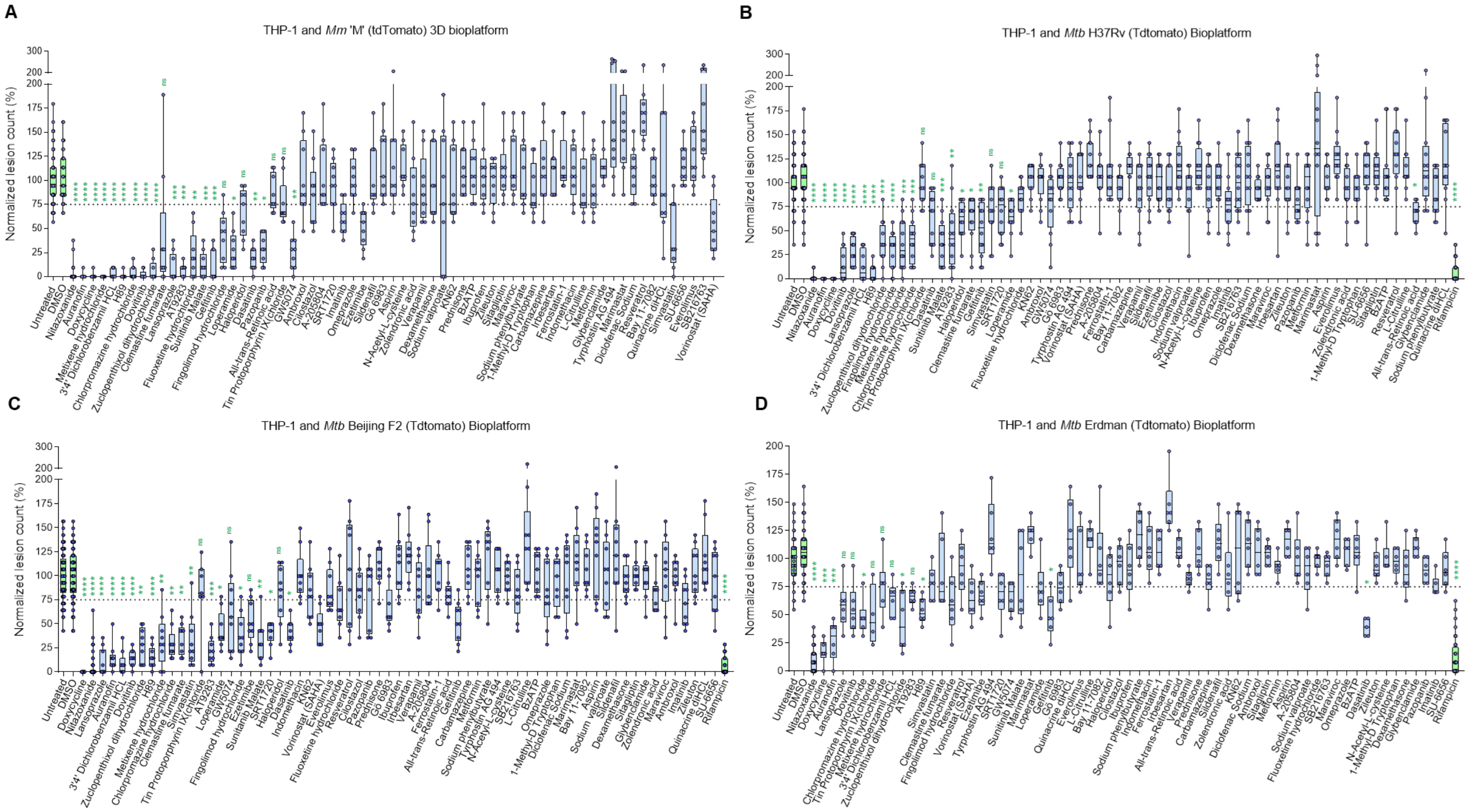


**S16 Fig. Inhibition of granuloma lesions following compound treatments in the 3D tuberculoma bioplatform of THP-1 cells infected with *Mm* or *Mtb* strains.**

(**A**–**D**) Inhibition of granuloma lesions following HDT compound treatments at 20 µM in 3D tuberculoma bioplatform of THP-1 cells infected individually with *Mm* 'M' (**A**), *Mtb* H37Rv (**B**), *Mtb* Beijing F2 (**C**), or *Mtb* Erdman (**D**) expressing tdTomato**.** Data are granuloma numbers counted in 3D co-cultures and expressed as normalized lesion counts (%) relative to untreated controls. To obtain granuloma lesion numbers, Z-projected images of 3D spheroids were subjected to cellular analysis and lesion counts using Gen5 software, followed by manual lesion counts by three blinded readers for quality control. The average of the three readers' counts was considered. The data are displayed as box plots with whiskers (minimum to maximum), showing all data points (replicates) as circles. The dotted line indicates a 25% inhibition of granuloma lesions. The data are from four (**B**), three (**A** and **C**), and two (**D**) independent experiments performed per strain (described in **Figs 5** and **S15**). *$p < 0.05$, **$p < 0.01$, ***$p < 0.001$, and ****$p < 0.0001$ compared to DMSO control by Kruskal-Wallis with Dunn's post-hoc test.

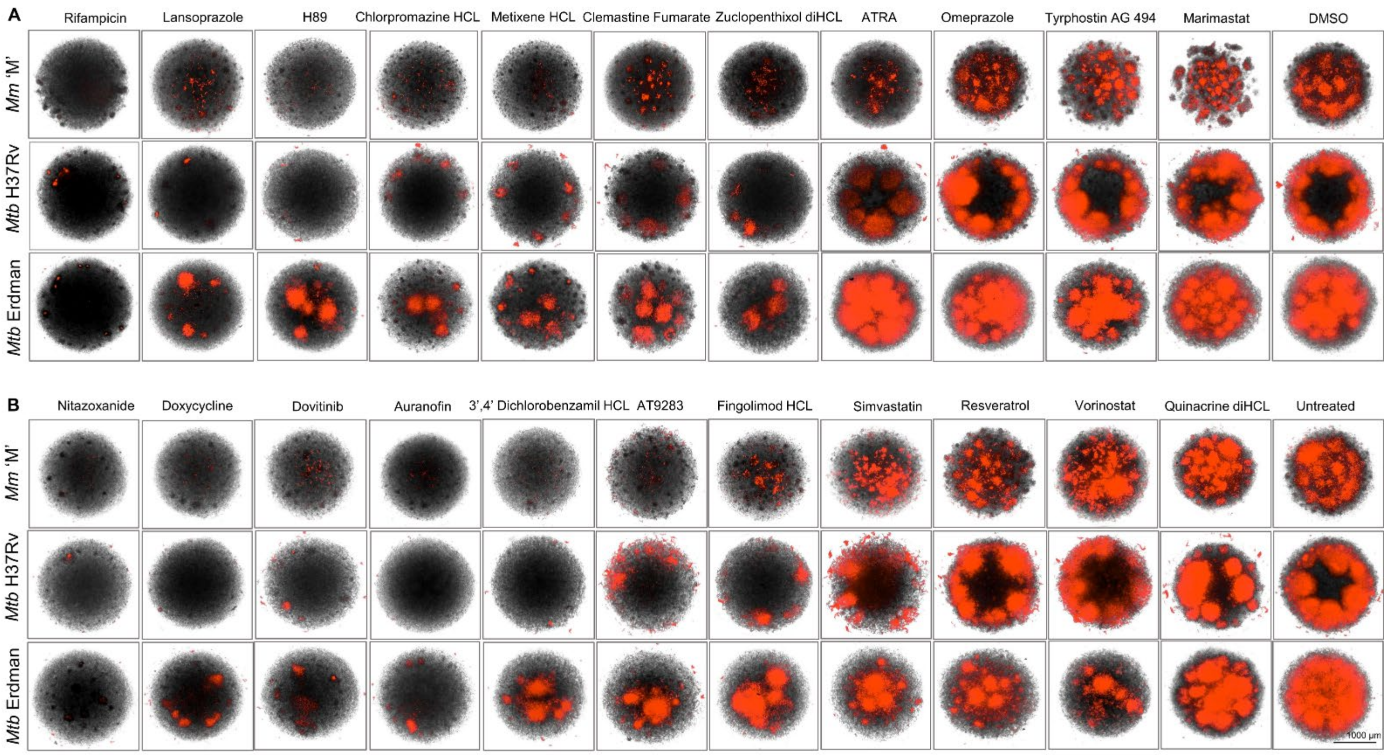


**S17 Fig. Representative images of inhibition of granuloma lesions following HDT compound treatments in 3D tuberculoma bioplatform.**

Images of 3D tuberculomas following treatment with (**A**) chemical compounds that did not decrease and (**B**) compounds that decreased cell viability by >50% at 20 μM in 3D cell cultures (For details of cytotoxicity results, refer to **S20 Fig**). Images represent 3D tuberculomas of THP-1 cells infected individually with *Mm* or *Mtb* strains and treated with HDT compounds on day 6 post-co-culture and captured on day 6 (*Mm* 3D) or between days 6 and 8 (*Mtb* 3D) post-treatment, from the experiments and replicates described in **Figs 5**, **S15,** and **S16**.

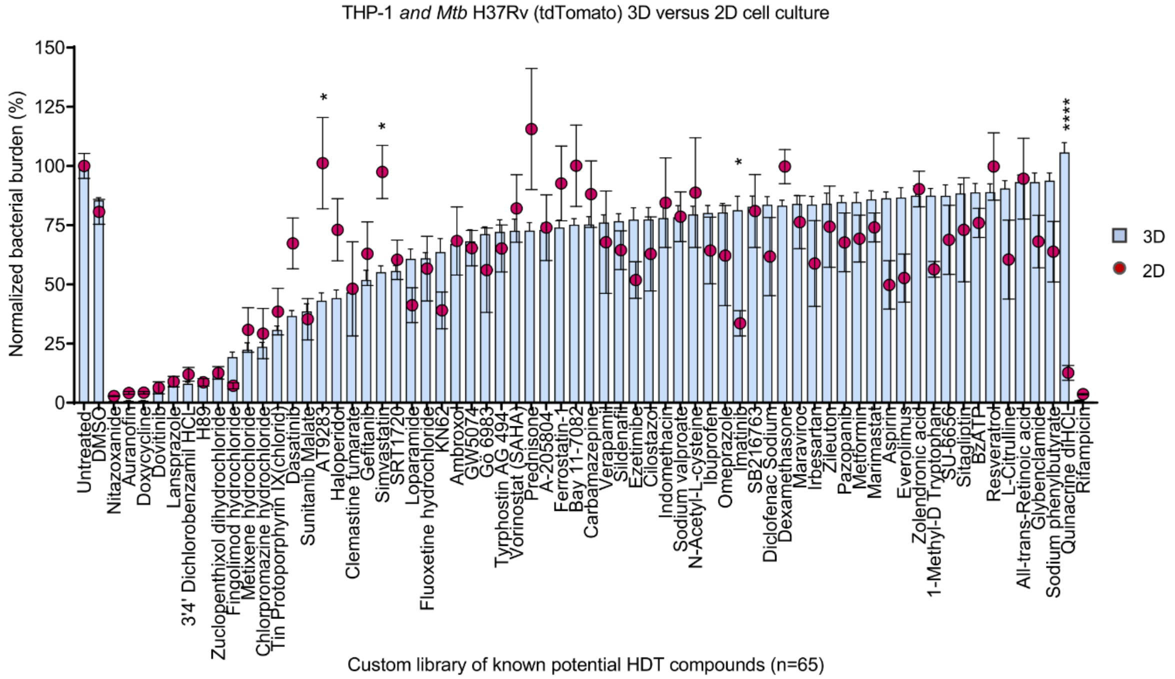


**S18 Fig. Effectiveness of chemical compounds on *Mtb* within 2D cell cultures compared to 3D tuberculoma bioplatform.**

**(A)** The reduction of bacterial burdens by 65 potential HDT compounds in 2D cell cultures compared to 3D tuberculomas of THP-1 cells and *Mtb* H37Rv (tdTomato) in 96-well 2D and 3D Corning ULA cell culture plates. Results are expressed as a normalized bacterial burden (%) relative to untreated controls. The data are mean ± SEM from two 2D cell culture experiments (n = 6–9 culture wells/compound) and four 3D cell culture experiments (n = 12 tuberculomas/compound). *$p$ < 0.05 and ****$p$ < 0.0001 comparing bacterial burdens in 2D cultures and 3D tuberculomas using Kruskal-Wallis with Dunn's post-hoc test of multiple comparisons of selected pairs.

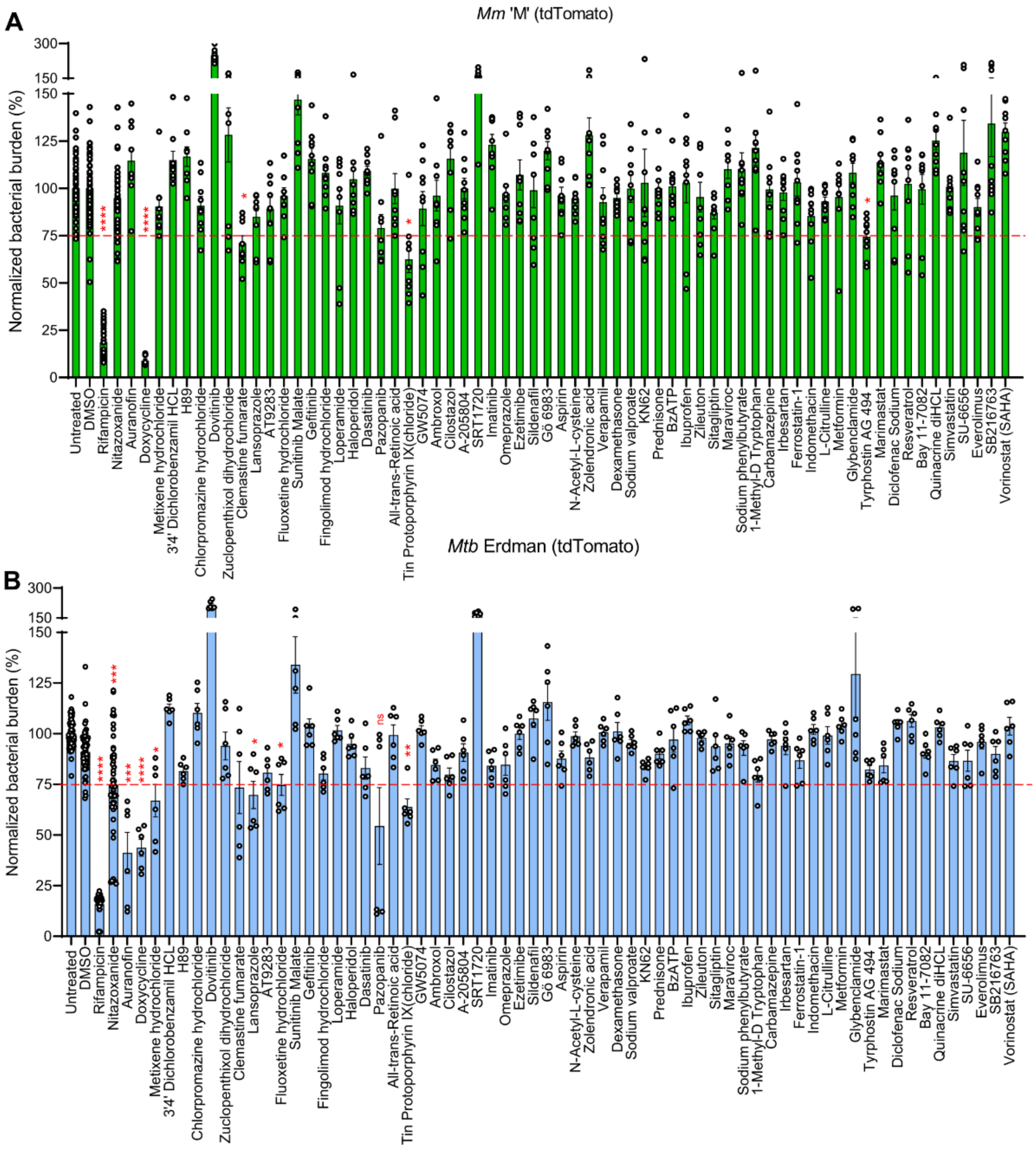


**S19 Fig. Compound screens in the axenic 3D broth cultures identify compounds with direct pathogen-targeting antimycobacterial effects in the 3D milieu.**

(**A**–**B**) Custom library compound screens at 20 µM in 3D broth microcultures of (**A**) *Mm* ‘M’ (tdTomato) or (**B**) *Mtb* H37Rv (tdTomato) in a final volume of 250 µl of Middlebrook 7H9 broth with hygromycin, incubated at 30 and 37 °C, respectively. Data are bacterial fluorescence measured and expressed as a normalized bacterial burden (%) relative to untreated controls at day 6 post-treatment. Generation of 3D broth microcultures for compound treatments involves plating 100 µl of mycobacterial culture (OD600, 0.05–0.06) in 7H9 broth containing hygromycin (50 µg/ml) in 3D culture plates (Corning), supplementing with 100 µl of fresh broth per microwell, and growing overnight at respective temperatures to form 3D clusters of bacteria. Data are from two experiments with each species. The bars represent the mean ± SEM, while the circles display bacterial burdens in replicate culture wells. A dashed line represents a 25% reduction of bacterial load. *$p < 0.05$, **$p < 0.01$, ***$p < 0.001$, and ****$p < 0.0001$ compared to DMSO controls using Kruskal-Wallis with Dunn’s post-test of multiple comparisons.

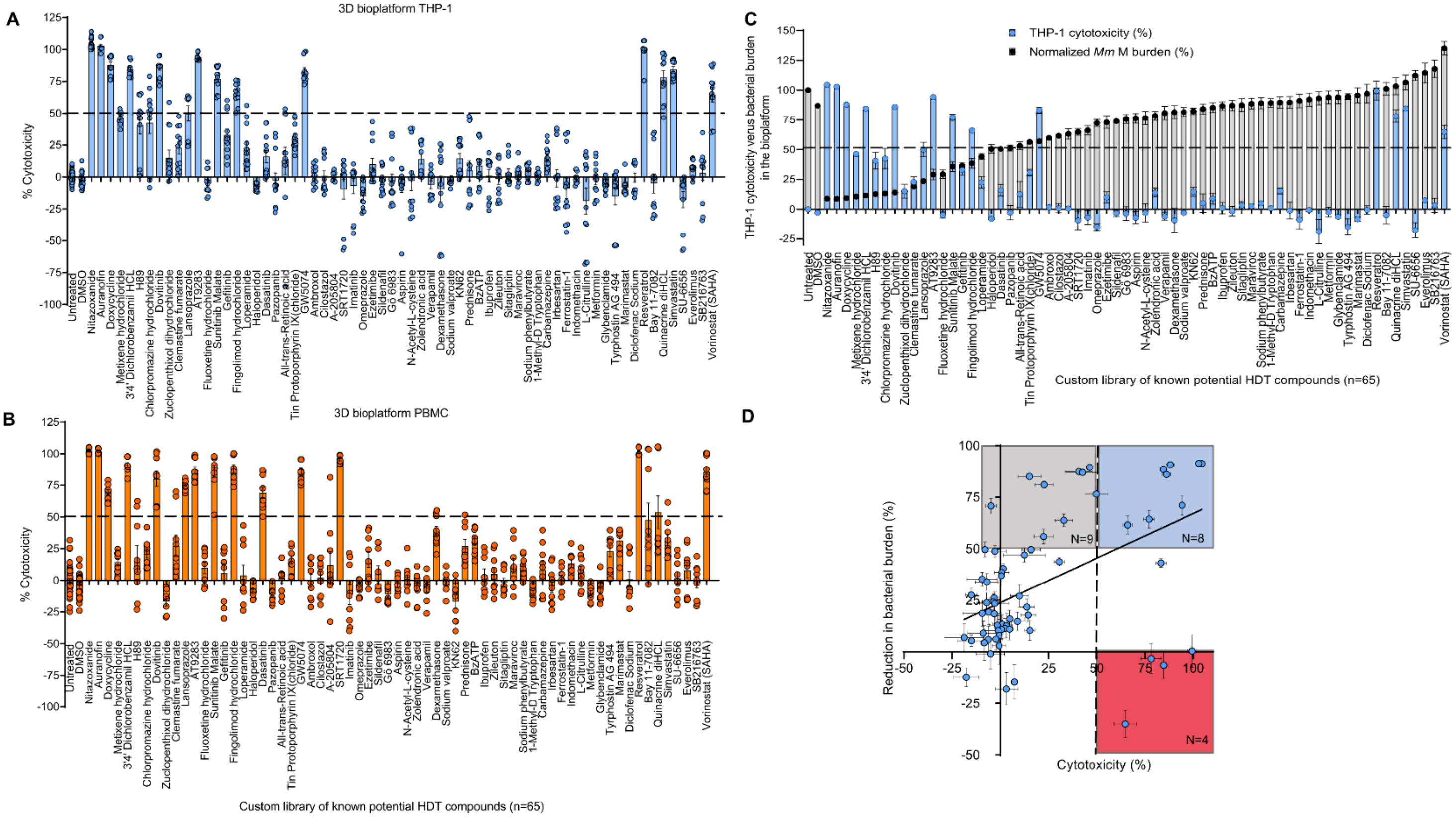


**S20 Fig. Identification of compounds causing host cell toxicity in 3D cell culture.**
(**A**–**B**) Cytotoxicity caused by HDT compounds at 20 µM in uninfected 3D cell cultures of (**A**) THP-1 monocytes or (**B**) human PBMCs ($CD14^+$ monocytes supplemented with $CD14^-$ subsets) following treatments on day six. Cytotoxicity was assessed using the CytoTox Glo assay on day 12 and expressed as a percentage of cytotoxicity compared to untreated controls. Data are of PBMCs isolated from 3 healthy donors and from three experiments performed for each cell type. Bars are mean ± SEM, while circles display cytotoxicity in replicate 3D cell cultures. A dashed line indicates a 50% decrease in cell viability. (**C**) Results of the cytotoxicity of compounds in the uninfected 3D cultures of THP-1 cells are plotted against the respective normalized bacterial burdens in the 3D tuberculoma bioplatform of THP-1 cells infected with *Mm* 'M' (tdTomato). Data are mean ± SEM. (**D**) Correlation between compound-induced THP-1 cytotoxicity and the reduction in bacterial burdens in the THP-1–*Mm* bioplatform. Simple linear regression $R^2 = 0.235$ and $p < 0.0001$.

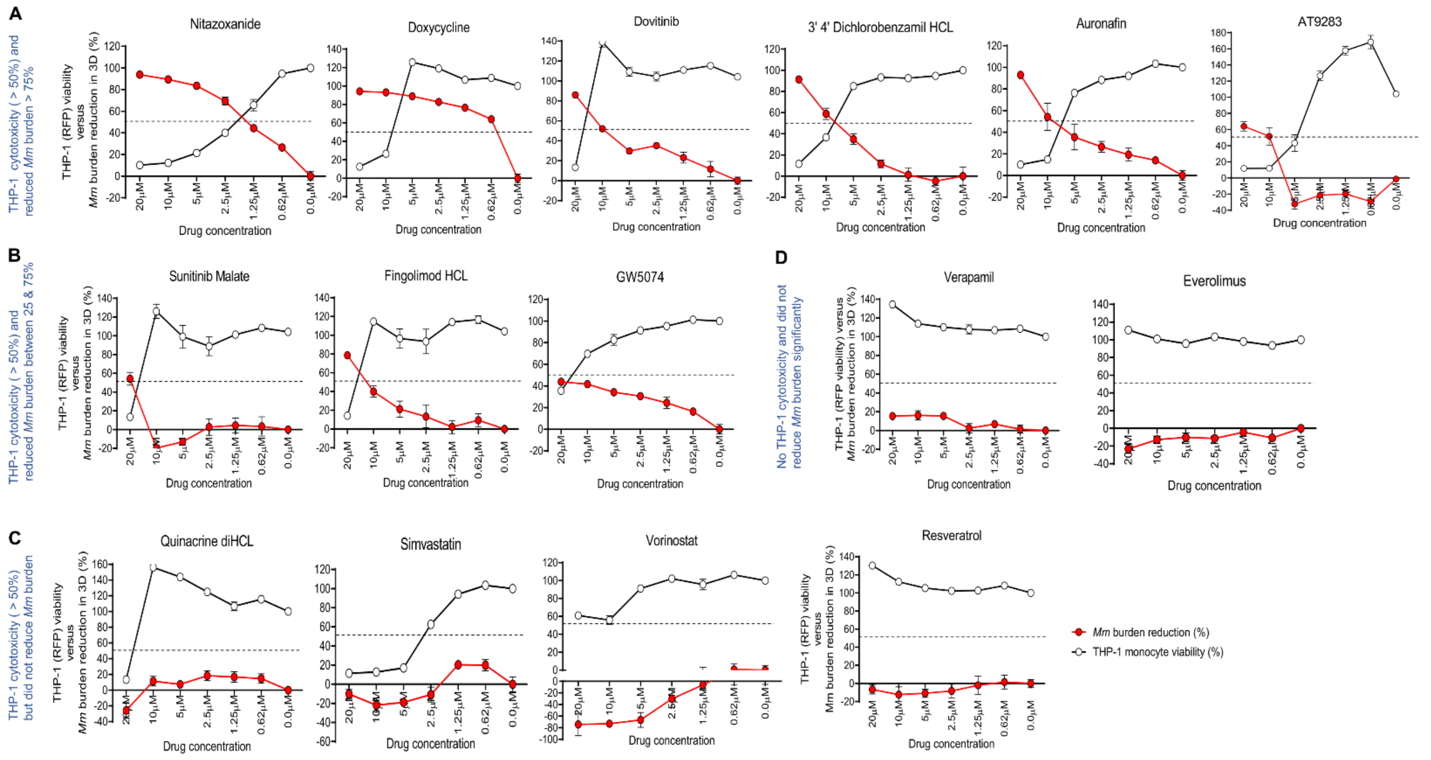


**S21 Fig. Determination of $CC_{50}$ and $EC_{50}$ values of cytotoxic compounds in 3D cell cultures.**

(**A**–**D**) The viability of THP-1 monocytes expressing RFP in 3D cultures treated on day 6 with a range of concentrations of each of the 13 HDT compounds identified as cytotoxic at 20 µM and measured on day 12 as fluorescence intensity (RFP). Cell viability is presented as a % of the no-drug controls. The cytotoxic concentration leading to a 50% reduction in cell viability ($CC_{50}$) is plotted against the % reduction in the bacterial burden in the THP-1–*Mm* 3D co-cultures, and the effective concentration 50 ($EC_{50}$) values (≥50% *Mm* growth inhibition) are determined. Each concentration was tested in duplicate cultures, and the data displayed are mean ± SEM from two experiments investigating cytotoxicity or bacterial burdens. A dashed line indicates a 50% reduction in THP-1-RFP viability and *Mm* burden. (**A**) Determining $CC_{50}$ and $EC_{50}$ for six cytotoxic 'top-hit' compounds. At nontoxic doses, auranofin provided <50% *Mm* growth inhibition, and AT9283 could not inhibit *Mm* growth. The $EC_{50}$ values could only be obtained for nitazoxanide, doxycycline, dovitinib, and 3'4' dichlorobenzamil HCl. (**B**) Determining $CC_{50}$ and $EC_{50}$ for three cytotoxic 'hit' compounds. At nontoxic doses, sunitinib malate could not inhibit *Mm* growth, and fingolimod HCl and GW5074 provided <50% *Mm* growth inhibition. (**C**) Investigating four cytotoxic 'non-hit' compounds. As expected, the 'non-hit' compounds, quinacrine diHCl, simvastatin, vorinostat, and resveratrol, did not inhibit *Mm* growth at nontoxic doses in 3D cell culture. Notably, resveratrol, which was found to be cytotoxic at 20 µM by CytoTox Glo assay, was consistently nontoxic in the THP-1-RFP assay. (**D**) Evaluating verapamil and everolimus as negative controls. Both compounds were found to be nontoxic and ineffective at all concentrations tested.

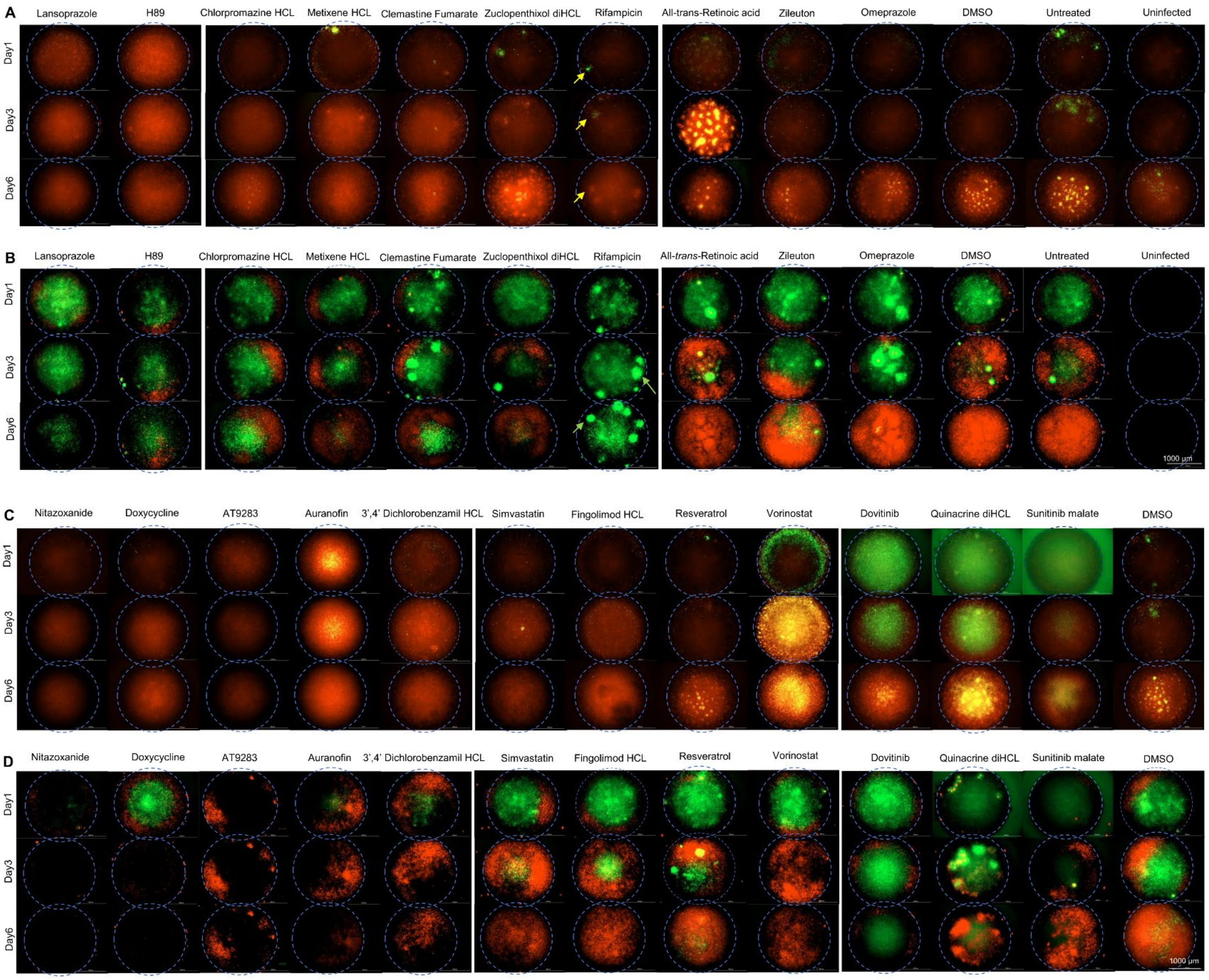


**S22 Fig. Kinetics of autophagy flux and inflammasome activation in 3D tuberculomas following treatment with HDT compounds.**

(**A**–**D**) Images of 3D tuberculomas of autophagy reporter cells and *Mtb* Erdman (WT) (**A** and **C**) or inflammasome reporter cells and *Mtb* Erdman (tdTomato) (**B** and **D**) treated with individual HDT compounds (20 μM), DMSO, or medium alone on day 6. (**A**) Autophagy flux in 3D tuberculomas following treatment with non-cytotoxic 'top-hit' or representative 'non-hit' compounds over six days. Tuberculomas treated with lansoprazole or H89 induced autophagy flux within 1-day post-treatment. Yellow arrows in the rifampicin-treated tuberculomas indicate a delay in autophagy induction in some cellular aggregates. (**B**) *Mtb* growth in 3D tuberculomas following treatment with non-cytotoxic 'top-hit' or 'non-hit' compounds and survival of ASC-GFP expressing reporter cells over six days. Treatments with lansoprazole, H89, or rifampicin effectively controlled *Mtb* (red) growth over six days. Green arrows in the rifampicin-treated granulomas indicate granulomatous cellular aggregates with strong ASC-GFP expression and inflammasome activation (green) and *Mtb* containment (yellow) in the center. (**C**) Autophagy flux in 3D tuberculomas following treatment with cytotoxic 'top-hit' compounds or compounds exhibiting *Mycobacterium* species-specific heterogeneity in growth inhibition. Tuberculomas treated with dovitinib ('top-hit'), quinacrine diHCl, or sunitinib malate exhibited autofluorescence in the GFP channel on day one post-treatment that decreased by day 3. Yet, quinacrine diHCl and sunitinib malate appeared to induce incomplete autophagy flux by day 6 (yellow-green fluorescence), although autofluorescence interfered with clear interpretation. (**D**) *Mtb* growth and survival of ASC-GFP expressing cells in 3D tuberculomas following treatment with cytotoxic 'top-hit' compounds or compounds exhibiting *Mycobacterium* species-specific heterogeneity in growth inhibition. Cytotoxic 'top-hit' compounds inhibited *Mtb* (red) growth despite the rapid loss of viability of ASC-GFP cells (green) and the known capability of intracellular *Mtb* to grow inside dead macrophages. (**A**–**D**) The images represent at least 3–6 tuberculomas treated per compound or control from one of the two experiments evaluating autophagy and one that investigated inflammasome activation.

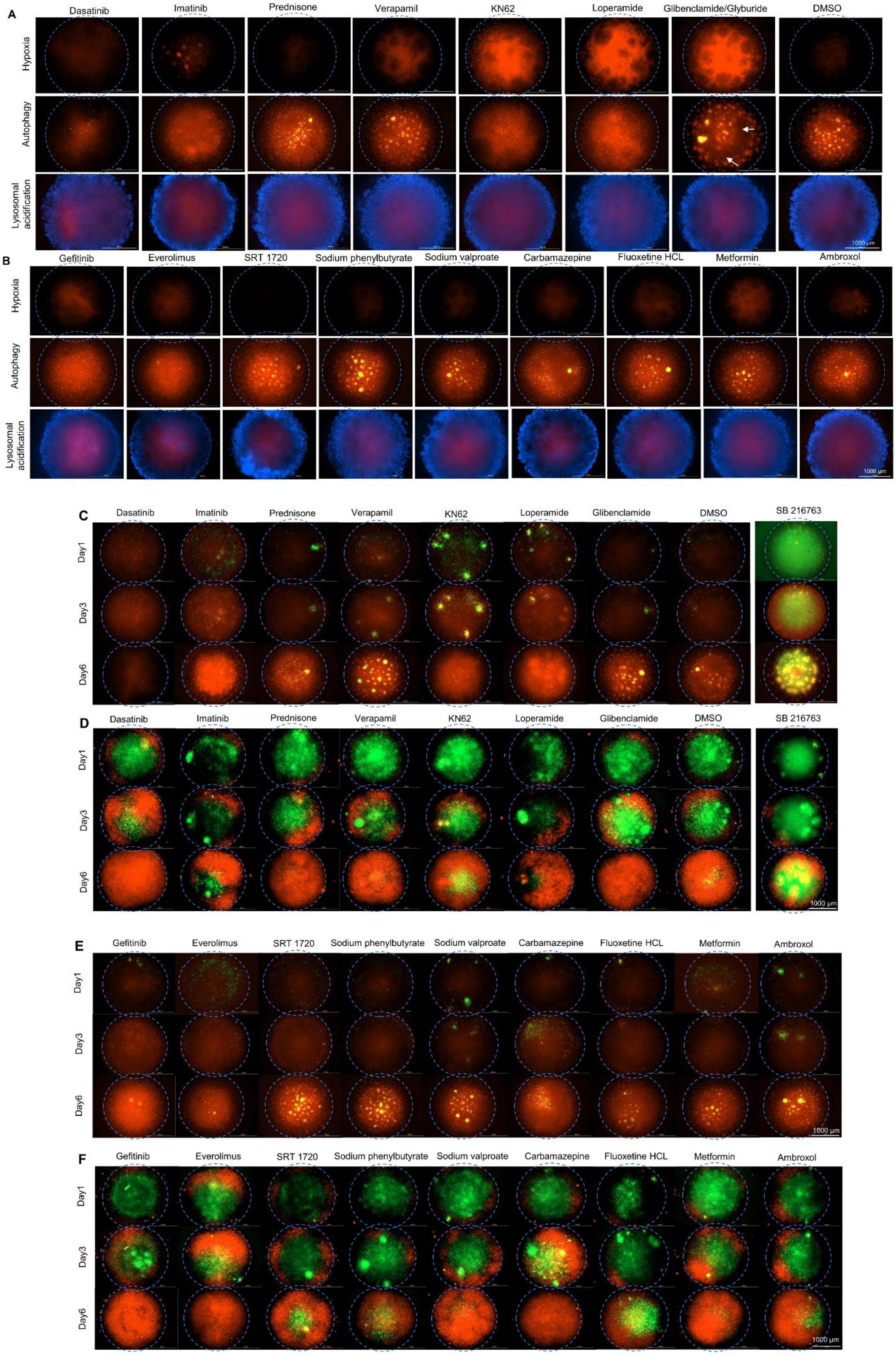
A
Dasatinib
Imatinib
Prednisone
Verapamil
KN62
Loperamide
Glibenclamide/Glyburide
DMSO
Hypoxia
Autophagy
Lysosomal acidification
1000 μm
B
Gefitinib
Everolimus
SRT 1720
Sodium phenylbutyrate
Sodium valproate
Carbamazepine
Fluoxetine HCL
Metformin
Ambroxol
Hypoxia
Autophagy
Lysosomal acidification
1000 μm
C
Dasatinib
Imatinib
Prednisone
Verapamil
KN62
Loperamide
Glibenclamide
DMSO
SB 216763
Day1
Day3
Day6
D
Dasatinib
Imatinib
Prednisone
Verapamil
KN62
Loperamide
Glibenclamide
DMSO
SB 216763
Day1
Day3
Day6
1000 μm
E
Gefitinib
Everolimus
SRT 1720
Sodium phenylbutyrate
Sodium valproate
Carbamazepine
Fluoxetine HCL
Metformin
Ambroxol
Day1
Day3
Day6
1000 μm
F
Gefitinib
Everolimus
SRT 1720
Sodium phenylbutyrate
Sodium valproate
Carbamazepine
Fluoxetine HCL
Metformin
Ambroxol
Day1
Day3
Day6
1000 μm

**S23 Fig. Association between any single innate immune mechanism induced by HDT compounds in 3D tuberculomas and inhibition of *Mtb* growth.**

Images of 3D tuberculomas of THP-1 cells and WT *Mtb* Erdman (**A** and **B**), autophagy reporter cells and WT *Mtb* Erdman (**A, B, C, and E**), and inflammasome reporter cells and *Mtb* Erdman (tdTomato) (**D** and **F**) treated with individual HDT compounds (20 μM) or DMSO on day 6 post-co-culture. Hypoxia and lysosomal acidification were investigated using the Hypoxia Red and Lyso-ID® Red reagents and Hoechst 33342 counterstain (blue) on day 12 in WT Erdman-infected THP-1 tuberculomas. (**A**) Induction of hypoxia, autophagy, and lysosomal acidification by non-cytotoxic compounds (n = 7). No direct association was found between compound-induced hypoxia, autophagy, or lysosomal acidification on day 6 post-treatment in 3D tuberculomas and the significant inhibition of *Mtb* Erdman growth or granuloma lesions compared to DMSO controls. (**B**) Induction of hypoxia, autophagy, and lysosomal acidification in 3D tuberculomas by known autophagy-inducing compounds (n = 9) that were identified previously using *Mtb*-infected 2D cell cultures. Besides gefitinib and everolimus, these compounds did not induce autophagy with autophagolysosome formation in the 3D milieu. (**C** and **E**) Autophagy flux induced in 3D tuberculomas over six days following treatment with (**C**) noncytotoxic compounds (n = 7) and (**E**) autophagy-inducing compounds (n = 9) identified in 2D systems. These compounds showed delayed (3–6 days) or incomplete autophagy flux. The noncytotoxic drug SB216763 showed autofluorescence in the GFP channel on day one post-treatment and appeared to induce incomplete autophagy by day 6. (**D** and **F**) ASC-GFP expression in inflammasome reporter cells and *Mtb* Erdman (tdTomato) growth over six days following treatment with (**C**) noncytotoxic compounds (n = 7) and (**E**) autophagy-inducing compounds (n = 9) identified in 2D systems. Autofluorescence of SB216763 interfered with the ASC-GFP fluorescence and reporter cell viability results. (**A**–**F**) The images represent at least 3–6 tuberculomas treated per compound or control from one of the two experiments evaluating hypoxia and autophagy, and one each that investigated lysosomal activation and inflammasome activation.

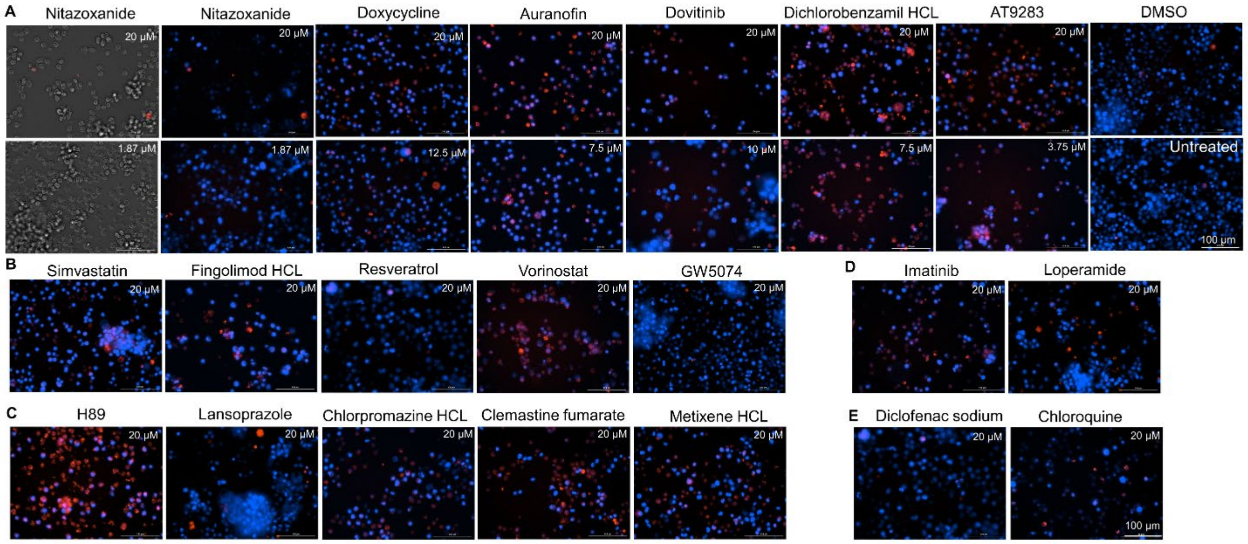


**S24 Fig. Autophagy modulation by selected HDT compounds in the *Mm*-infected THP-1 monocytes transfected with autophagy sensor LC3B-RFP.**

Modulation of autophagy in the *Mm* 'M' (WT)-infected 2D cell cultures of THP-1 monocytes, transfected with autophagy sensor LC3B-RFP, following treatments on day six post-infection with (**A**) cytotoxic 'top-hit' Cluster-1 compounds (n = 6). Representative fluorescence images captured on day 3 post-treatment of both cytotoxic concentration (20 µM) and one additional concentration ($CC_{50}$ or $EC_{50}$, if available for the compound) using Texas Red (LC3B, red) and DAPI (nucleus, blue) channels in a Cytation-5 are shown. Images of DMSO-treated and untreated cultures are presented as controls. Since Cluster-1 compounds cause a significant loss of THP-1 cell viability, corresponding brightfield images for nitazoxanide, a representative of the group, are shown to identify the total number of cells in the field. Representative images of modulation of autophagy by (**B**) cytotoxic compounds that did not significantly inhibit *Mm* burdens compared to controls (n = 5, all Cluster-5 compounds, except GW5074), (**C**) noncytotoxic 'top-hit' Cluster-2 compounds (n = 5), (**D**) compounds (n = 2) previously known to induce autophagy in *Mycobacterium*-infected macrophages, and (**E**) compounds (n = 2) thought to inhibit autophagy at 20 µM are also shown. The autophagy inhibitor diclofenac sodium impairs autophagy flux via oxidative stress and lysosomal dysfunction. Chloroquine has been demonstrated to inhibit autophagy by blocking autophagosome fusion with lysosomes and slowing down lysosomal acidification. Yet, chloroquine-induced lysosomal inhibition can inhibit mTORC1 and secondarily induce autophagy.

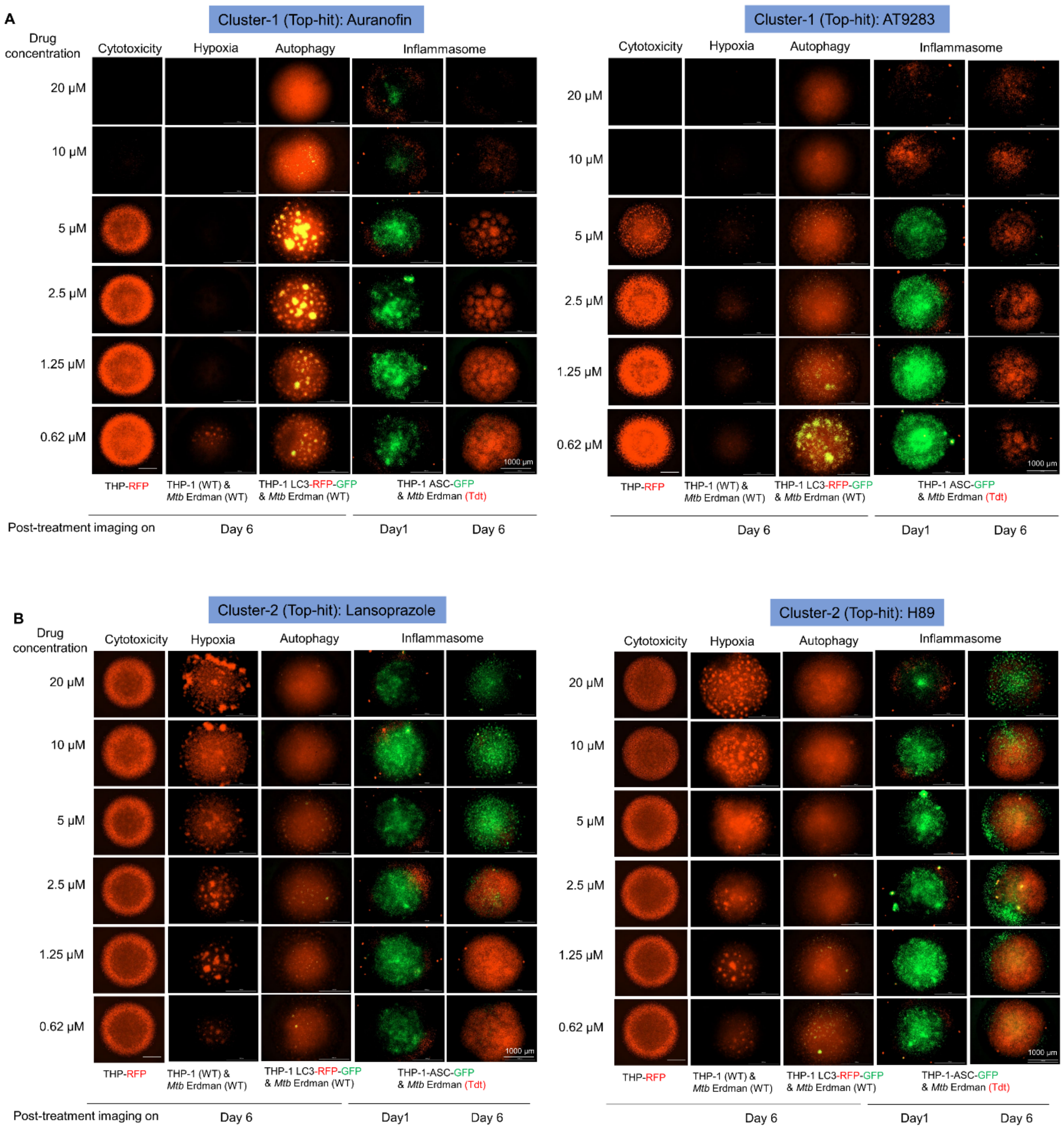


**S25 Fig. The effects of different concentrations of selected top-hit Cluster-1 and Cluster-2 compounds on the induction of innate defenses and *Mtb* burdens.**

Effects of six different concentrations between 20 and 0.62 µM of (**A**) auranofin and AT9283, and (**B**) lansoprazole and H89 on cytotoxicity, hypoxia, autophagy, inflammasome/pyroptosis induction, and *Mtb* burdens in 3D cell cultures. Treatments were performed on day 6 post-3D cell culture, and representative images were captured on day 1 or 6 post-treatment. The THP-1 monocytes, reporter cells, and *Mtb* Erdman WT or tdTomato used in 3D cell cultures are specified. Images from one of the two to three independent experiments, each with 2–3 replicates per dose of the compound, are shown.

### Supplementary table legends

**S1 Table. The levels of protein markers in the cell lysates and culture supernatants of infected and control 3D cell cultures supplemented with or without an extracellular matrix.**

The quantities of 1,000 protein markers were measured in cell lysate and culture supernatant samples collected on day 16 of 3D cultures using the Human Kiloplex Quantitative Proteomics Arrays. The table has three sheets, including two Excel data sheets for the cell lysate and culture supernatant proteins. The first sheet outlines the sample types and the methodology for calculating the fold-change in protein expression levels between infected and uninfected 3D samples, as well as the number of replicates used.

**S2 Table. Selected pathway proteins that are differentially expressed in infected versus control 3D cell cultures supplemented with or without an extracellular matrix.**

The levels of selected pathway protein markers, as measured in the cell lysates of 3D cultures generated with or without ECM using Human Kiloplex Quantitative Proteomics Arrays, are presented. The table has eight sheets, including 7 Excel data sheets for the selected protein lists in the seven pathways. The first sheet explains the sample types and the number of replicates tested. Each subsequent sheet displays a fold change in the amounts of selected protein members from a single pathway in the infected compared to uninfected control 3D cell culture samples.

**S3 Table. The quality of transcriptomics data.**

FastQC software was used to assess the quality of the raw FASTQ sequencing files. The number of total reads, GC%, quality scores of forward and reverse reads, and adapter content are presented. All forward reads showed a high-quality score (QC > 25). Some reverse reads showed decreased sequencing quality at the end of the reads (QC < 25). Only high-quality forward reads of each sample were used for alignment. After the STAR alignment, uniquely mapped reads, mapping percentage across the reference, and the mean coverage were quantified by Qualimap software.

**S4 Table. The differentially expressed genes in infected versus control 3D cell cultures at four different time points.**

The table is related to **Fig 5C**. It has two sheets: one for up-regulated genes and another for down-regulated genes. The long name and potential function or pathway are provided for each gene. The $\log_2$ fold change in expression (calculated for infected samples versus control samples) and adjusted *p*-values at four different time points are presented. The number of biological and technical replicates used is described in **Fig 5**.

**S5 Table. Selected pathway genes that are differentially expressed in infected versus control 3D cell cultures at four different time points.**

The table is related to **Fig 5** and **S12 Fig**. Differentially expressed genes within the selected pathway were identified for nine pathways. The table has ten sheets. The 'description tab' sheet describes the methodology of gene selection. The remaining nine sheets contain $\log_2$ fold-changes calculated as the ratio of infected/control samples and adjusted *p*-values for selected members from 9 different pathways. The biological and technical replicates used are described in Fig. 5.

**S6 Table. A custom-made library of known potential HDT compounds.**

The custom-made chemical compound library includes 65 potential HDT compounds purchased from the Small Molecule Screening Library Service, Aldrich Market Select, Millipore Sigma, USA. The table has two sheets. The first sheet provides general information about the chemical compounds, including their published activity in 2D cell culture, animal models, and humans. The second sheet provides additional information about the chemical names, molecular weights, formulas, and smiles.

**S7 Table. Summary of the therapeutic efficacy of chemical compounds in the 3D milieu.**

The therapeutic activity of 65 compounds regarding THP-1 and PBMC toxicity in 3D cell cultures, inhibition of bacterial burdens and granuloma lesions in 3D tuberculomas, and direct antibacterial activity in 3D axenic cultures are summarized. Average values for percent cytotoxicity, bacterial burden, and granuloma lesion number inhibition in the experiments performed are presented.

**S8 Table.  The key materials, reagents, and resources used in the study.**

The table lists the material or regent types, bacterial strains, and other resources used in the study, along with their identifiers and sources.

**S9 Table. Description of the 3D *in-vitro* granuloma and *Mtb*-infected animal and human samples from published studies used for comparative transcriptomic analysis.**

The table has six sheets. The first sheet describes the control and *Mtb*-infected samples from the human PBMC 3D cultures from the study published by Elkington and colleagues. The second sheet includes a description of the clinical samples from the control and TB lymph node biopsies in the same study. The third sheet describes the naïve and *Mtb*-infected DO mice and NHP lung samples from the study published by Khader and colleagues. The fourth sheet describes the blood samples from control and active TB patients in the UK cohort, as published in the study by O'Garra and colleagues. The fifth sheet includes the differentially expressed 70 signature genes in the blood of active TB patients compared to latent TB patients from the two cohorts described by O'Garra and colleagues. The sixth sheet describes the 16 signature genes that are differentially expressed between progressors and non-progressors in the South African adolescent cohort, which identifies immune correlates of TB disease across animal models and human TB, as reported in the published study by Khader and colleagues. These last two sheets describe the $\log_2$ fold-change and adjusted *p*-values for these signature genes in the *Mm* ‘M’-infected compared to control 3D THP-1 cell culture samples from the current study.

#### Supplementary references


1. Velu V, Titanji K, Zhu B, Husain S, Pladevega A, Lai L, et al. Enhancing SIV-specific immunity in vivo by PD-1 blockade. Nature. 2009;458(7235):206-10. Epub 20081210. doi: 10.1038/nature07662. PubMed PMID: 19078956; PubMed Central PMCID: PMCPMC2753387.
2. L. McInnes, J. Healy, J. Melville, Umap: Uniform manifold approximation and projection for dimension reduction. arXiv (2018) https://arxiv.org/abs/1802.03426 arXiv:1802.03426.
3. Bolger AM, Lohse M, Usadel B. Trimmomatic: a flexible trimmer for Illumina sequence data. Bioinformatics. 2014;30(15):2114-20. Epub 20140401. doi: 10.1093/bioinformatics/btu170. PubMed PMID: 24695404; PubMed Central PMCID: PMCPMC4103590.
4. Frankish A, Diekhans M, Jungreis I, Lagarde J, Loveland JE, Mudge JM, et al. Gencode 2021. Nucleic Acids Res. 2021;49(D1):D916-D23. doi: 10.1093/nar/gkaa1087. PubMed PMID: 33270111; PubMed Central PMCID: PMCPMC7778937.
5. Dobin A, Davis CA, Schlesinger F, Drenkow J, Zaleski C, Jha S, et al. STAR: ultrafast universal RNA-seq aligner. Bioinformatics. 2013;29(1):15-21. Epub 20121025. doi: 10.1093/bioinformatics/bts635. PubMed PMID: 23104886; PubMed Central PMCID: PMCPMC3530905.
6. Ewels P, Magnusson M, Lundin S, Kaller M. MultiQC: summarize analysis results for multiple tools and samples in a single report. Bioinformatics. 2016;32(19):3047-8. Epub 20160616. doi: 10.1093/bioinformatics/btw354. PubMed PMID: 27312411; PubMed Central PMCID: PMCPMC5039924.
7. Liao Y, Smyth GK, Shi W. The R package Rsubread is easier, faster, cheaper and better for alignment and quantification of RNA sequencing reads. Nucleic Acids Res. 2019;47(8):e47. doi: 10.1093/nar/gkz114. PubMed PMID: 30783653; PubMed Central PMCID: PMCPMC6486549.
8. Love MI, Huber W, Anders S. Moderated estimation of fold change and dispersion for RNA-seq data with DESeq2. Genome Biol. 2014;15(12):550. doi: 10.1186/s13059-014-0550-8. PubMed PMID: 25516281; PubMed Central PMCID: PMCPMC4302049.
9. Pisu D, Huang L, Grenier JK, Russell DG. Dual RNA-Seq of Mtb-Infected Macrophages In Vivo Reveals Ontologically Distinct Host-Pathogen Interactions. Cell Rep. 2020;30(2):335-50 e4. doi: 10.1016/j.celrep.2019.12.033. PubMed PMID: 31940480; PubMed Central PMCID: PMCPMC7032562.
10. Yu G, Wang LG, Han Y, He QY. clusterProfiler: an R package for comparing biological themes among gene clusters. OMICS. 2012;16(5):284-7. Epub 20120328. doi: 10.1089/omi.2011.0118. PubMed PMID: 22455463; PubMed Central PMCID: PMCPMC3339379.
11. Luo W, Brouwer C. Pathview: an R/Bioconductor package for pathway-based data integration and visualization. Bioinformatics. 2013;29(14):1830-1. Epub 20130604. doi: 10.1093/bioinformatics/btt285. PubMed PMID: 23740750; PubMed Central PMCID: PMCPMC3702256.

12. Ge SX, Jung D, Yao R. ShinyGO: a graphical gene-set enrichment tool for animals and plants. Bioinformatics. 2020;36(8):2628-9. doi: 10.1093/bioinformatics/btz931. PubMed PMID: 31882993; PubMed Central PMCID: PMCPMC7178415.
13. Reichmann MT, Tezera LB, Vallejo AF, Vukmirovic M, Xiao R, Reynolds J, et al. Integrated transcriptomic analysis of human tuberculosis granulomas and a biomimetic model identifies therapeutic targets. J Clin Invest. 2021;131(15). doi: 10.1172/JCI148136. PubMed PMID: 34128839; PubMed Central PMCID: PMCPMC8321576.
14. Ahmed M, Thirunavukkarasu S, Rosa BA, Thomas KA, Das S, Rangel-Moreno J, et al. Immune correlates of tuberculosis disease and risk translate across species. Sci Transl Med. 2020;12(528). doi: 10.1126/scitranslmed.aay0233. PubMed PMID: 31996462; PubMed Central PMCID: PMCPMC7354419.
15. Singhania A, Verma R, Graham CM, Lee J, Tran T, Richardson M, et al. A modular transcriptional signature identifies phenotypic heterogeneity of human tuberculosis infection. Nat Commun. 2018;9(1):2308. Epub 20180619. doi: 10.1038/s41467-018-04579-w. PubMed PMID: 29921861; PubMed Central PMCID: PMCPMC6008327.
16. McCaffrey EF, Delmastro AC, Fitzhugh I, Ranek JS, Douglas S, Peters JM, et al. The immunometabolic topography of tuberculosis granulomas governs cellular organization and bacterial control. bioRxiv. 2025. Epub 20250223. doi: 10.1101/2025.02.18.638923. PubMed PMID: 40027668; PubMed Central PMCID: PMCPMC11870603.

17. Kimura S, Noda T, Yoshimori T. Dissection of the autophagosome maturation process by a novel reporter protein, tandem fluorescent-tagged LC3. Autophagy. 2007;3(5):452-60. Epub 20070521. doi: 10.4161/auto.4451. PubMed PMID: 17534139.
18. Zhang JH, Chung TD, Oldenburg KR. A Simple Statistical Parameter for Use in Evaluation and Validation of High Throughput Screening Assays. J Biomol Screen. 1999;4(2):67-73. doi: 10.1177/108705719900400206. PubMed PMID: 10838414.
19. Malo N, Hanley JA, Cerquozzi S, Pelletier J, Nadon R. Statistical practice in high-throughput screening data analysis. Nat Biotechnol. 2006;24(2):167-75. doi: 10.1038/nbt1186. PubMed PMID: 16465162.

20. Sable SB, Kline A, Li W, Posey JE. Methodology for a freshly engineered or cryo-preserved 3D tuberculoma bioplatform for studying tuberculosis biology and high-content screening of therapeutics. bioRxiv. 2025.
